# Systematic Characterization of Optical Aberrations Reveals Cryo-FLM Localization Fidelity

**DOI:** 10.1101/2025.07.25.666845

**Authors:** Hongjia Li, Lauren Ann Metskas, Fang Huang

## Abstract

Cryo-correlative light and electron microscopy (cryo-CLEM) facilitates *in situ* imaging and structural analysis by combining the molecular specificity of fluorescence microscopy with the ultrastructural resolution of cryo-electron microscopy. By further combining single molecule localization with cryo-CLEM, molecular positions of individual emitters can be revealed in the context of the electron density map of a cell, providing unique insights to profound questions in cell biology and virology. However, cryogenic fluorescence light microscopy (cryo-FLM) suffers from severe and spatially heterogeneous optical aberrations that distort the point spread function, limiting the accuracy of molecular localizations as well as downstream cryo-transmission electron microscopy workflows. Here, we present a systematic and quantitative analysis of optical aberrations in a commercial cryo-FLM system, uncovering the sources of significant distortions such as system imperfections, refractive index mismatches, and sample-induced heterogeneities. These system and sample induced aberrations lead to localization errors up to 90 nm laterally and over 300 nm axially, challenging the feasibility of precise molecular positioning within the vitrified specimen. We demonstrate that these errors are partially mitigated by spatially matched or adaptive point spread function models pushing the error rate down to ten nanometers or less, offering practical guidance for aberration-aware cryo-FLM and cryo-CLEM strategies. Our findings highlight the necessity of accurate, *in situ* point spread function modeling to achieve nanometer-scale localization in cryo-FLM. The experimental pipeline developed in this work establishes a novel tool to assess optical performance in cryo-CLEM and cryogenic focused ion beam milling workflows as the field strives toward accurate and precise molecular localization.

## Introduction

Sub-tomogram averaging (STA) has enabled near-atomic resolution reconstructions of large, morphologically distinct macromolecular assemblies including HIV capsids, COP-II lattices, and ribosomes^1–4^. However, STA requires unambiguous particle identification at approximately 3 nm resolution, restricting its application to large macromolecules with distinct and recognizable features^5–7^. Compounding this limitation, the field of view (FOV) in cryo-electron tomography (cryo-ET) is limited to roughly 1-3 µm, making it challenging to target rare or sparsely distributed structures. Cryo-correlative light and electron microscopy (cryo-CLEM) addresses these limitations by labeling target proteins with fluorescent tags and projecting their spatial distribution onto the transmission electron microscopy (TEM) field^8^. In theory, this could enable molecular identification of structures otherwise indistinguishable in electron density maps, allowing STA to be used for a wider range of *in situ* targets. For specimens thicker than the roughly 300 nm limit for TEM, cryo-focused ion beam (FIB) milling is typically used to thin the sample, with cryo-fluorescence light microscopy (cryo-FLM) guiding the milling process to ensure retention of the target region^9,10^.

Central to the potential for using cryo-FLM to identify specific proteins in correlative workflows is its capability to localize fluorescently tagged molecules with nanometer precision, a goal not currently achieved in the field. Even small localization errors at this stage can propagate into substantial inaccuracies in the subsequent cryo-TEM interpretation. The fidelity of molecular localization is fundamentally determined by the retrieving accuracy of the point spread function (PSF), which represents the three-dimensional (3D) emission pattern of a single fluorophore after transmitting through the vitrified specimen and the downstream optical train^11–13^. To date, this response function is commonly modeled theoretically by using two-dimensional (2D) or 3D Gaussians^14–17^. However, under cryogenic conditions, fidelity of Gaussian PSFs can be severely compromised by the spatially heterogeneous optical aberrations, including system imperfections, refractive index mismatch between objective immersion and the vitrified specimen, and sample-induced heterogeneities^18–20^, representing a critical barrier limiting localization accuracy and compromises downstream cryo-electron microscopy workflows. This may be one cause of current cryo-CLEM precision limitations at the 10^−8^ m scale.

To date, optical aberrations in cryo-FLM systems remain poorly characterized. This is largely due to their complexity and unpredictability, coupled with severe illumination power limitations inherent to cryogenic fluorescence imaging^21–23^. The unique optical configuration, featuring a long-working-distance air objective positioned above a liquid-nitrogen-cooled immersion interface, a vitrified biological specimen, a complex support substrate, and heterogeneous refractive indices within the sample, introduces substantial and spatially variable aberrations^8,24,25^. These factors collectively pose significant challenges to accurately model the aberrated PSF and the subsequent aberration correction strategies.

To address these challenges, we developed a comprehensive experimental pipeline and an open-source MATLAB toolbox for analyzing and modeling aberrations in cryo-FLM. We found that aberrations under cryogenic conditions are significantly more complex and severe than those observed at room temperature (RT) on the same instrument. By isolating and characterizing system-, sample-, and FOV-induced aberrations, we show that these distortions are spatially variable and often unpredictable. Critically, we demonstrate that mismatched PSF models introduce substantial localization errors, particularly along the axial dimension, up to 300 nm. This error can be partially mitigated by using PSF models derived from spatially proximal regions, underscoring the importance of spatially matched or adaptive PSF representations. Our study establishes the first systematic and quantitative analysis pipeline of optical aberrations in cryo-FLM relevant to cryo-CLEM workflows, offering practical guidance on PSF modeling and aberration-aware localization strategies. We hope this development provides a critical step toward achieving nanometer-scale molecular mapping within complex cryogenic environments.

## Results

### Visualization and Characterization of PSF Aberrations in Cryo-FLM

The primary distinction between cryo-FLM and RT FLM lies in the complexity and severity of optical aberrations. Beyond the intrinsic system-specific aberrations common to both modalities, cryo-FLM is subject to pronounced aberrations arising from refractive index (RI) mismatches^26,27^. These stem from the use of an air objective positioned above a sample with a higher RI medium, as well as from the complex, heterogeneous refractive indices present within the vitrified specimen^28–30^. These effects are further exacerbated by the vitrification process itself and the subtle deformations of the TEM grid, such as warping or bending during plunge-freezing and cryo-transfer^31^ (**Fig. 1b, Supplementary Fig. 1**). In contrast, conventional RT FLM typically mitigates RI mismatch using index-matched immersion and mounting media, and sample-induced RI heterogeneity is comparatively minor^32^. To isolate system-induced aberrations and assess the specific contributions of cryogenic conditions, we performed RT imaging of the same fluorescent bead sample used in cryo-FLM, under air conditions on the same instrument, thereby minimizing the influence of sample itself and potential FOV-dependent system-induced aberrations. All experiments were conducted on a commercial widefield illumination system (see ***Methods*** for details).

**Fig. 1.**
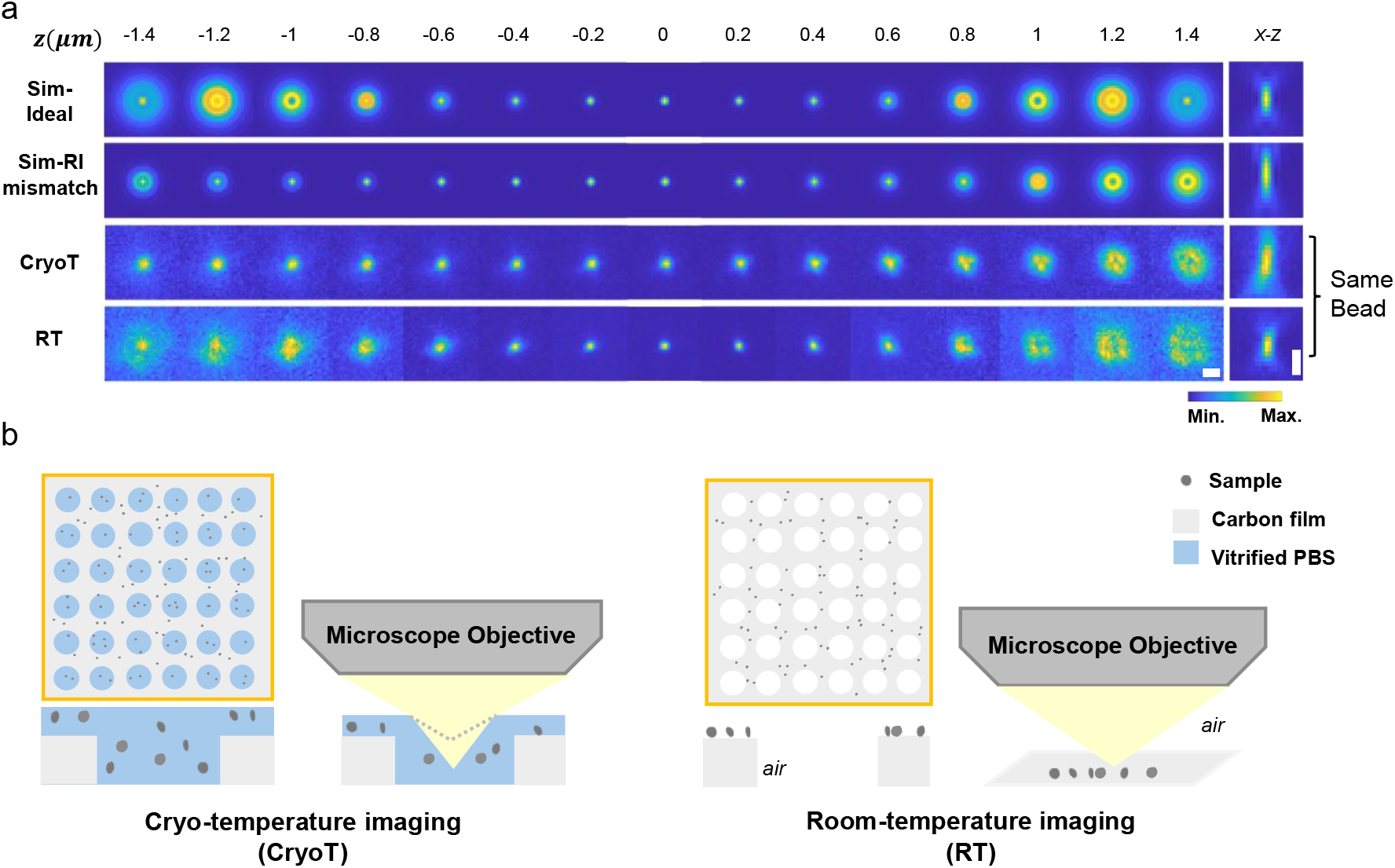
Visualization of simulated and experimentally measured PSF models. **a**. Simulated PSFs under ideal optical conditions (*Sim-Ideal*), with refractive index mismatch (*Sim-RI mismatch*), and experimentally measured PSFs under cryogenic conditions (*cryoT*) and at room temperature (*RT*), using the same fluorescent bead (200 nm TetraSpeck) across a focal range of −1.4 μm to +1.4 μm. **b**. Left: Bead distribution on a TEM grid and the light propagation in cryo-FLM imaging. Right: Bead distribution on a TEM grid and the light propagation in RT FLM imaging. Scale bars: 1 μm.

We found RI mismatch introduces pronounced spherical aberrations, resulting in an asymmetry in out-of-focus planes and axial elongation of the PSF profile (*Sim-RI mismatch* and *CryoT PSF* in **Fig. 1a**). Interestingly, experimental PSFs acquired at both cryogenic and room temperatures exhibit additional deformations, such as a tilt in the xz view, likely arising from system-induced factors. The PSF measured under cryogenic conditions displays even more severe distortions in both lateral shape and axial extent, despite being acquired from the same fluorescent bead. These additional aberrations likely originate from a combination of system-and sample-induced factors, underscoring the need for comprehensive aberration characterization.

### Room Temperature vs. Cryogenic Temperature Fluorescence Imaging

#### Direct Comparison of Aberrations under Cryogenic and Room Temperature Imaging

To investigate the specific impact of cryogenic conditions on optical aberrations, we designed an imaging strategy that enables direct comparison between cryogenic and room-temperature FLM using the same sample and hardware (**Fig. 2a; *Methods*; Supplementary Fig. 1**). Cryogenic conditions are known to introduce dominant and complex aberrations in cryo-FLM^33^. To isolate these effects, we imaged the same fluorescent bead sample under both vitrified (cryogenic temperature, cryoT) and dry (room-temperature, RT) conditions using an identical microscope setup and illumination source, and London Finder TEM grids that facilitate tracking specific beads^28,34^. This allows for a direct, spatially matched comparison across both conditions, eliminating confounding factors, such as bead-specific variability and FOV-dependent system differences. By maintaining fixed bead positions across imaging modalities, we are able to attribute observed differences primarily to cryogenic condition-induced aberrations, including refractive index mismatches and heterogeneous refractive indices of the vitrified sample. To minimize photobleaching, cryo-FLM imaging is performed first, followed by room-temperature imaging.

**Fig. 2.**
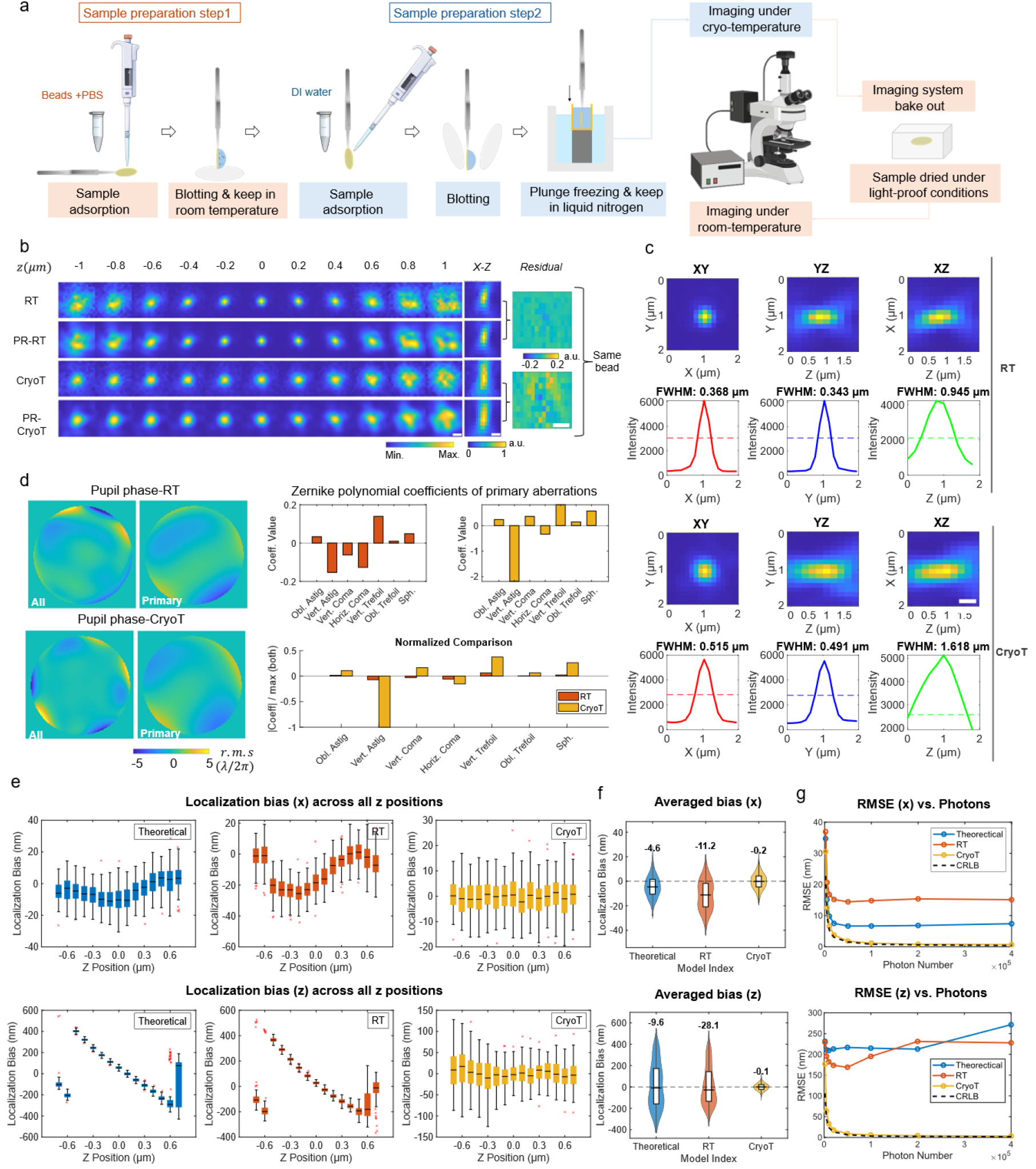
Room-temperature vs. cryo-temperature fluorescence microscopy imaging of the same bead. **a**. Schematic of the sample preparation and imaging workflow, showing sequential imaging of the same fluorescent bead under RT and cryoT conditions on the same instrument. **b**. Experimental and phase-retrieved PSF image stacks across axial positions from −1 µm to +1 µm for both RT and cryoT conditions. Corresponding xz views and residuals between experimental and reconstructed PSFs are also shown. **c**. Cross-sections of the PSFs in the xy, xz, and yz planes, along with intensity profiles and full width at half maximum (FWHM) measurements derived from the experimental PSFs in **b. d**. Pupil phase maps showing all aberration modes (5^th^-36^th^ Zernike terms) and primary aberrations (5^th^-11^th^ Zernike terms) for RT and cryoT PSF models, along with the corresponding Zernike coefficients representing the primary modes. **e**. Localization bias analysis across axial positions using three PSF models: (1) a theoretical pupil function, (2) the RT model, and (3) the cryoT model. The test dataset was generated using the cryoT PSF model, spanning axial positions from −0.7 µm to +0.7 µm in 100 nm increments (100 frames per z-position). Simulations were performed with a signal photon count of 1 × 10^4^ and a background photon count of 400. **f**. Lateral (x) and axial (z) bias distributions aggregated over all positions. **g**. Localization root-mean-square error (RMSE) versus the Cramér-Rao lower bound (CRLB) as a function of photon count. Scale bar: 500 nm.

#### Cryo-FLM Exhibits Stronger and More Complex Aberrations than RT-FLM

To investigate optical aberrations in the imaging system, we apply a maximum likelihood estimation (MLE)- based phase retrieval algorithm that is robust to high noise levels and capable of resolving complex wavefront distortions (see ***Methods***; **Extended Data Fig. 1; Supplementary Fig. 2**)^35,36^. This approach enables accurate reconstruction of PSF models from individual fluorescent bead image stacks, with residuals in the xz plane consistently below ± 0.2 a.u. (arbitrary units; the xz-plane is normalized to a range of 0-1, with the maximum intensity set to 1) (**Fig. 2b**). Compared to the RT PSF, the cryoT PSF appeared more diffuse near the focal plane and exhibited distinct patterns in the xy plane at defocused positions. In the xz view, it showed both tilt and pronounced axial elongation. Moreover, the cryoT PSF exhibits a 1.4- to 1.7-fold increase in full width at half maximum (FWHM), reflecting degraded resolution and reduced localization precision (**Fig. 2c**).

To quantify these distortions, we further decompose the aberrated wavefronts into primary modes using Zernike polynomials (**Fig. 2d**)^37,38^. Cryo-FLM reveals substantially stronger and distinct aberration profiles compared to RT imaging. Dominant modes include astigmatism (with an amplitude of 2.18 λ/2π, corresponding to the Zernike polynomial coefficients), coma (0.82 λ/2π), and spherical aberration (0.57 λ/2π). In contrast, RT FLM shows astigmatism, coma, and trefoil as the dominant modes, all with much lower amplitudes of less than 0.15 λ/2π. Interestingly, spherical aberration under cryogenic conditions is weaker than anticipated, despite the expected influence of refractive index mismatch. This observation underscores the multifactorial and unpredictable nature of aberrations in cryo-FLM and highlights the limitations of relying solely on theoretical assumptions for PSF modeling.

#### Localization Error from PSF Mismatch

Wavefront distortions alter the PSF and consequently impact localization accuracy. To assess the impact of PSF mismatch, we evaluated the localization errors introduced when using standard theoretical (flat pupil) or RT-derived models, compared to the matched cryogenic PSF model (cryoT). We found that localization with the cryoT model achieves the highest accuracy, with root mean square errors (RMSE) of approximately 6 nm laterally and 30 nm axially with a signal photon count of 1 × 10^4^ (**Supplementary Table 1**). In contrast, the theoretical and RT models introduce substantial inaccuracies, with lateral RMSEs in the range of 10-20 nm and axial RMSEs around 200 nm (**Supplementary Table 2**). The localization performance also varied as a function of axial position (**Fig. 2e; Extended Data Fig. 2a-c**). Mismatched PSF models not only resulted in large localization biases but also increased localization uncertainty (**Fig. 2f**). Using the cryoT model, localization error distributions are tightly centered with low variability: −0.17 ± 8.92 nm (x), −0.25 ± 8.45 nm (y), and −0.10 ± 44.55 nm (z) (median ± interquartile range [IQR], N = 1500). In contrast, the theoretical and RT models resulted in pronounced lateral and axial biases, along with broader error distributions—representing a 20- to 70-fold increase in bias and a 2- to 6-fold increase in uncertainty.

Common fluorophores exhibit significantly enhanced photon emission at cryogenic temperatures due partially to markedly improved photostability. For example, a single GFP molecule has been reported to emit up to 1 million photons before photobleaching at cryogenic temperatures^39^. Motivated by this, we investigated localization inaccuracies under high-photon-count conditions (photon counts=4 × 10^5^) by comparing the RMSE to the corresponding Cramér-Rao lower bound (CRLB) (**Fig. 2g; Supplementary Notes 4.4**). Matched cryogenic PSF models yielded localization RMSE below 1 nm laterally and below 5 nm axially, approaching the theoretical limit set by the CRLB. In contrast, mismatched PSF models consistently produced elevated RMSEs, ranging from 7 to 15 nm laterally and 220 to 270 nm axially. Additionally, due to the complex aberrations present in cryo-FLM, we observe the standard theoretical PSF model performed significantly worse than in room-temperature FLM, particularly in the axial direction (**Extended Data Fig. 2d; Supplementary Notes 1.1**).

### FOV-dependent aberrations in cryo-FLM

An initial observation that aberrations in cryo-FLM are more complex than those in RT-FLM motivates a comprehensive investigation into their sources. In general, aberrations can be classified into three primary categories: (1) System-specific aberrations arising from intrinsic imperfections in the optical setup; (2) RI mismatch between the objective immersion medium (air, RI ≈ 1.00) and the sample medium (vitrified PBS, RI ≈ 1.33)^27^; (3) Inhomogeneous refractive indices within the specimen itself, including structural heterogeneity and composition gradients^19,40^. These aberrations can vary across both the FOV and the sample volume. In our experimental setup where fluorescent beads are suspended in PBS and vitrified within a thin layer of ice, all three types of aberrations are present. To investigate the spatial variability of aberrations in cryo-FLM, we examined three distinct scenarios: *FOV-dependent* aberrations, referring to variations in aberrations across different positions within a single FOV or nearby FOV (**Fig. 3a**); *square-dependent* aberrations, representing variations across different locations within the same TEM grid square; and *grid-dependent* aberrations, which capture broader aberration patterns across the entire sample, spanning the whole TEM grid.

**Fig. 3.**
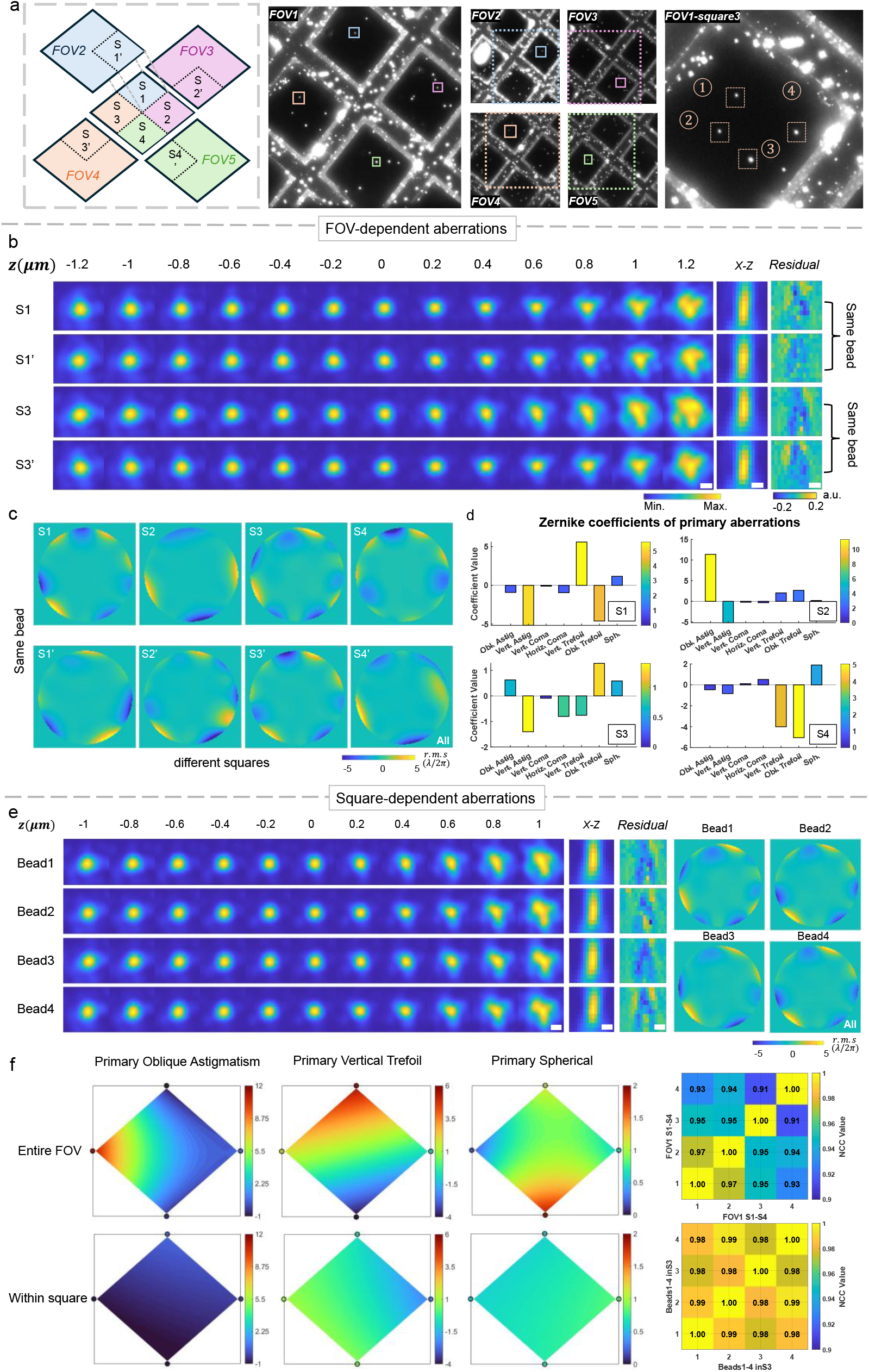
FOV-dependent and square-dependent aberrations in cryo-FLM. **a**. Imaging strategy designed to decouple system-specific aberrations from sample-induced ones. The same fluorescent beads were imaged in overlapping FOVs across multiple TEM grid squares (S1-S4 and S1′-S4′), allowing each bead to be captured at two distinct FOV-dependent positions, such as {S1, S1′}, {S2, S2′}, {S3, S3′}, and {S4, S4′}. **b**. Phase-retrieved PSFs and corresponding xz-view residuals for representative bead pairs imaged at different FOV positions (e.g., S1 vs. S1′ and S3 vs. S3′), illustrating variations arising solely from system-induced aberrations. **c**. Pupil phase maps of all aberrations for beads imaged in S1-S4 and S1′-S4′, revealing spatial variation in wavefront distortions (system-induced ones and combined aberrations) across the imaging field. **d**. Zernike coefficients for primary aberration modes (5^th^ −11^th^) from PSF models of the same beads in S1-S4, further highlighting FOV-dependent differences in aberration type and amplitude. **e**. Phase-retrieved PSFs and corresponding pupil phase maps of four different beads within the same grid square (FOV1-S3), showing relatively consistent aberration patterns due to reduced spatial variation at this scale. **f**. Contour maps of Zernike coefficients for four primary aberration modes across the full FOV (FOV1) and within a single grid square (FOV1-S3), along with partial normalized cross-correlation (NCC) values quantifying the similarity of corresponding PSFs. Scale bar: 500 nm.

To isolate the contributions of different aberration sources, we implement an imaging strategy that separates system-specific effects from sample-induced ones (**Fig. 3a; *Methods***). This setup allows each bead to be captured at two different FOV-dependent positions. By comparing images of the same bead across nearby FOVs, we eliminate variations caused by sample-induced aberrations. As a result, any differences in the retrieved PSF models of same bead reflect only system-specific aberrations across different positions within the FOV. This strategy also enables assessment of *FOV-dependent* aberrations (FOV1 S1-4 in **Fig. 3**) and *square-dependent* aberrations (FOV1 S3 Bead1-4 in **Fig. 3**), where both system and sample aberrations are simultaneously present.

#### System-Specific and FOV-Dependent Aberrations Show Strong Spatial Variability

We observed substantial variability in PSF models retrieved from identical beads imaged at different locations within nearby FOVs (e.g., FOV1 S1 and FOV2 S1’ in **Fig. 3a**), indicating that system-induced aberrations vary spatially across the FOV. Both the amplitude and type of aberrations, as well as pupil phase maps, differ markedly between positions (**Fig. 3b-c**; **Supplementary Fig. SS3**). Across the FOV (e.g., FOV1), the combined aberrations from both system- and sample-induced sources also show considerable spatial variability, differing in both nature and strength from the purely system-induced contributions (*S1 - S4* in **Fig. 3c**).

To quantify these discrepancies, we perform Zernike polynomial decomposition, revealing both qualitative and quantitative differences in primary aberration modes such as astigmatism, trefoil, and coma (**Fig. 3d**). For the same bead imaged across different positions within nearby FOVs (n = 4) which reveals the variety of system-induced aberrations, the partial normalized cross-correlation (PNCC, **Supplementary Notes 4.3**) of the retrieved PSF models range from 87% to 97% (**Supplementary Fig. 7**). The cosine similarity (**Supplementary Notes 4.1**) of the primary Zernike coefficients spans a broad range from −0.82 to 0.76, indicating that distinct aberration modes dominate at different positions. The corresponding root-mean-square (RMS) phase difference (**Supplementary Notes 4.1**) ranges from 0.89 λ/2π to 9.21 λ/2π, capturing both amplitude and modal differences in the aberrations (**Extended Data Fig. 3**). Even within the FOV containing combined system and sample-dependent aberrations, the PSF models display notable variability, with PNCC values between 91% and 97% (**Fig. 3f**), cosine similarity of the primary Zernike coefficients between −0.25 and 0.55, and RMS phase difference spanning 2.27 λ/2π to 10.60 λ/2π. These results indicate that both sample-induced and residual system-induced aberrations contribute to the observed variability, making the aberration landscape complex and spatially heterogeneous. Overall, these findings underscore the need to model spatially varying aberrations when constructing PSF models for high-precision single-molecule localization.

#### TEM Grid Square-Dependent Aberrations Display Spatial Consistency

Within a confined region, such as a single TEM grid square, the retrieved PSF models exhibit higher correlation and more consistent aberration profiles in both amplitude and dominant aberration types compared to those observed across the full imaging FOV (**Fig. 3e; Supplementary Fig. SS3**). Specifically, for four beads imaged within a single grid square (*FOV1-square* in **Fig. 3a**), the PNCC of the retrieved PSF models remain consistently high, ranging from 98% to 99% (**Fig. 3f, right**). The cosine similarity of the primary Zernike coefficients ranged from 0.69 to 0.97 for three beads, indicating the presence of similar aberration modes. One bead showed lower cosine similarity values, between 0.06 and 0.61, suggesting a modestly different aberration profile, though still more consistent than the variability observed across the entire FOV. The corresponding RMS phase difference ranged from 0.17 λ/2π to 1.41 λ/2π, which is about ten times smaller than the difference observed across the full FOV (**Extended Data Fig. 3**).

To further assess this spatial consistency, we compare the variability of Zernike coefficients for several primary aberration modes between the full FOV and a single TEM grid square. Within the square, the variability of dominant aberrations is only about 0.15% to 0.33% of the variability seen across the full FOV (**Fig. 3f, left**). These results indicate that aberrations are more spatially consistent within small imaging regions such as individual TEM grid squares. This supports the use of localized aberration modeling to enable subregion-specific PSF correction and improve the accuracy of high-precision localization.

#### Localization Error Induced by *FOV-dependent* and *Square-dependent* Aberrations in Cryo-FLM

To quantitatively assess the impact of spatially varying aberrations on localization accuracy in cryo-FLM, we perform 3D localizations using (i) a theoretical PSF model without aberrations, (ii) mismatched PSF models derived from the same or different TEM grid squares, and (iii) the matched *in situ* PSF model. The matched model yielded the highest accuracy, with localization bias of 0.36 ± 9.37 nm and −0.15 ± 10.00 nm laterally, and −1.36 ± 51.23 nm axially (median ± IQR, N = 1500, photon count=1 × 10^4^). In contrast, mismatched models introduced substantial bias and uncertainty, reaching up to 62.02 ± 11.83 nm and 85.27 ± 27.86 nm laterally, and 197.85 ± 97.12 nm axially (**Fig. 4a, b; Supplementary Fig. 5**) where axial localization was particularly sensitive to PSF mismatch, although mismatched models still outperformed the theoretical model (**Fig. 4c, d; Supplementary Fig. 6; Supplementary Fig. SS4**). Notably, using PSF models retrieved from three different beads within the same grid square partially mitigated localization bias, reducing errors to ranges of −6.93 ± 10.58 to −20.31 ± 15.73 nm and −0.02 ± 13.06 to 28.18 ± 13.19 nm laterally, and 59.59 ± 73.13 to 78.15 ± 79.07 nm axially. These results represent more than a threefold improvement compared to localization using mismatched PSF models from other grid squares or nearby FOVs (**Fig. 4e; Extended Data Fig. 4b; Supplementary Fig. 10; Supplementary Fig. SS5**).

**Fig. 4.**
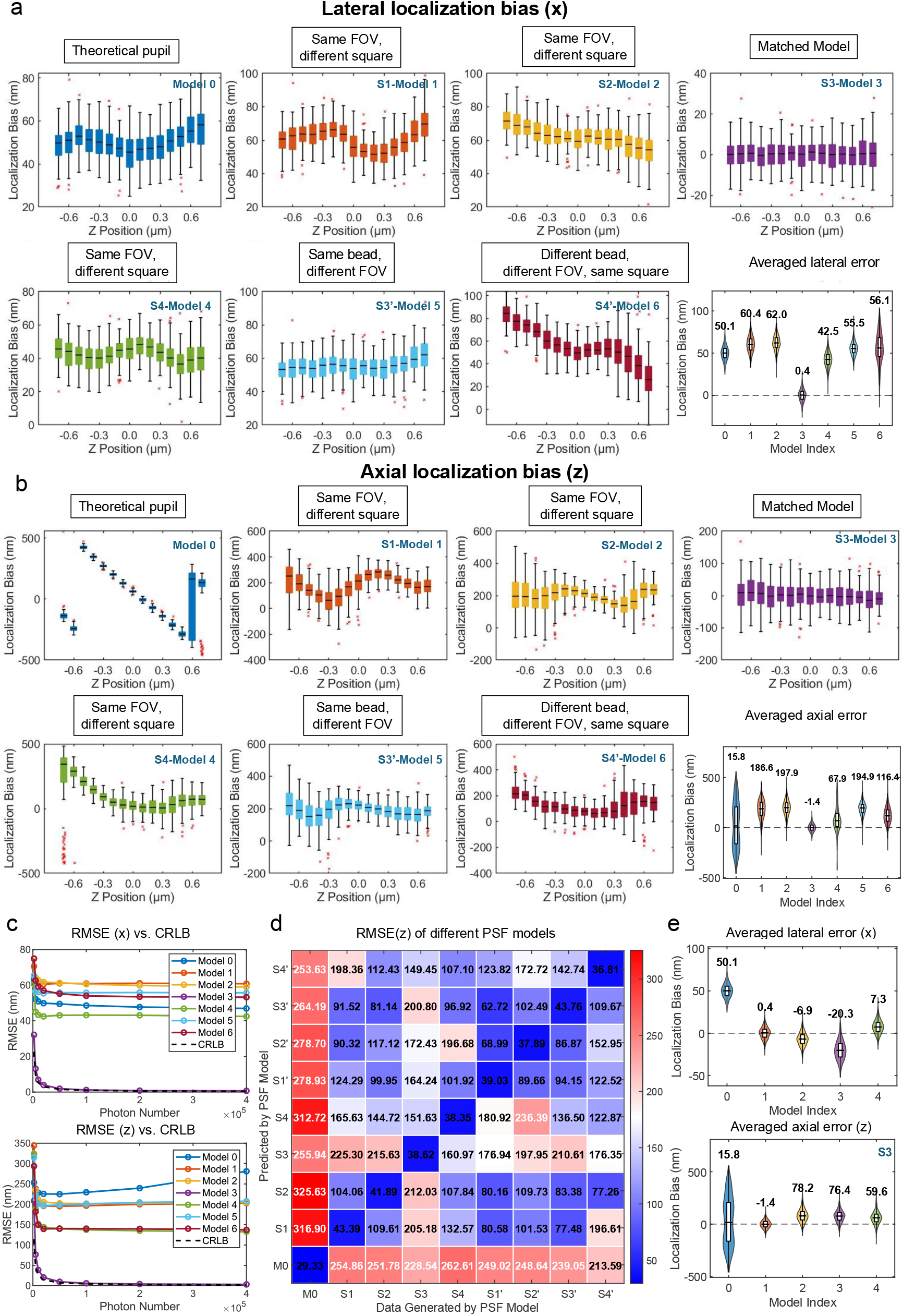
Localization bias induced by FOV- and square-dependent PSF mismatches in cryo-FLM. Test data were generated using the PSF model retrieved from a bead in FOV1-S3 (**Fig. 3a**), spanning axial positions from −0.7 µm to +0.7 µm in 100 nm steps (100 frames per z-position). Simulations were performed with a signal photon count of 1 × 10^5^ and a background photon count of 400. Seven PSF models were used for 3D localization: Model 0 (theoretical pupil function), Models 1-4 (experimental PSFs from beads in FOV1-S1 to S4), and Models 5-6 (PSFs from S3′ and S4′, representing system-induced and sample-induced aberrations, respectively). **a-b**. Lateral (x) and axial localization bias as a function of axial position, and corresponding average localization bias across all z positions. **c**. Localization RMSE compared to the CRLB across varying photon counts. **d**. Summary of axial localization bias when using theoretical and experimental PSFs from S1-S4 and S1′-S4′ for both data simulation and localization, highlighting performance degradation due to PSF model mismatches. **e**. Averaged lateral (x) and axial localization bias using PSF models retrieved from four beads within the same grid square (FOV1-S3), illustrating improved consistency when models are locally matched. Scale bar (if applicable): 500 nm.

**Fig. 5.**
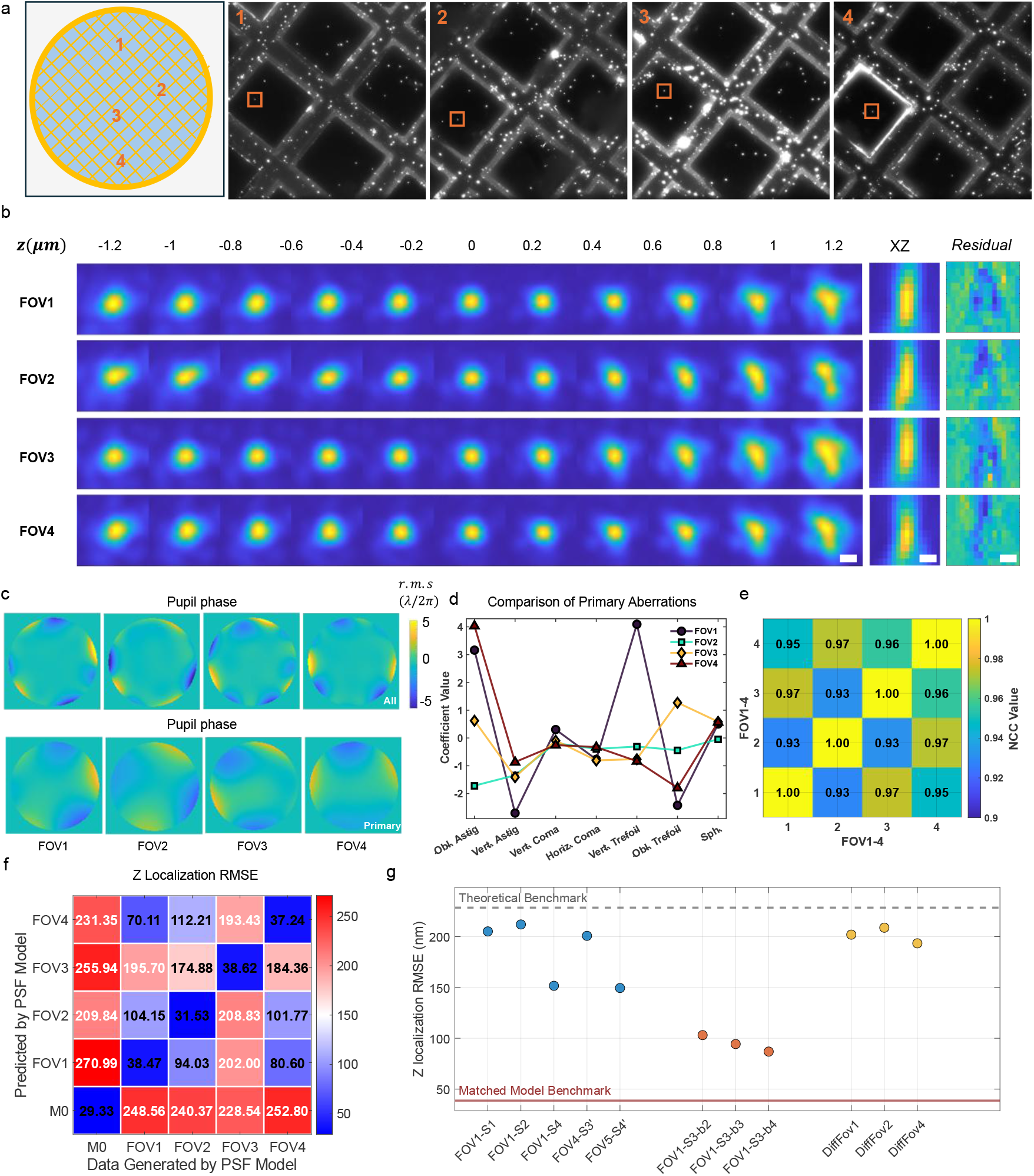
Aberrations across the TEM grid induced by refractive index mismatch and sample heterogeneity. **a**. Overview of four distinct FOVs distributed across the TEM grid. Beads were selected from consistent relative positions within each FOV to assess spatial variability in aberrations. **b**. Phase-retrieved PSFs for the selected beads, displayed alongside xz-view residual maps showing differences between experimental and reconstructed PSFs. **c**. Pupil phase maps illustrating total aberrations (top row) and primary aberrations (bottom row) for each bead. **d**. Zernike polynomial coefficients corresponding to the primary aberration modes retrieved from each PSF model. **e**. PNCC values computed among the four phase-retrieved PSF image stacks, reflecting the similarity of aberration profiles across FOVs. **f**. Axial localization RMSE introduced by matched and mismatched PSF models during both data simulation and fitting. **g**. Axial localization RMSE as a function of *FOV-dependent, square-dependent*, and *grid-dependent* aberration mismatches. Test data were generated using the PSF model from FOV3 (identical to the PSF model used in FOV1-S3 and FOV1-S3-b1 in **Fig. 3a**). Scale bar: 500 nm.

Increasing photon counts did not eliminate these errors. Even at 4 × 10^5^ photons, localization RMSEs remained at 40-90 nm laterally and 130-280 nm axially for *FOV-dependent* aberrations. The *square-dependent* mismatches showed modest improvement (7-30 nm laterally; 60-110 nm axially) but remained significantly worse than the matched PSF model, which achieved sub-nanometer lateral (0.68 nm for x, 0.94 nm for y) and sub-5 nm axial (3.25 nm) accuracy (**Fig. 4c**). These results demonstrate that only spatially matched *in situ* PSF models enable nanometer-precision localization in cryogenic fluorescence microscopy.

### Aberrations across the whole TEM grid

#### TEM Grid-Dependent Aberrations Are Highly Variable and Unpredictable

Because vitrified samples typically exhibit heterogeneous refractive index distributions in cryo-FLM, we investigated the spatial variation of aberrations across the entire TEM grid. To minimize the influence of the system-induced FOV-dependent aberrations in our system (**Fig. 3**) and focus specifically on aberrations originating from the sample itself, such as those caused by refractive index mismatches and local inhomogeneities, we selected four fluorescent beads located at similar relative positions across four different FOVs (**Fig. 5a)**.

Although phase-retrieved PSFs from different beads show similar elongation and tilt in the xy plane, clear differences emerge in their overall shape, particularly in the xz view (**Fig. 5b**). While the selected FOVs exhibit comparable primary aberration types, including astigmatism, trefoil, weaker coma, and spherical aberration, the amplitudes of these aberrations vary across regions. This variability is evident in the retrieved pupil phase maps (**Fig. 5c**) and the corresponding Zernike coefficients (**Fig. 5d**). Cosine similarity between the primary Zernike modes ranges from

-0.43 to 0.56, and the RMS phase differences span 0.08 λ/2π to 4.11 λ/2π (**Extended Data Fig. 3a, b**). Among the four beads, the maximum amplitude differences reach up to 5.7 λ/2π for astigmatism and 4.9 λ/2π for trefoil. These differences are smaller than those observed in *FOV-dependent* aberrations, which incorporate both sample-induced and system-specific components. This interpretation is further supported by the high PNCC values between PSFs, ranging from 0.93 to 0.97 (**Fig. 5e**), which means the grid-dependent aberrations (sample-induced aberrations) are not as significant as FOV-dependent aberrations (combined system and sample-induced aberrations) in this case.

To evaluate the impact of these aberrations on localization accuracy, we compared localization RMSEs obtained using (i) the theoretical PSF model (*M0* in **Fig. 5f**), (ii) mismatched PSF models from other selected FOVs, and (iii) the matched *in situ* PSF model. As expected, the matched PSF model yielded the highest localization precision (**Fig. 5f, Extended Data Fig. 4a**), with RMSEs of ∼7 nm laterally and ∼40 nm axially (N = 1500). In contrast, mismatched PSF models introduced spatially variable and often unpredictable localization errors, with worst-case errors reaching ∼60 nm laterally and ∼200 nm axially. In some instances, however, localization performance approached that of typical *FOV-dependent* results (**Fig. 3**), with errors reduced to <30 nm in y and <110 nm in z. Occasionally, the accuracy approaches that of the matched PSF model.

We further compared the localization bias induced by *FOV-, square-*, and *grid-dependent* aberrations. In this analysis, the same PSF model (FOV1-S3 and FOV1-S3-Bead1 from **Fig. 3a**; FOV3 from **Fig. 5a**) was used for data simulation, while mismatched PSF models from the corresponding sets were used for localization analysis. We found the z-localization RMSE remains high for both FOV-dependent and grid-dependent aberrations (**Fig. 5g**). In contrast, the lateral (x and y) localization errors for grid-dependent aberrations are comparatively low—approaching those observed in square-dependent cases (**Extended Data Fig. 4b**). This trend aligns with the high similarity seen in the XY projections of the PSFs (**Supplementary Fig. SS7**), suggesting that lateral localization is preserved when XY shape similarity is maintained. Since grid-dependent aberrations primarily reflect sample-induced effects, they tend not to alter the XY shape of the PSF drastically, which explains the preserved lateral localization accuracy. However, axial localization proves far more sensitive to subtle aberrations, particularly those introduced by sample-specific effects. Overall, these results demonstrate that aberrations vary unpredictably across FOVs, squares, and grids, producing severe axial biases and substantial lateral errors that exceed cryo-ET precision requirements. Although PSF models from adjacent beads can reduce these errors by 3–10 fold, only true in situ PSF models consistently achieve sub-1 nm lateral and sub-5 nm axial accuracy, enabling reliable molecular localization (**Supplementary Notes 1, Supplementary Fig. 12; Supplementary Fig. SS8**).

## Discussion

Cryo-FLM plays a vital role in cryo-CLEM, providing positional information to guide downstream cryo-FIB milling and structural analysis by cryo-ET. However, the reliability of these workflows depends critically on accurate 3D localization of fluorescent markers. Unlike room-temperature fluorescence microscopy, cryo-FLM suffers from more complex and unpredictable aberrations, including those from the imaging system, refractive index mismatches, and sample-induced heterogeneity. These aberrations distort the PSF, leading to significant localization errors. Combined with Poisson noise, limitations in illumination powers, and autofluorescence under cryogenic conditions, aberration correction in cryo-FLM remains a major challenge.

In this study, we established an experimental and analytical pipeline to systematically characterize and evaluate optical aberrations across the cryo-CLEM interface using an established commercial cryo-FLM system. This approach enabled a quantitative assessment of aberrations across system, sample, and refractive index mismatch dimensions, revealing the extent and origin of localization errors in realistic cryo-imaging conditions. Specifically, we found: 1. The aberrations in cryo-FLM are much more complicated than room temperature FLM, changing both in amplitude and type of aberration. Interestingly, spherical aberration under cryogenic conditions is weaker than anticipated, despite the expected influence of refractive index mismatch. This observation underscores the multifactorial and unpredictable nature of aberrations in cryo-FLM and highlights the limitations of relying solely on theoretical assumptions for PSF modeling. 2. Both the system-specific and sample-induced aberrations vary across the FOV and TEM grid, making it complex and unpredictable. 3. The FOV-dependent aberrations introduce substantial 3D localization biases (up to 90 nm laterally and more than 300 nm axially) that exceed acceptable thresholds for downstream cryo-ET workflows even when using infinite photon numbers. Interestingly, sample-induced effects tend not to drastically distort the XY shape of the PSF, thereby preserving lateral localization accuracy. This suggests that lateral precision in CLEM is more resilient to sample-induced aberrations (though not to system-specific aberrations). In contrast, axial precision which is poorly characterized remains highly susceptible to such distortions.. Cryo-FIB-milled lamellae are typically only 150-200 nm thick, making a 300 nm axial localization error highly problematic. 4. PSF models from spatially adjacent beads within a single TEM grid square show consistent aberration profiles and offer partial mitigation of localization bias, though they remain inferior to true *in situ* models. These findings underscore the need for precise, localized aberration characterization and correct PSF models in cryo-FLM to achieve precise molecular localization in cryogenic imaging workflows. It is important to note that the sample used in this study represents an idealized condition, consisting solely of PBS and fluorescent beads. In more complex biological specimens, such as cells or tissue sections, sample-induced aberrations are expected to be stronger and more spatially heterogeneous.

In this work, we used a scalar PSF model for phase retrieval and localization analysis. While vectorial PSF models account for polarization effects and high-angle emission in high-NA systems, the scalar approximation remains widely used due to its computational efficiency and suitability for a broad range of microscopy applications. Given that cryo-FLM systems operate at a moderate numerical aperture and that our fluorescent emitters are embedded in vitrified samples with relatively isotropic emission, the scalar model provides a sufficiently accurate representation of the system PSF. Moreover, the primary aberration types we observe, such as astigmatism, coma, and spherical aberration, are well captured within the scalar framework. The agreement between phase-retrieved PSFs and experimental bead images, along with the high localization precision achieved using matched PSF models, further supports the adequacy of the scalar model for assessing aberrations and guiding localization in cryo-FLM.

While we used a common commercial cryo-FLM system, other systems may utilize different objectives or configurations that could have substantial effects on aberrations. Our Leica system utilizes a ceramic-tipped objective that is operated cold; other systems often use longer working distance objectives that operate warm. For this reason, we have provided code, instructions and evaluation protocols to aid cryo-FLM users to evaluate these effects on their own systems.

Future directions in this field may include improved phase retrieval under high-noise conditions, adaptive *in situ* PSF estimation strategies, and the development of 3D localization-enabled cryo-FLM systems. Progress in these areas would further enhance the accuracy and utility of cryo-CLEM techniques for *in situ* structural biology. Together, these efforts aim to overcome current limitations in cryo-FLM, enabling accurate, reliable, and high-resolution localization of fluorescent markers in complex cryogenic samples. Ultimately, we hope this study will help to advance the precision and effectiveness of cryo-CLEM workflows for *in situ* structural biology.

## Online Methods

### Preparation of fluorescent beads sample under cryogenic temperature

To prepare fluorescent bead samples for cryo-FLM, 200-nm diameter TetraSpeck fluorescent beads (Thermo Fisher Scientific) were diluted 1:50 in PBS to achieve a sparse distribution suitable for single-molecule imaging. To prevent aggregation and ensure uniform dispersion, the stock solution was sonicated for 5 minutes prior to dilution, and the diluted suspension was vortexed briefly immediately before application. To minimize photobleaching, all sample handling was performed under low-light conditions, and samples were wrapped in aluminum foil prior to freezing.

Sample vitrification was carried out using an automated plunge-freezing system (Vitrobot Mark IV, Thermo Fisher Scientific). Holey carbon-coated TEM grids (CF-2/2-300Au) were first glow-discharged using an Emitech K100X Glow Discharge Unit to remove surface contaminants and enhance hydrophilicity, thereby promoting uniform spreading of the aqueous sample. Samples were plunge-frozen into liquid ethane and stored in a liquid nitrogen dewar until imaging.

### Preparation of fluorescent beads under room temperature

The sample preparation protocol for experiments involving imaging the same fluorescent bead sample under both room-temperature and cryogenic conditions deviated from conventional cryo-FLM workflows. A stock solution of 200-nm diameter TetraSpeck fluorescent beads (Thermo Fisher Scientific) was diluted 1:100 in PBS and prepared as above. To enable identification and localization of the same beads across imaging conditions, we coated gold London Finder TEM grids (Quantifoil 2/2-200LF) with a thin layer of amorphous carbon prior to glow discharge. The TetraSpecks were adhered to the carbon layer by applying a 3-µL droplet of the diluted bead suspension and allowing it to settle for 5 seconds before wicking away excess fluid with Whatman paper, leaving a uniform, thin distribution of beads. Prepared grids were stored in the dark overnight at room temperature to ensure complete evaporation of residual liquid. This drying step promoted strong adhesion of the beads to the carbon film and preserved their spatial distribution— critical for accurately comparing PSFs between room-temperature and cryogenic imaging.

To prepare the same grid for cryo-FLM while maintaining the original bead positions, a 3-µL droplet of DI water was gently applied to the dried grid before plunge-freezing on a Vitrobot Mark IV as above.

### Microscope Setup

Imaging was performed using a Leica THUNDER Imager EM Cryo CLEM system (Leica Microsystems), a fully motorized upright widefield fluorescence microscope optimized for cryo-CLEM. The system is built around a DM6 FS microscope body and equipped with a 50×/0.90 NA cryo-objective and cryo-stage to maintain the sample below −135°C throughout imaging. Fluorescence excitation was provided by a white lamp source delivered through filter cubes optimized for GFP (excitation 470/40, DCR 495, emission 525/50). Fluorescence was collected through the same objective and detected using a Leica DFC9000 GT deep-cooled sCMOS camera, offering high sensitivity (82% QE), low noise, and fast acquisition speeds (50 fps). The system was controlled via LAS X software, which supported sample navigation, autofocus, image stitching, and export of coordinate data in CLEM-compatible formats. The combination of precise cryo handling, high-resolution imaging, and CLEM-ready software enabled accurate correlation of fluorescence and electron microscopy data.

### Data acquisition

The dataset collection was performed separately for cryo-temperature and room-temperature imaging while maintaining positional consistency across both conditions.

- **Cryo-Temperature Imaging**. The vitrified sample was mounted onto a cryo-stage within the Leica EM Cryo CLEM microscope, ensuring stable cryogenic conditions throughout the imaging process. All handling and mounting steps were performed within a cryo workstation cooled by liquid nitrogen to prevent devitrification and minimize ice contamination. Prior to imaging, a series of bright-field and green-channel fluorescence tiles were acquired at rough focus. These sub-regions were then computationally stitched into a complete montage of the EM grid, enabling precise identification of regions of interest (ROIs) based on both the spatial distribution of fluorescent beads and ice thickness. Following ROI selection, fluorescence imaging was performed by acquiring a z-stack with a step size of 200 nm. At each axial position, 10 frames were collected and averaged to improve the signal-to-noise ratio. Imaging was conducted in low-noise mode using a camera offset of approximately 100 and a gain setting of 7. Given that fluorophores exhibit minimal photobleaching under cryogenic conditions, a higher illumination intensity was employed to further enhance signal quality while retaining vitrification (**more details in Supplementary Notes 5**).
- **Room-Temperature Imaging**. To enable room-temperature imaging while maintaining consistent sample positioning with cryo-FLM imaging, fluorescence imaging was first performed under cryogenic conditions as described above. Following cryo-imaging, the sample and stage insert were removed and stored at room temperature in the dark to dry, while the microscope’s cryostage was baked out at 60 C to defrost and remove condensation. Because the sample remained securely mounted in the designated stage, we preserved its spatial orientation for subsequent imaging. Accurate repositioning was further facilitated by reference markings on the Finder EM grid. After the microscope and sample were both dry and at room temperature, room-temperature imaging was carried out using the same imaging workflow as for cryo-FLM: bright-field and fluorescence tiles were collected for grid-wide montage generation, followed by ROI selection (corresponding to the same locations imaged under cryo conditions), and z-stack acquisition. To account for the increased susceptibility of fluorophores to photobleaching at room temperature, a lower lamp intensity was used during fluorescence excitation. All other imaging parameters, including low-noise acquisition mode, a z-step size of 200 nm, and 10-frame averaging at each axial position were kept consistent with the cryo-FLM setup to ensure data comparability across both temperature conditions.

### MLE-Based Phase Retrieval Algorithm

#### Aberrated PSF Model

We modeled the aberrated PSF using scalar diffraction theory, in which the PSF intensity at a spatial position (*x, y, z*) is computed as the squared amplitude of the inverse Fourier transform of the pupil function. The pupil function encodes both amplitude and phase, with phase deviations arising from optical aberrations expressed via a Zernike polynomial expansion. The aberration-influenced PSF intensity is defined as:

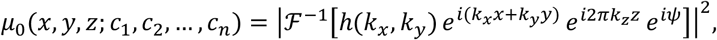

where *h*(*k*_*x*_, *k*_*y*_) denotes the amplitude of the pupil function, 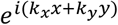 accounts for lateral spatial phase shifts, and 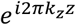 represents defocus. The axial spatial frequency *k*_*z*_ is given by:

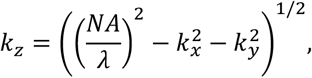

with *NA* as the numerical aperture of the objective and *λ* the emission wavelength.

The phase distortion *ψ*, caused by aberrations, is modeled as a weighted sum of Zernike polynomials:

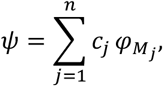

Where 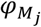 represents the Zernike mode associated with aberration type *M*_*j*_, and *c*_*j*_ are the mode amplitudes to be estimated. This representation provides a compact and interpretable basis for modeling complex phase distortions in high-resolution imaging systems.

To account for experimental variability, such as differences in photon counts and background levels across focal planes, we introduced scaling terms. The modeled intensity for image *k* becomes:

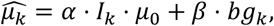

where *α* is a global photon count scaling factor, *I*_*k*_ is the estimated total photon count at plane *z*_*k*_, *β* is the background scaling coefficient, and *bg*_*k*_ is the estimated background level at plane *z*_*k*_.

Large fluorescent beads are substantially brighter than smaller ones. They contain more fluorophores and thus show less intensity fluctuations and photobleaching. As the bead images can be described as a convolution of the PSF with the bead shape, they are more blurred than the PSF itself. We can include this blurring in the forward model by a convolution with a uniform fluorophore distribution in the bead^41^:

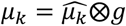

#### Maximum Likelihood Estimation

The workflow of our MLE-based phase retrieval algorithm is shown in **Extended Data Fig. 1**. To estimate the optimal Zernike coefficients *c*_*j*_, along with the global intensity scaling factor *α* and background contribution *β*, we formulated a Poisson likelihood model. We assumed that the photon counts *X*_*ks*_ at each pixel *s* in image *k* follows an independent Poisson distribution with expectation given by the modeled intensity:

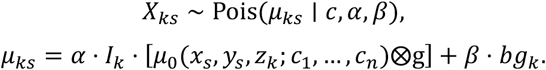

Given a set of *K* images acquired at focal positions {*z*_1_, *z*_2_, …, *z*_*K*_}, with each image containing *S* pixels, the joint likelihood is:

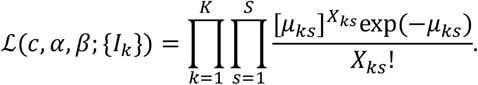

We obtained the maximum likelihood estimates by minimizing the negative log-likelihood:

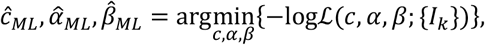

which expands to:

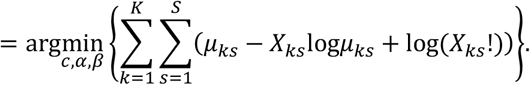

Since the final term log (*X*_*ks*_!) is independent of the parameters, it is omitted during optimization.

After estimating the optimal parameters 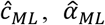 and 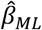,we reconstructed the aberration phase function *ψ*, pupil function, and the corresponding PSF. This phase-retrieved PSF model accounts for system aberrations, photon count variations, and background contributions, enabling accurate PSF characterization and aberration correction for 3D single-molecule localization microscopy.

#### Data Preprocessing

To prepare the single-bead image stacks for phase retrieval, we performed a targeted preprocessing workflow designed to isolate the fluorescent signal while minimizing background interference (**Supplementary Fig. 2-3**). For each bead, two distinct regions were selected: one centered on the bead to capture the signal, and another in a bead-free area to estimate the background. Images were cropped to a region of size *R*_1_ × *R*_1_ pixels, where *R*_1_ *=* 3, corresponding to an approximately 4 *μ*m×4 *μ*m field of view. This cropping preserved the bead signal while removing peripheral regions dominated by background noise.

To ensure alignment of the bead across axial planes, the in-focus image was fit to a 2D Gaussian intensity profile. The resulting centroid was used to align all cropped frames, maintaining spatial consistency across the z-stack. Photon count estimation was then performed for both the signal and background regions. The raw intensity values in both regions were first converted to photon counts using calibration parameters. For each frame, the background photon count was computed by averaging the photon values in the background region. This background estimate was subtracted from the corresponding bead frame to isolate the bead signal. The total number of photons per frame was then calculated by summing the corrected values in the signal region. These estimated signal and background photon counts were used to initialize the global intensity (signal) scaling factor *α* and background scaling factor *β* in the MLE-based phase retrieval algorithm. This initialization provided an optimized starting point for robust and accurate estimation of aberration parameters.

### Metrics

#### PSF model difference

To quantify differences in aberrations between PSF models, we used two complementary metrics: the cosine similarity of the Zernike polynomial coefficients and the root mean square (RMS) phase error. Cosine similarity captures the angular similarity of aberration modes, indicating how closely the aberration types align, while RMS phase error reflects the overall amplitude of phase distortion across the pupil. Together, these metrics provide insight into both the nature and severity of aberration differences between FOV-, square-, and grid-dependent PSF models (more details in **Supplementary 4.1**).

#### Localization error

Localization accuracy was assessed using both the median localization error with interquartile range (median ± IQR) and the RMSE. The median ± IQR highlights the central tendency and variability of localization distributions, offering a robust measure less sensitive to outliers. RMSE, in contrast, captures both bias and variance across the full dataset, providing an overall measure of localization accuracy. These two metrics together enable a comprehensive evaluation of localization performance under different aberration conditions (more details in **Supplementary 4.2**).

## Data availability

The authors declare that data supporting the findings of this study are available within the paper and its Supplementary Information.

## Code availability

We provide three MATLAB toolboxes as supplementary software for this manuscript: MLE-based phase retrieval algorithm, data simulation for biplane setup, and 3D localization for biplane setup. All three toolboxes are available as Supplementary Software with the manuscript. Future updates and maintenance will be freely available at: https://github.com/LAMetskas/2025_CryoFLMAberrations/.

## Acknowledgments

The authors thank Louise Bertrand from Leica Microsystems, Inc. for providing technical and applications support. This work was supported by NIAID award 1DP2AI164293-01 to L.A.M. and NIGMS MIRA award R35GM119785 to F.H.

## Author contributions

H.L, L.A.M and F.H conceived the idea. H.L and L.A.M conducted the experiments. H.L wrote the codes and performed the analysis. F.H and L.A.M reviewed and revised the paper. F.H and L.A.M supervised the project.

---

**Extended Data Figure 1.**
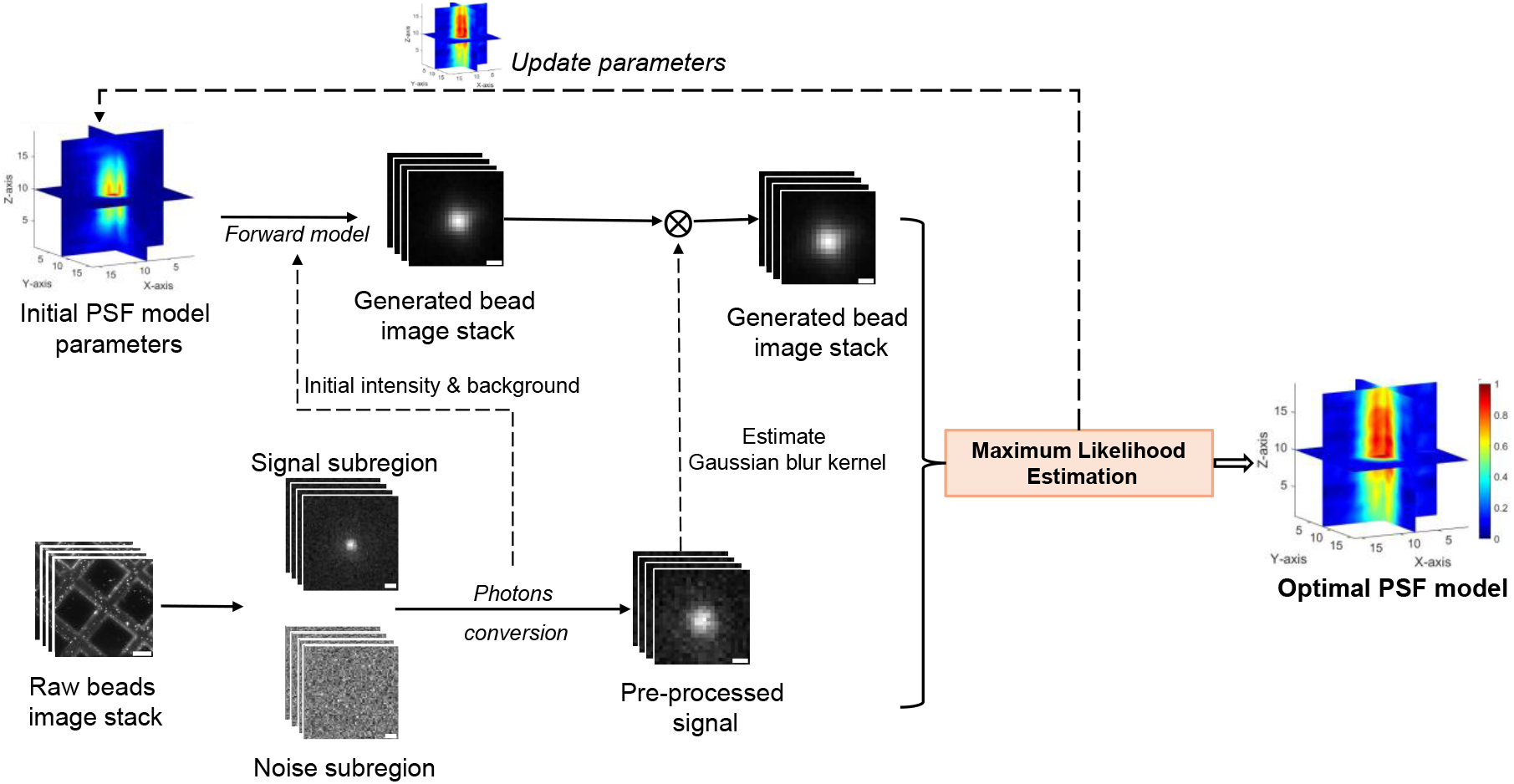
The workflow of MLE based phase retrieval algorithm.

**Extended Data Figure 2.**
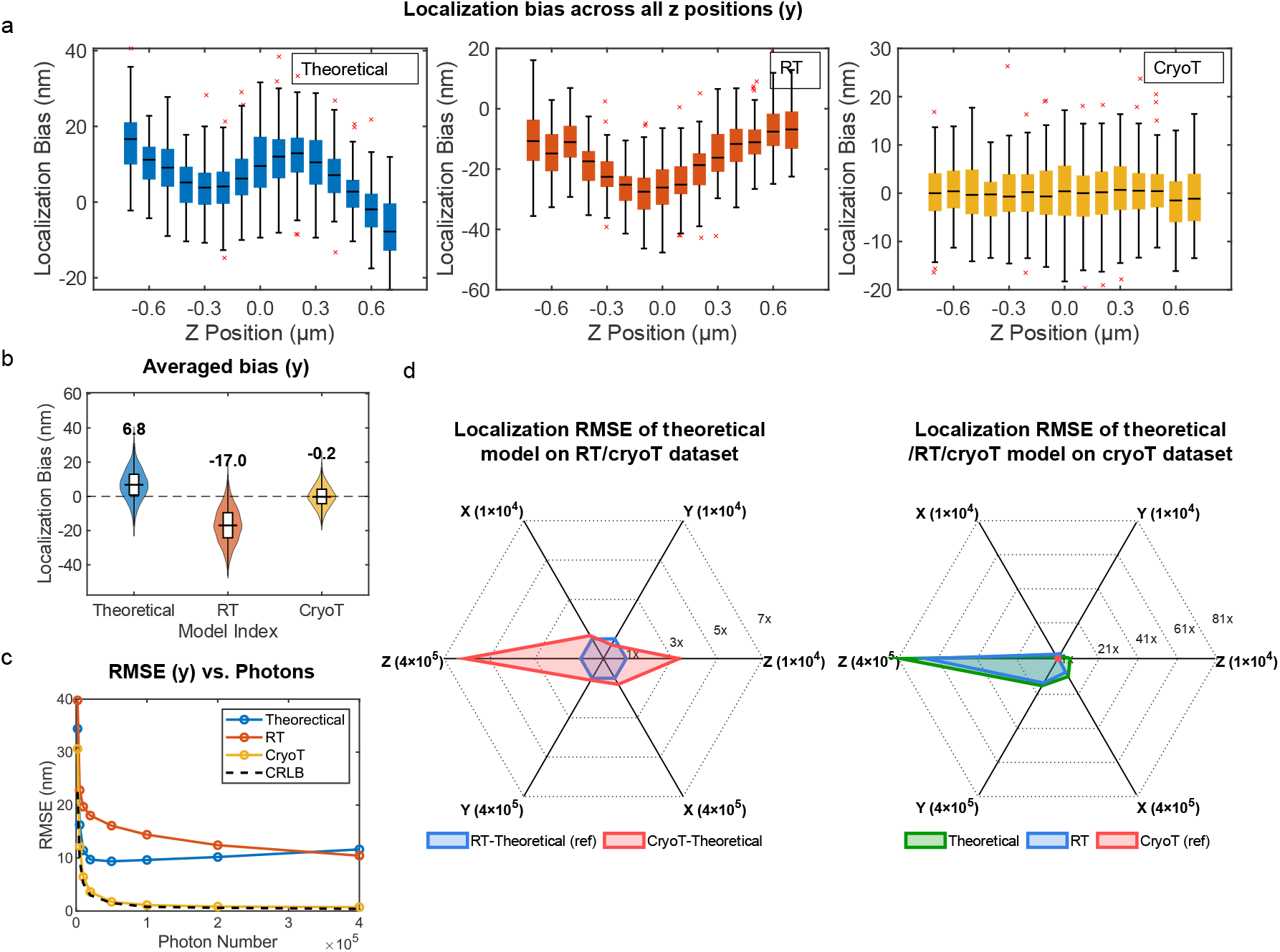
Localization analysis of room-temperature versus cryogenic fluorescence microscopy of the same bead. **a**. Localization bias in the y-dimension across axial positions (from −0.7 µm to +0.7 µm in 100 nm increments; 100 frames per position; simulation model: cryoT), analyzed using three PSF models: (1) theoretical pupil function, (2) RT model, and (3) cryoT model. Simulations were performed with a signal photon count of 1 × 10^5^ and a background photon count of 400. **b**. Localization bias distributions aggregated over all positions for y. **c**. RMSE in y localization versus the CRLB as a function of photon count. **d**. Left: Spider plot comparing localization performance using the theoretical model on test datasets generated by the RT and cryoT PSF models. Simulations were performed with a signal photon count of 1 × 10^4^ and 4 × 10^5^, and a background photon count of 400. Performance on the RT dataset is used as the baseline (unit circle); the red polygon shows the ratio of performance on the cryoT dataset to the RT dataset. Right: Spider plot comparing localization performance on a cryoT-generated test dataset using three PSF models (theoretical, RT, and cryoT). Simulations were performed with a signal photon count of 1 × 10^4^ and 4 × 10^5^, and a background photon count of 400. Performance using the cryoT model is the baseline; the green and blue polygons indicate the performance ratios of the theoretical and RT models relative to the cryoT model.

**Extended Data Figure 3.**
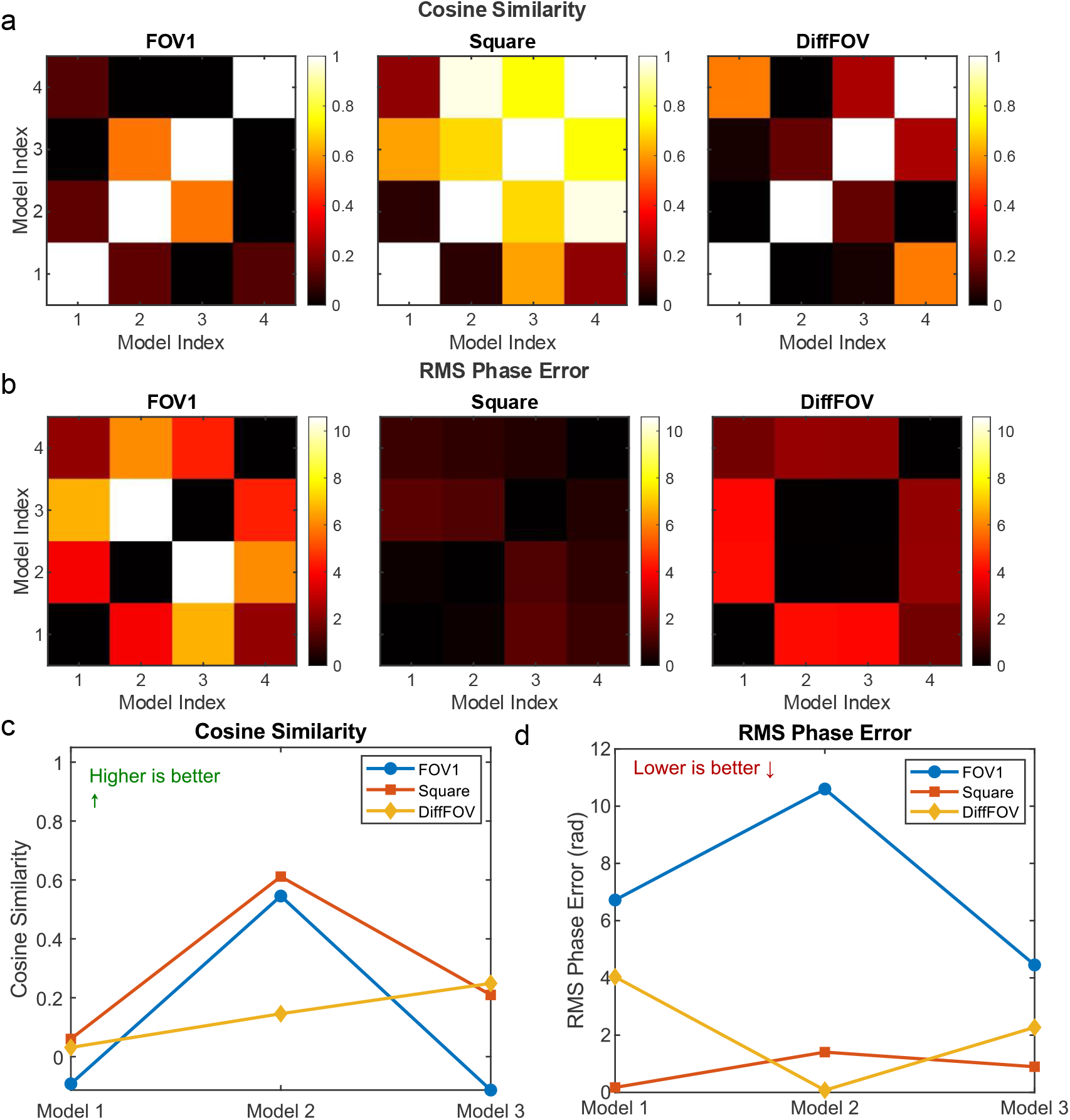
Comparison of FOV-dependent (FOV1), square-dependent (Square), and grid-dependent aberrations (DiffFOV) The primary aberration set includes Zernike polynomials of Noll orders 5^th^ to 11^th^. **a**. Cosine similarity matrix of Zernike coefficients within FOV-dependent (FOV1), square-dependent (Square), and grid-dependent (DiffFOV) aberration groups. **b**. RMS phase error matrix for the same three aberration groups. **c**. Direct comparison of cosine similarity and RMS phase error for one representative PSF model in each group. The same PSF model is used across all three comparisons—S3 in FOV1 (FOV-dependent, **Fig. 3a**), Bead 1 in Square1 (square-dependent, **Fig. 3a**), and FOV3 in a different FOV (grid-dependent, **Fig. 5a**). Comparisons are also made with three other models from the same respective groups to evaluate intra-group variability.

**Extended Data Figure 4.**
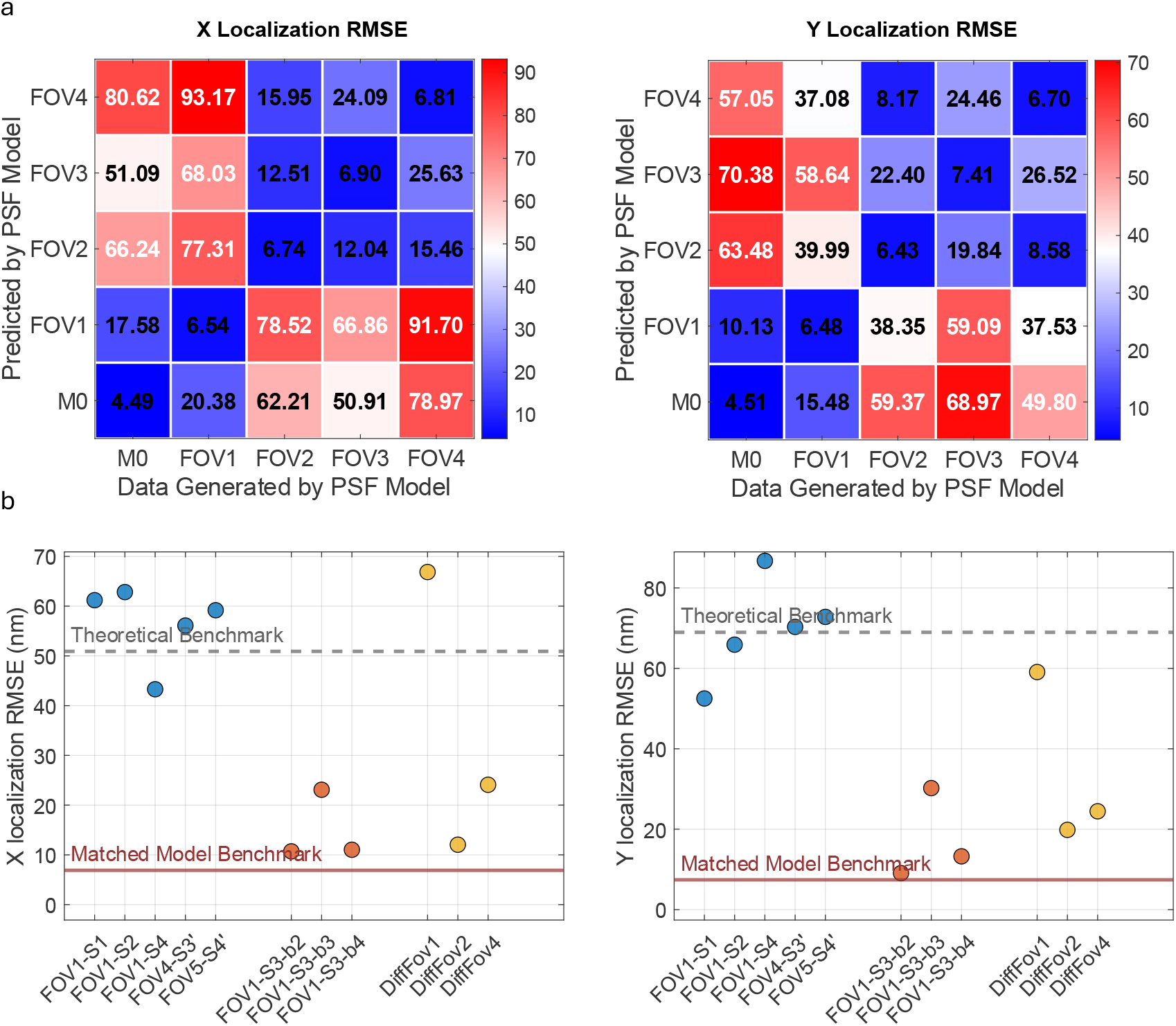
Localization errors induced by FOV-, square-, and grid-dependent aberrations. **a**. Lateral (x and y) localization RMSE induced by grid-dependent aberrations across different regions of the imaging field. **b**. Test datasets were generated using the PSF model from FOV3 in **Fig. 5a** (identical to the PSF models used in FOV1-S3 and FOV-S3-b1 in **Fig. 3a**), spanning axial positions from −0.7 μm to +0.7 μm in 100 nm steps, with 100 frames per z-position. Simulations were conducted with a signal photon count of 1 × 10^5^ and a background photon count of 400 photons per pixel. Comparison of localization RMSE in x, and y directions arising from FOV-dependent (FOV1), square-dependent (Square), and grid-dependent (DiffFOV) aberrations.

## Reference

(1) Metskas, L. A.; Ortega, D.; Oltrogge, L. M.; Blikstad, C.; Lovejoy, D. R.; Laughlin, T. G.; Savage, D. F.; Jensen, G. J. Rubisco Forms a Lattice inside Alpha-Carboxysomes. Nat Commun 2022, 13 (1). 10.1038/S41467-022-32584-7.

(2) Tegunov, D.; Xue, L.; Dienemann, C.; Cramer, P.; Mahamid, J. Multi-Particle Cryo-EM Refinement with M Visualizes Ribosome-Antibiotic Complex at 3.5 Å in Cells. Nat Methods 2021, 18 (2), 186–193. 10.1038/S41592-020-01054-7.

(3) Hutchings, J.; Stancheva, V.; Miller, E. A.; Zanetti, G. Subtomogram Averaging of COPII Assemblies Reveals How Coat Organization Dictates Membrane Shape. 10.1038/s41467-018-06577-4.

(4) Schur, F. K. M.; Obr, M.; Hagen, W. J. H.; Wan, W.; Jakobi, A. J.; Kirkpatrick, J. M.; Sachse, C.; Kräusslich, H. G.; Briggs, J. A. G. An Atomic Model of HIV-1 Capsid-SP1 Reveals Structures Regulating Assembly and Maturation. Science 2016, 353 (6298), 506–508. 10.1126/SCIENCE.AAF9620.

(5) Zivanov, J.; Otón, J.; Ke, Z.; von Kügelgen, A.; Pyle, E.; Qu, K.; Morado, D.; Castaño-Díez, D.; Zanetti, G.; Bharat, T. A. M.; Briggs, J. A. G.; Scheres, S. H. W. A Bayesian Approach to Single-Particle Electron Cryo-Tomography in RELION-4.0. Elife 2022, 11. 10.7554/ELIFE.83724.

(6) Castaño-Díez, D.; Kudryashev, M.; Arheit, M.; Stahlberg, H. Dynamo: A Flexible, User-Friendly Development Tool for Subtomogram Averaging of Cryo-EM Data in High-Performance Computing Environments. J Struct Biol 2012, 178 (2), 139–151. 10.1016/J.JSB.2011.12.017.

(7) Chaillet, M. L.; van der Schot, G.; Gubins, I.; Roet, S.; Veltkamp, R. C.; Förster, F. Extensive Angular Sampling Enables the Sensitive Localization of Macromolecules in Electron Tomograms. Int J Mol Sci 2023, 24 (17). 10.3390/IJMS241713375.

(8) Schorb, M.; Briggs, J. A. G. Correlated Cryo-Fluorescence and Cryo-Electron Microscopy with High Spatial Precision and Improved Sensitivity. Ultramicroscopy 2014, 143, 24–32. 10.1016/J.ULTRAMIC.2013.10.015.

(9) Rigort, A.; Bäuerlein, F. J. B.; Villa, E.; Eibauer, M.; Laugks, T.; Baumeister, W.; Plitzko, J. M. Focused Ion Beam Micromachining of Eukaryotic Cells for Cryoelectron Tomography. Proc Natl Acad Sci U S A 2012, 109 (12), 4449–4454. 10.1073/PNAS.1201333109/SUPPL_FILE/PNAS.1201333109_SI.PDF.

(10) Sibert, B. S.; Kim, J. Y.; Yang, J. E.; Wright, E. R. Micropatterning Transmission Electron Microscopy Grids to Direct Cell Positioning within Whole-Cell Cryo-Electron Tomography Workflows. J Vis Exp 2021, 2021 (175). 10.3791/62992.

(11) Nehme, E.; Freedman, D.; Gordon, R.; Ferdman, B.; Weiss, L. E.; Alalouf, O.; Naor, T.; Orange, R.; Michaeli, T.; Shechtman, Y. DeepSTORM3D: Dense 3D Localization Microscopy and PSF Design by Deep Learning. Nature Methods 2020 17:7 2020, 17 (7), 734–740. 10.1038/s41592-020-0853-5.

(12) Small, A.; Stahlheber, S. Fluorophore Localization Algorithms for Super-Resolution Microscopy. Nature Methods 2014 11:3 2014, 11 (3), 267–279. 10.1038/nmeth.2844.

(13) Juette, M. F.; Gould, T. J.; Lessard, M. D.; Mlodzianoski, M. J.; Nagpure, B. S.; Bennett, B. T.; Hess, S. T.; Bewersdorf, J. Three-Dimensional Sub–100 Nm Resolution Fluorescence Microscopy of Thick Samples. Nature Methods 2008 5:6 2008, 5 (6), 527–529. 10.1038/nmeth.1211.

(14) Ovesný, M.; Křížek, P.; Borkovec, J.; Švindrych, Z.; Hagen, G. M. ThunderSTORM: A Comprehensive ImageJ Plug-in for PALM and STORM Data Analysis and Super-Resolution Imaging. Bioinformatics 2014, 30 (16), 2389–2390. 10.1093/BIOINFORMATICS/BTU202.

(15) Pope, I.; Tanner, H.; Masia, F.; Payne, L.; Arkill, K. P.; Mantell, J.; Langbein, W.; Borri, P.; Verkade, P. Correlative Light-Electron Microscopy Using Small Gold Nanoparticles as Single Probes. Light: Science & Applications 2023 12:1 2023, 12 (1), 1–14. 10.1038/s41377-023-01115-4.

(16) Tuijtel, M. W.; Koster, A. J.; Jakobs, S.; Faas, F. G. A.; Sharp, T. H. Correlative Cryo Super-Resolution Light and Electron Microscopy on Mammalian Cells Using Fluorescent Proteins. Scientific Reports 2019 9:1 2019, 9 (1), 1–11. 10.1038/s41598-018-37728-8.

(17) Samuylov, D. K.; Purwar, P.; Szekely, G.; Paul, G. Modeling Point Spread Function in Fluorescence Microscopy with a Sparse Gaussian Mixture: Tradeoff between Accuracy and Efficiency. IEEE Transactions on Image Processing 2019, 28 (8), 3688–3702. 10.1109/TIP.2019.2898843.

(18) Li, Y.; Wu, Y.-L.; Ries, J.; Mund, M.; Hoess, P. Depth-Dependent PSF Calibration and Aberration Correction for 3D Single-Molecule Localization. Biomedical Optics Express, Vol. 10, Issue 6, p. 2708-2718 2019, 10 (6), 2708–2718. 10.1364/BOE.10.002708.

(19) Petrov, P. N.; Shechtman, Y.; Moerner, W. E.; Qian, H.; Sheetz, M. P.; Elson, E. L.; Betzig, E.; Patterson, G. H.; Sougrat, R.; Lindwasser, O. W.; Olenych, S.; Bonifacino, J. S.; Davidson, M. W.; Lippincott-Schwartz, J.; Hess, H. F. Measurement-Based Estimation of Global Pupil Functions in 3D Localization Microscopy. Optics Express, Vol. 25, Issue 7, pp. 7945-7959 2017, 25 (7), 7945–7959. 10.1364/OE.25.007945.

(20) Fölling, J.; Belov, V.; Riedel, D.; Schönle, A.; Egner, A.; Eggeling, C.; Bossi, M.; Hell, S. W. Three Dimensional Single Molecule Localization Using a Phase Retrieved Pupil Function. Optics Express, Vol. 21, Issue 24, pp. 29462-29487 2013, 21 (24), 29462–29487. 10.1364/OE.21.029462.

(21) Briegel, A.; Chen, S.; Koster, A. J.; Plitzko, J. M.; Schwartz, C. L.; Jensen, G. J. Correlated Light and Electron Cryo-Microscopy. Methods Enzymol 2010, 481 (C), 317–341. 10.1016/S0076-6879(10)81013-4.

(22) Moser, F.; Prazák, V.; Mordhorst, V.; Andrade, D. M.; Baker, L. A.; Hagen, C.; Grünewald, K.; Kaufmann, R. Cryo-SOFI Enabling Low-Dose Super-Resolution Correlative Light and Electron Cryo-Microscopy. Proc Natl Acad Sci U S A 2019, 116 (11), 4804–4809. 10.1073/PNAS.1810690116/SUPPL_FILE/PNAS.1810690116.SM05.MOV.

(23) Koning, R. I.; Celler, K.; Willemse, J.; Bos, E.; van Wezel, G. P.; Koster, A. J. Correlative Cryo-Fluorescence Light Microscopy and Cryo-Electron Tomography of Streptomyces. Methods Cell Biol 2014, 124, 217–239. 10.1016/B978-0-12-801075-4.00010-0.

(24) Schellenberger, P.; Kaufmann, R.; Siebert, C. A.; Hagen, C.; Wodrich, H.; Grünewald, K. High-Precision Correlative Fluorescence and Electron Cryo Microscopy Using Two Independent Alignment Markers. Ultramicroscopy 2014, 143, 41–51. 10.1016/J.ULTRAMIC.2013.10.011.

(25) Sartori, A.; Gatz, R.; Beck, F.; Rigort, A.; Baumeister, W.; Plitzko, J. M. Correlative Microscopy: Bridging the Gap between Fluorescence Light Microscopy and Cryo-Electron Tomography. J Struct Biol 2007, 160 (2), 135–145. 10.1016/J.JSB.2007.07.011.

(26) Wang, W.; Wang, W.; Wang, W.; Wu, B.; Wu, B.; Zhang, B.; Zhang, B.; Li, X.; Li, X.; Tan, J.; Tan, J. Correction of Refractive Index Mismatch-Induced Aberrations under Radially Polarized Illumination by Deep Learning. Optics Express, Vol. 28, Issue 18, pp. 26028-26040 2020, 28 (18), 26028–26040. 10.1364/OE.402109.

(27) Yang, J.; Vrbovská, V.; Franke, T.; Sibert, B.; Larson, M.; Coomes, T.; Rigort, A.; Mitchels, J.; Wright, E. R. Precise 3D Localization by Integrated Fluorescence Microscopy (IFLM) for Cryo-FIB-Milling and In-Situ Cryo-ET. Microscopy and Microanalysis 2023, 29 (Supplement_1), 1055–1057. 10.1093/MICMIC/OZAD067.541.

(28) Klumpe, S.; Fung, H. K. H.; Goetz, S. K.; Zagoriy, I.; Hampoelz, B.; Zhang, X.; Erdmann, P. S.; Baumbach, J.; Müller, C. W.; Beck, M.; Plitzko, J. M.; Mahamid, J. A Modular Platform for Automated Cryo-FIB Workflows. Elife 2021, 10, e70506. 10.7554/ELIFE.70506.

(29) Zanetti-Domingues, L. C.; Hirsch, M.; Wang, L.; Eastwood, T. A.; Baker, K.; Mulvihill, D. P.; Radford, S.; Horne, J.; White, P.; Bateman, B. Toward Quantitative Super-Resolution Methods for Cryo-CLEM. Methods Cell Biol 2024, 187, 249–292. 10.1016/BS.MCB.2024.02.028.

(30) Faul, N.; Chen, S. Y.; Lamberz, C.; Bruckner, M.; Dienemann, C.; Burg, T. P. Cryo-ICLEM: Cryo Correlative Light and Electron Microscopy with Immersion Objectives. J Struct Biol 2025, 217 (1), 108179. 10.1016/J.JSB.2025.108179.

(31) Zhang, P. Advances in Cryo-Electron Tomography and Subtomogram Averaging and Classification. Curr Opin Struct Biol 2019, 58, 249–258. 10.1016/J.SBI.2019.05.021.

(32) Diel, E. E.; Lichtman, J. W.; Richardson, D. S. Tutorial: Avoiding and Correcting Sample-Induced Spherical Aberration Artifacts in 3D Fluorescence Microscopy. Nature Protocols 2020 15:9 2020, 15 (9), 2773–2784. 10.1038/s41596-020-0360-2.

(33) de Beer, M.; Daviran, D.; Roverts, R.; Rutten, L.; Macías-Sánchez, E.; Metz, J. R.; Sommerdijk, N.; Akiva, A. Precise Targeting for 3D Cryo-Correlative Light and Electron Microscopy Volume Imaging of Tissues Using a FinderTOP. Communications Biology 2023 6:1 2023, 6 (1), 1–9. 10.1038/s42003-023-04887-y.

(34) Fukuda, Y.; Stapleton, K.; Kato, T. Progress in Spatial Resolution of Structural Analysis by Cryo-EM. Microscopy 2023, 72 (2), 135–143. 10.1093/JMICRO/DFAC053.

(35) Cam, L. Le. Maximum Likelihood: An Introduction. Int Stat Rev 1990, 58 (2), 153. 10.2307/1403464.

(36) Liu, X.; Kuang, C.; Huang, Y.; Li, H. Method of Super-Resolution Based on Array Detection and Maximum-Likelihood Estimation. Applied Optics, Vol. 55, Issue 35, pp. 9925-9931 2016, 55 (35), 9925–9931. 10.1364/AO.55.009925.

(37) Silva, D. E.; Wang, J. Y. Wave-Front Interpretation with Zernike Polynomials. Applied Optics, Vol. 19, Issue 9, pp. 1510-1518 1980, 19 (9), 1510–1518. 10.1364/AO.19.001510.

(38) Lakshminarayanan, V.; Fleck, A. Zernike Polynomials: A Guide. J Mod Opt 2011, 58 (7), 545–561. 10.1080/09500340.2011.554896.

(39) Kaufmann, R.; Hagen, C.; Grünewald, K. Fluorescence Cryo-Microscopy: Current Challenges and Prospects. Curr Opin Chem Biol 2014, 20 (100), 86–91. 10.1016/J.CBPA.2014.05.007.

(40) Zheng, G.; Ou, X.; Horstmeyer, R.; Yang, C.; Brady, D. J.; Gehm, M. E.; Stack, R. A.; Marks, D. L.; Kittle, D. S.; Golish, D. R.; Vera, E. M.; Feller, S. D. Characterization of Spatially Varying Aberrations for Wide Field-of-View Microscopy. Optics Express, Vol. 21, Issue 13, pp. 15131-15143 2013, 21 (13), 15131–15143. 10.1364/OE.21.015131.

(41) Liu, S.; Chen, J.; Hellgoth, J.; Müller, L. R.; Ferdman, B.; Karras, C.; Xiao, D.; Lidke, K. A.; Heintzmann, R.; Shechtman, Y.; Li, Y.; Ries, J. Universal Inverse Modeling of Point Spread Functions for SMLM Localization and Microscope Characterization. Nature Methods 2024 21:6 2024, 21 (6), 1082–1093. 10.1038/s41592-024-02282-x.

