## Supplementary Notes for "Systematic Characterization of Optical Aberrations Reveals Cryo-FLM Localization Fidelity"

**This PDF file includes:**

Supplementary Figures 1 to 15

Supplementary Tables 1 to 2

Supplementary Notes

References

**Other supplementary materials for this manuscript include the following:**

Supplementary Software 1: Software package for Phase Retrieval.

Supplementary Software 2: Software package for Biplane Dataset Simulation.

Supplementary Software 3: Software package for Biplane 3D Localization.

#### Supplementary Figures

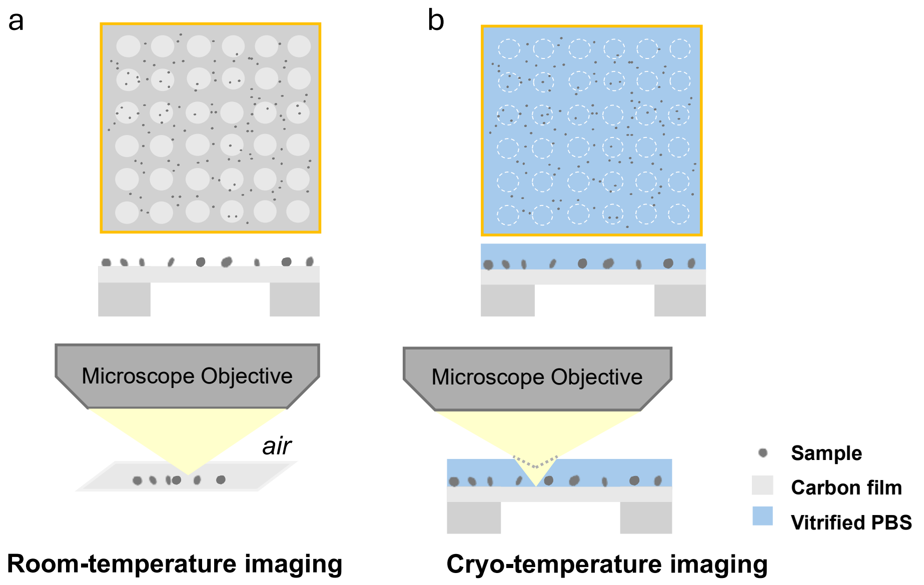

**Supplementary Fig. 1. Sample and optical path schematic for joint room-temperature and cryo-temperature fluorescence microscopy imaging.**

**(a)** Room-temperature (RT) imaging configuration. TetraSpeck beads are directly adhered to the carbon film, and imaging is performed with an air objective at ambient temperature (refractive index, RI ≈ 1.00). In this setup, there is no refractive index mismatch along the optical path. **(b)** Cryo-temperature (cryoT) imaging configuration. TetraSpeck beads are embedded in vitrified phosphate-buffered saline (PBS, RI ≈ 1.33) while remaining on the carbon film. The objective is immersed in nitrogen vapor, creating a refractive index mismatch between the cryogenic immersion medium and the sample. This mismatch bends the light and introduces additional optical aberrations. A carbon-coating step is performed prior to sample preparation to establish a thin layer of carbon.

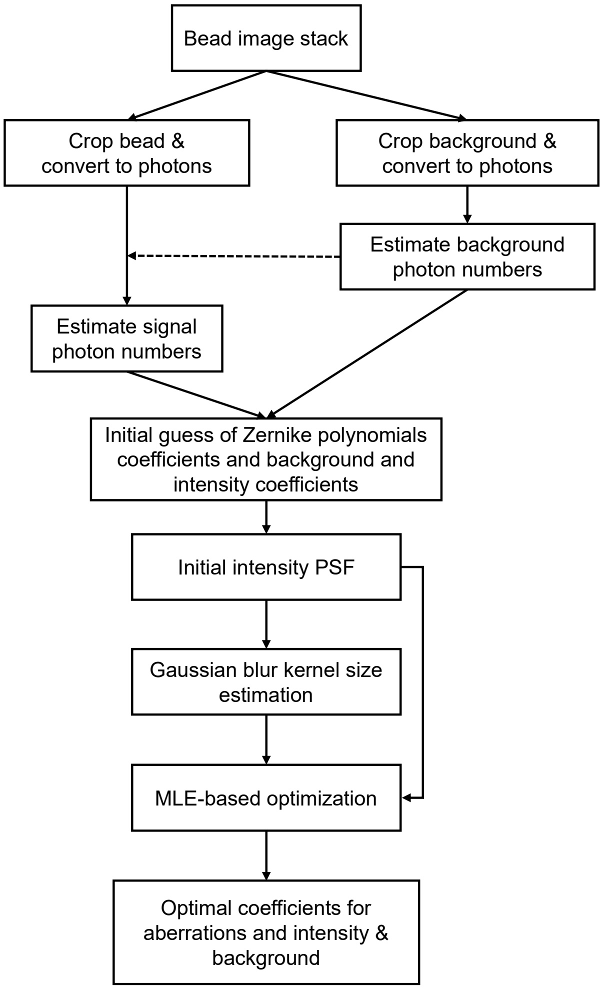

**Supplementary Fig. 2. Workflow for data preprocessing and MLE-based phase retrieval.**

The input is a stack of TetraSpeck bead images acquired at different axial positions (e.g., −2 to +2 μm, in 200 nm steps). Subregions (e.g., 32×32 pixels) are cropped around a bead and a nearby background area. Initial background and signal photon numbers are estimated from these subregions (**Supplementary Notes 2.4**), and the initial Zernike polynomial coefficients are randomly generated. An initial intensity PSF is then constructed and used to calculate a Gaussian kernel to compensate for the size of the beads (**Supplementary Notes 2.5**). Finally, a maximum likelihood estimation (MLE)-based phase retrieval algorithm is applied to jointly optimize the Zernike coefficients, background, and intensity.

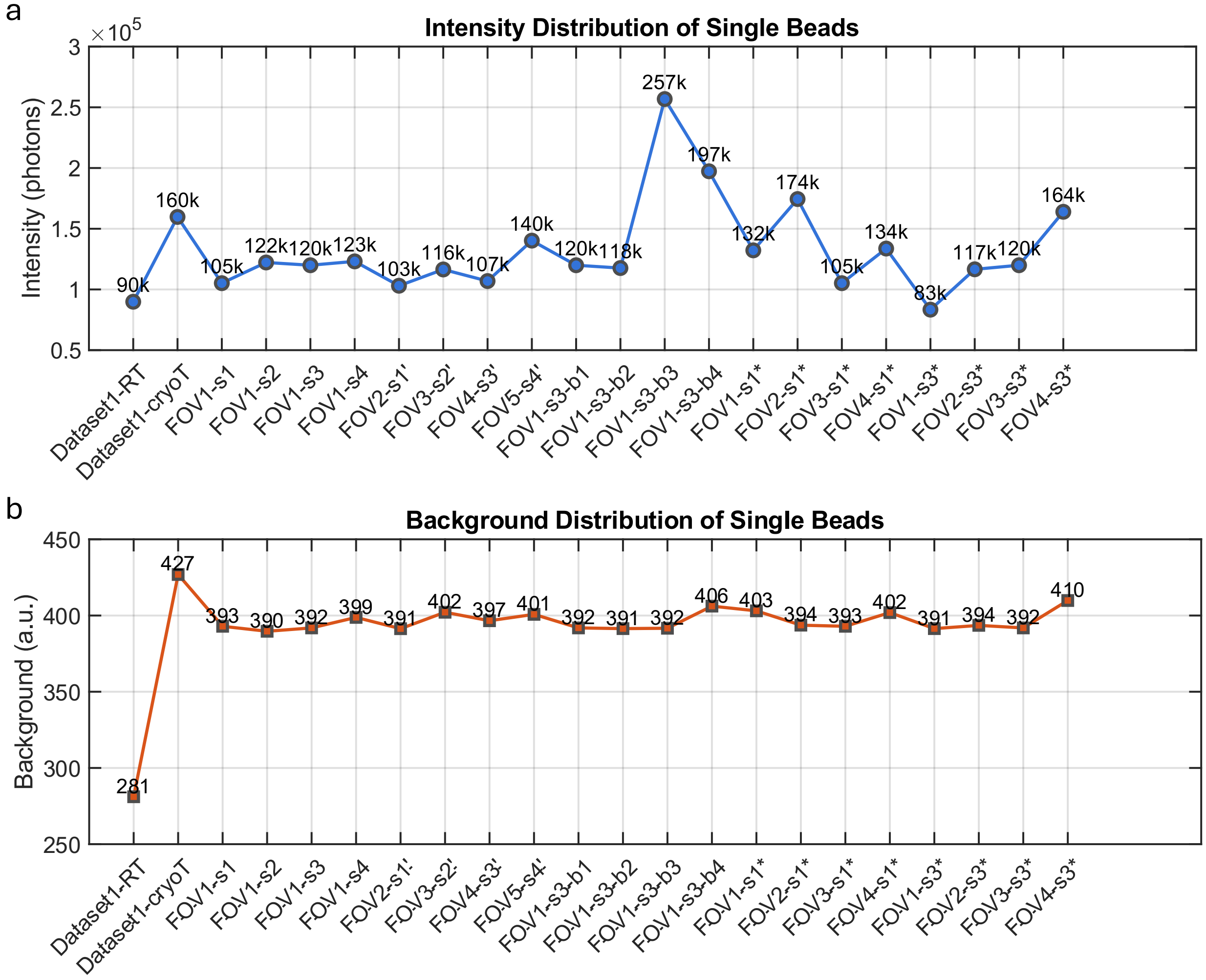

**Supplementary Fig. 3. Summary of signal and background photon counts across all experimental datasets.**

We quantified the signal **(a)** and background **(b)** photon counts from all bead image stacks (32×32 pixel subregions) used in the experiments. The background value is calculated as the average across all pixels in the background region, while the signal intensity is averaged over all axial frames for each bead (**Supplementary Notes 2.4**). The average background is approximately 400 photons, and the signal intensity ranges from ~1 × 10⁵ to ~1.5 × 10⁵ photons.

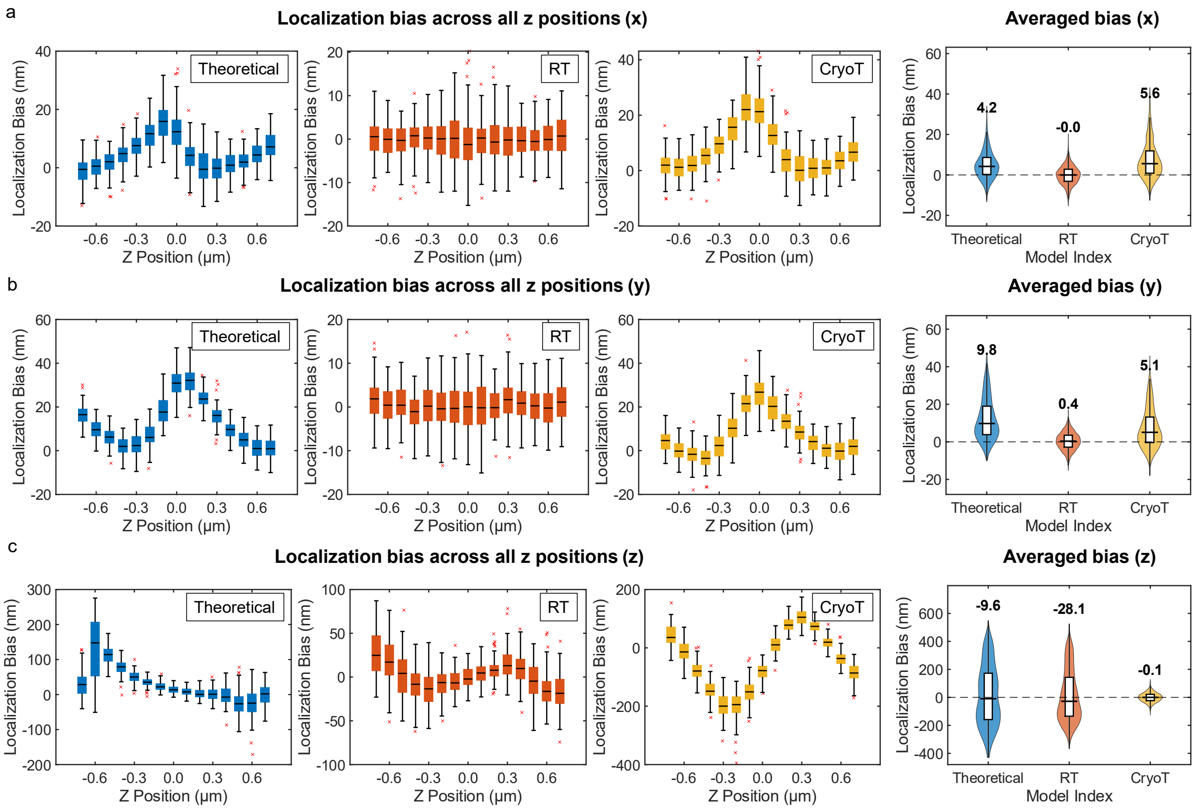

**Supplementary Fig. 4. Localization bias analysis of room-temperature FLM.**

Simulated datasets were generated using the room-temperature (RT) PSF model across axial positions from −0.7 μm to +0.7 μm in 100 nm increments, with 100 frames per z-position. Simulations were performed with a signal photon count of $1 \times 10^4$ and a background photon count of 400. Three PSF models were used for 3D localization: a theoretical model (flat pupil), the RT PSF model, and the cryogenic-temperature (CryoT) PSF model. **(a–c)** Localization bias in x, y, and z directions, respectively, across the axial range and bias distributions aggregated over all positions are also shown.

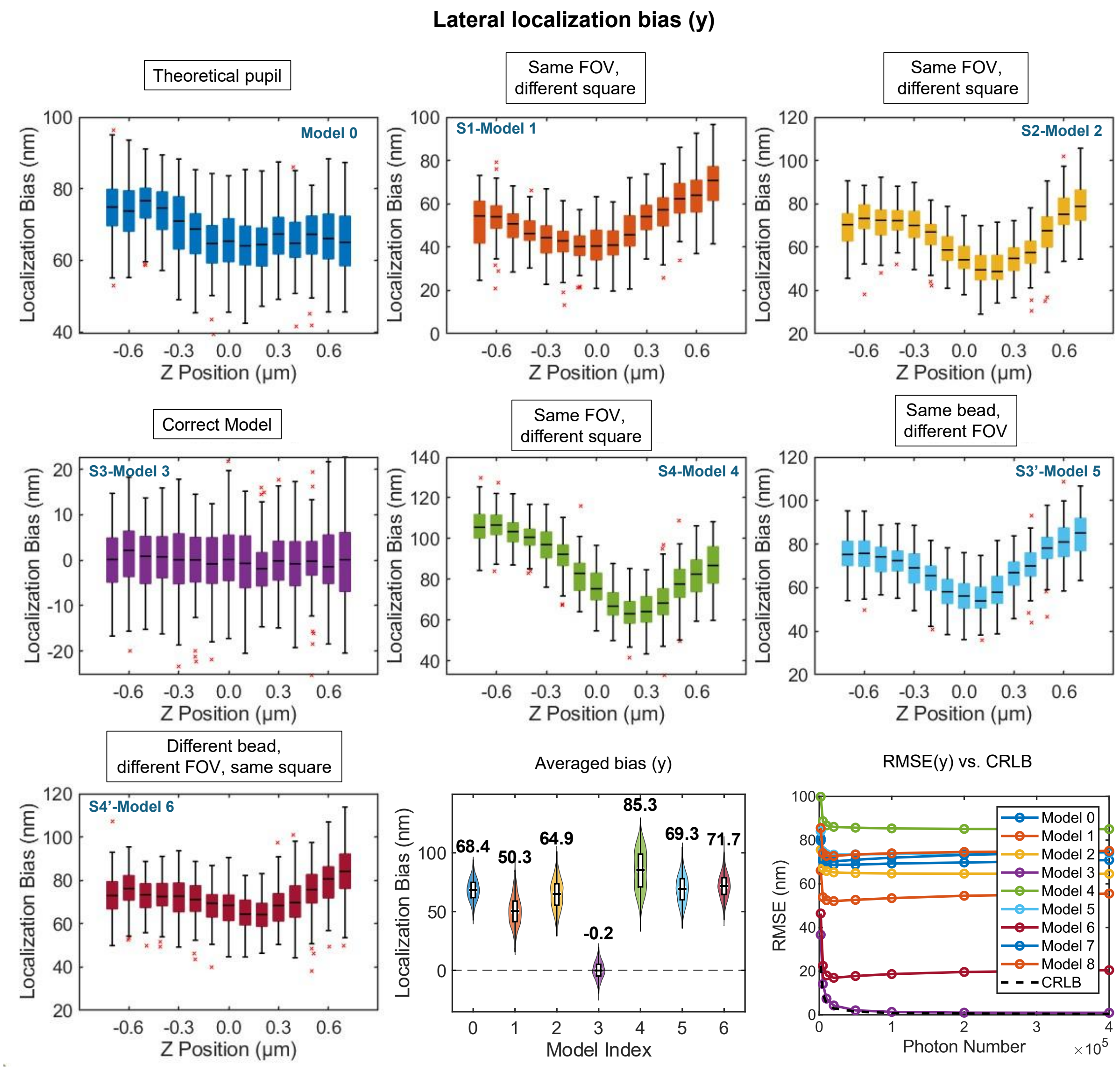

**Supplementary Fig. 5. Localization bias analysis and RMSE (y) for FOV-dependent PSF models.** Test data were generated using the PSF model retrieved from a bead in FOV1-S3 (**Fig. 3a**), spanning axial positions from –0.7 µm to +0.7 µm in 100 nm steps (100 frames per z-position). Simulations were performed with a signal photon count of $1 \times 10^4$ and a background photon count of 400. Seven PSF models were used for 3D localization: Model 0 (theoretical pupil function), Models 1–4 (experimental PSFs from beads in FOV1-S1 to S4), and Models 5–6 (PSFs from S3′ and S4′, representing system-induced and sample-induced aberrations, respectively). Lateral (y) localization bias as a function of axial position, and corresponding average localization bias across all z positions, bias distributions aggregated over all positions and localization RMSE compared to the CRLB across varying photon counts are shown.

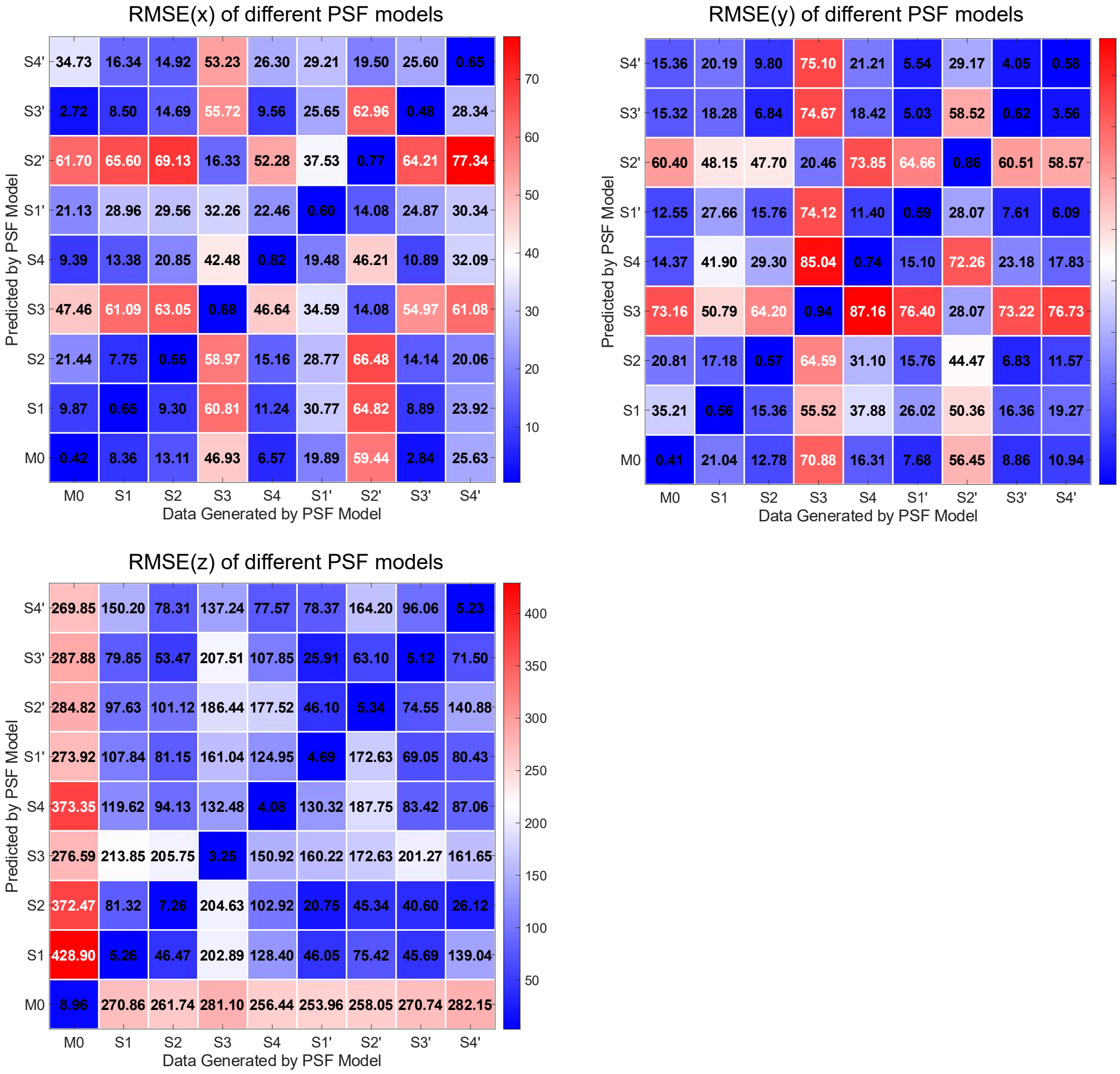

**Supplementary Fig. 6. Localization RMSE analysis of FOV-dependent PSF models at high photon counts (4 × $10^5$).**

Simulated datasets generated using the theoretical PSF model and PSF models retrieved from FOV1–FOV5 (**Fig. 3**), spanning axial positions from −0.7 µm to +0.7 µm in 100 nm steps, with 100 frames per z-position. Simulations were performed with a signal photon count of 4 × $10^5$ and a background photon count of 400, approximating the performance limit at infinite photon numbers. These models were used for 3D localization, and the RMSE in x-, y- and z-localization was calculated.

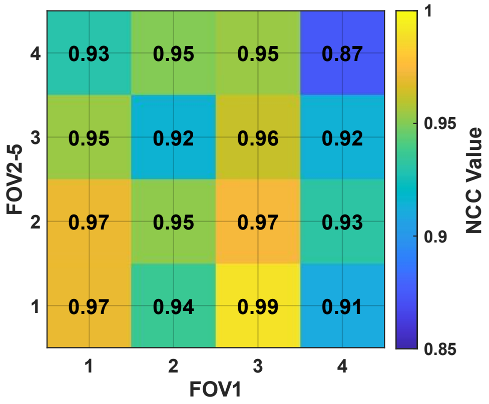

**Supplementary Fig. 7. Partial normalized cross correlation (PNCC) matrix for FOV-dependent PSF models.**

The diagonal values represent the PSF dissimilarity arising from system-induced aberration differences.

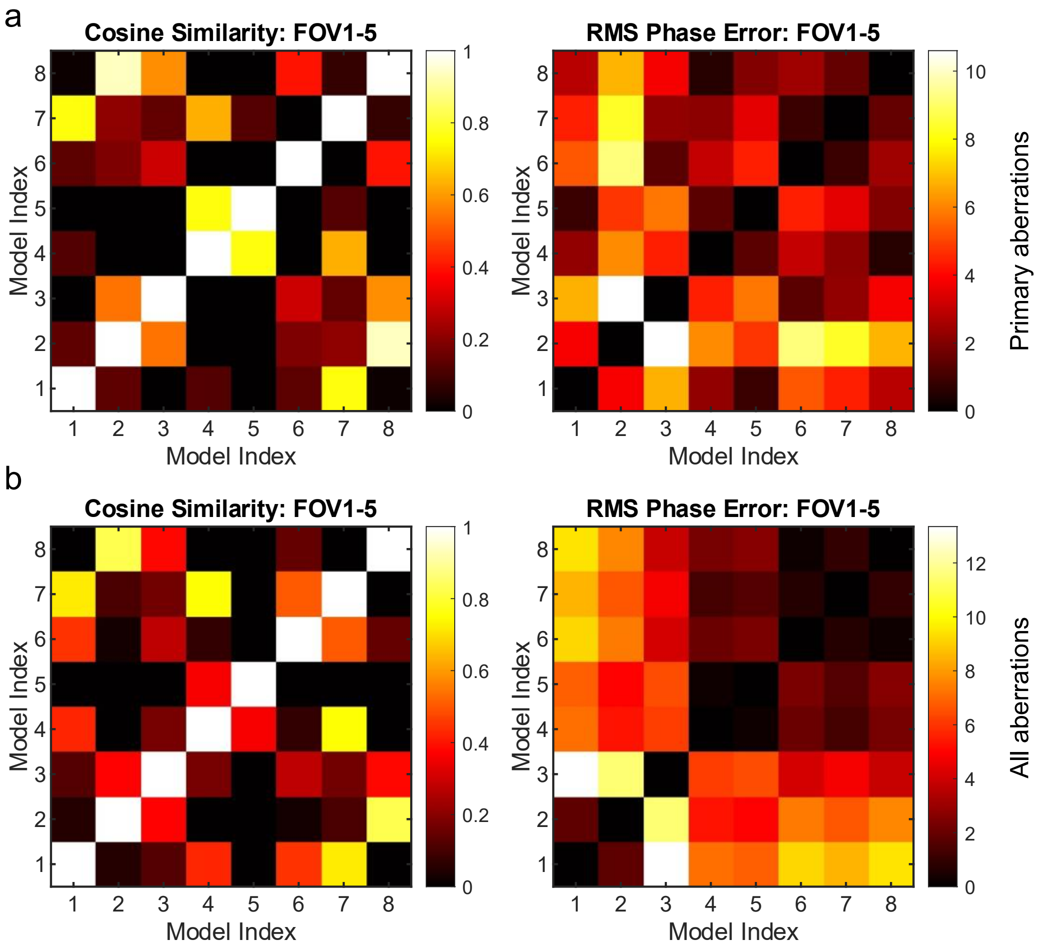

**Supplementary Fig. 8. The comparison of the FOV-dependent aberrations.**

**(a)** Primary aberrations (Zernike polynomials of Noll orders 5th to 11th). Cosine similarity matrix of Zernike coefficients and RMS phase error matrix among FOV-dependent PSF models retrieved from FOV1–FOV5 (see **Fig. 3**). **(b)** Full aberration set (Zernike polynomials of Noll orders 5th to 36th). Cosine similarity and RMS phase error matrices among the same FOV-dependent PSF models.

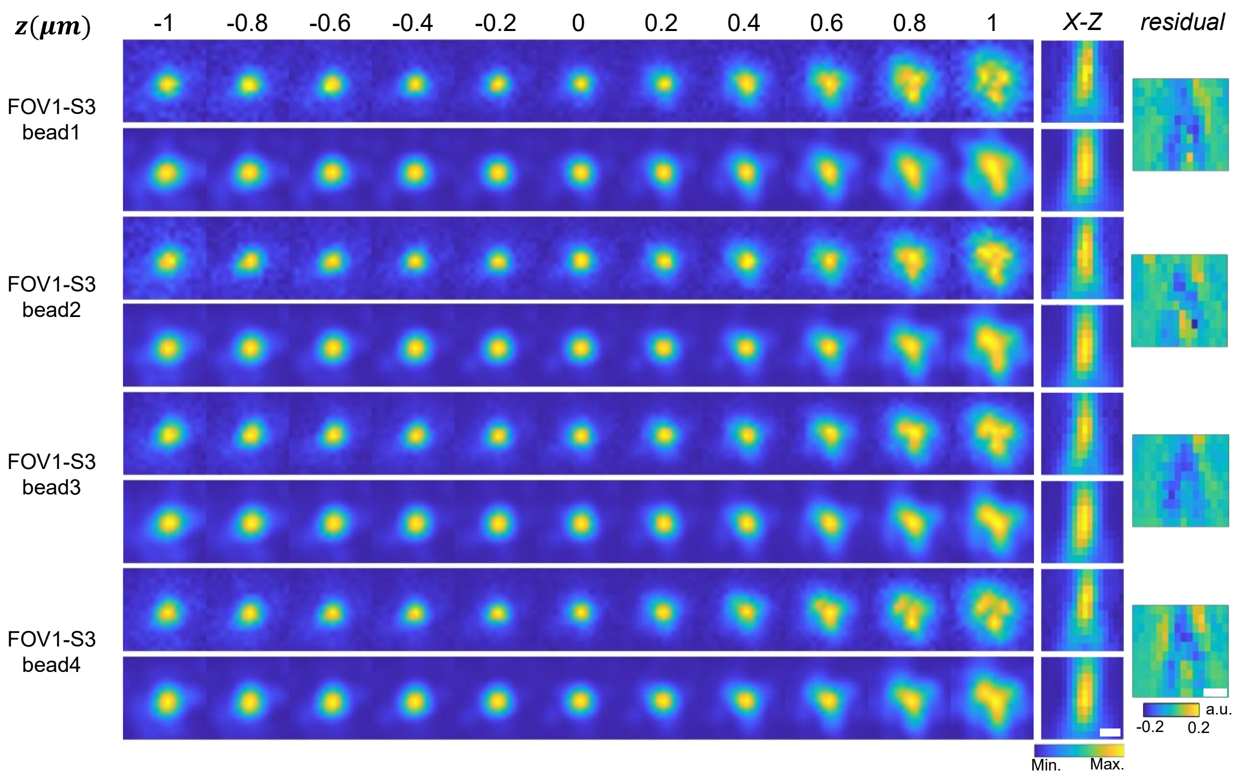

**Supplementary Fig. 9**. **Experimental and phase-retrieved PSFs of four beads in same TEM square (FOV1-S3).**

xy projections of the experimental and phase-retrieved PSFs across a z-range from –1 μm to 1 μm for beads in FOV1-S3 (**Fig. 3**). Corresponding xz views and residuals between the experimental and reconstructed PSFs are also presented. Scale bar: 500 nm.

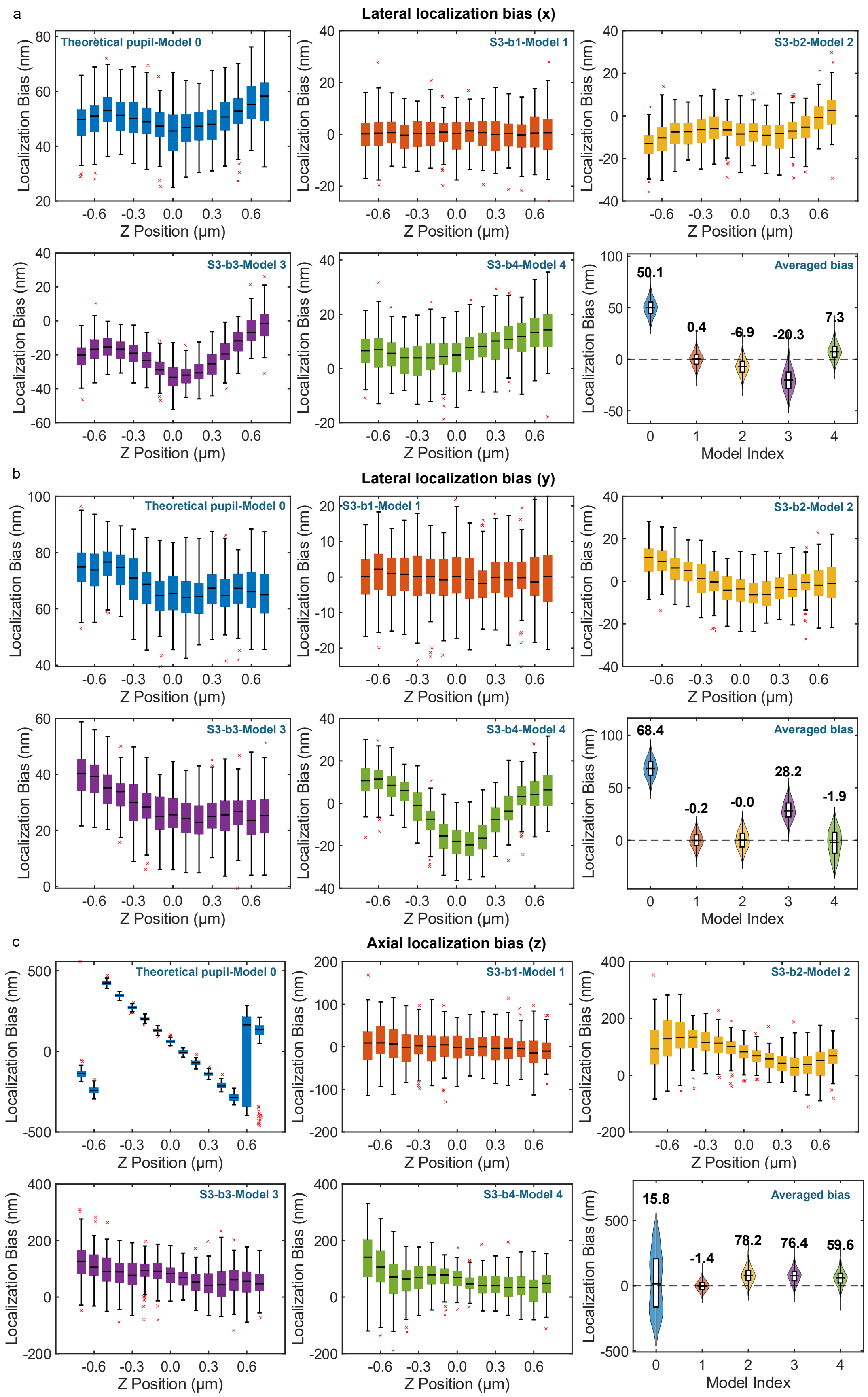
a
Lateral localization bias (x)
Theoretical pupil-Model 0
S3-b1-Model 1
S3-b2-Model 2
S3-b3-Model 3
S3-b4-Model 4
Averaged bias
50.1
0.4
-6.9
-20.3
7.3
Localization Bias (nm)
Z Position (μm)
Model Index
b
Lateral localization bias (y)
Theoretical pupil-Model 0
S3-b1-Model 1
S3-b2-Model 2
S3-b3-Model 3
S3-b4-Model 4
Averaged bias
68.4
-0.2
-0.0
28.2
-1.9
Localization Bias (nm)
Z Position (μm)
Model Index
c
Axial localization bias (z)
Theoretical pupil-Model 0
S3-b1-Model 1
S3-b2-Model 2
S3-b3-Model 3
S3-b4-Model 4
Averaged bias
15.8
-1.4
78.2
76.4
59.6
Localization Bias (nm)
Z Position (μm)
Model Index

**Supplementary Fig. 10**. **Localization bias analysis for square-dependent PSF models.**

Test data were generated using the PSF model retrieved from a bead in FOV1-S3-Bead1 (**Fig. 3a**), spanning axial positions from –0.7 μm to +0.7 μm in 100 nm steps (100 frames per z-position). Simulations were performed with a signal photon count of $1 \times 10^4$ and a background photon count of 400. Five PSF models were used for 3D localization: Model 0 (theoretical pupil function), Models 1–4 (experimental PSFs from beads in FOV1-S3). **(a-c)** localization bias as a function of axial position, and corresponding bias distributions aggregated over all positions for x, y, z.

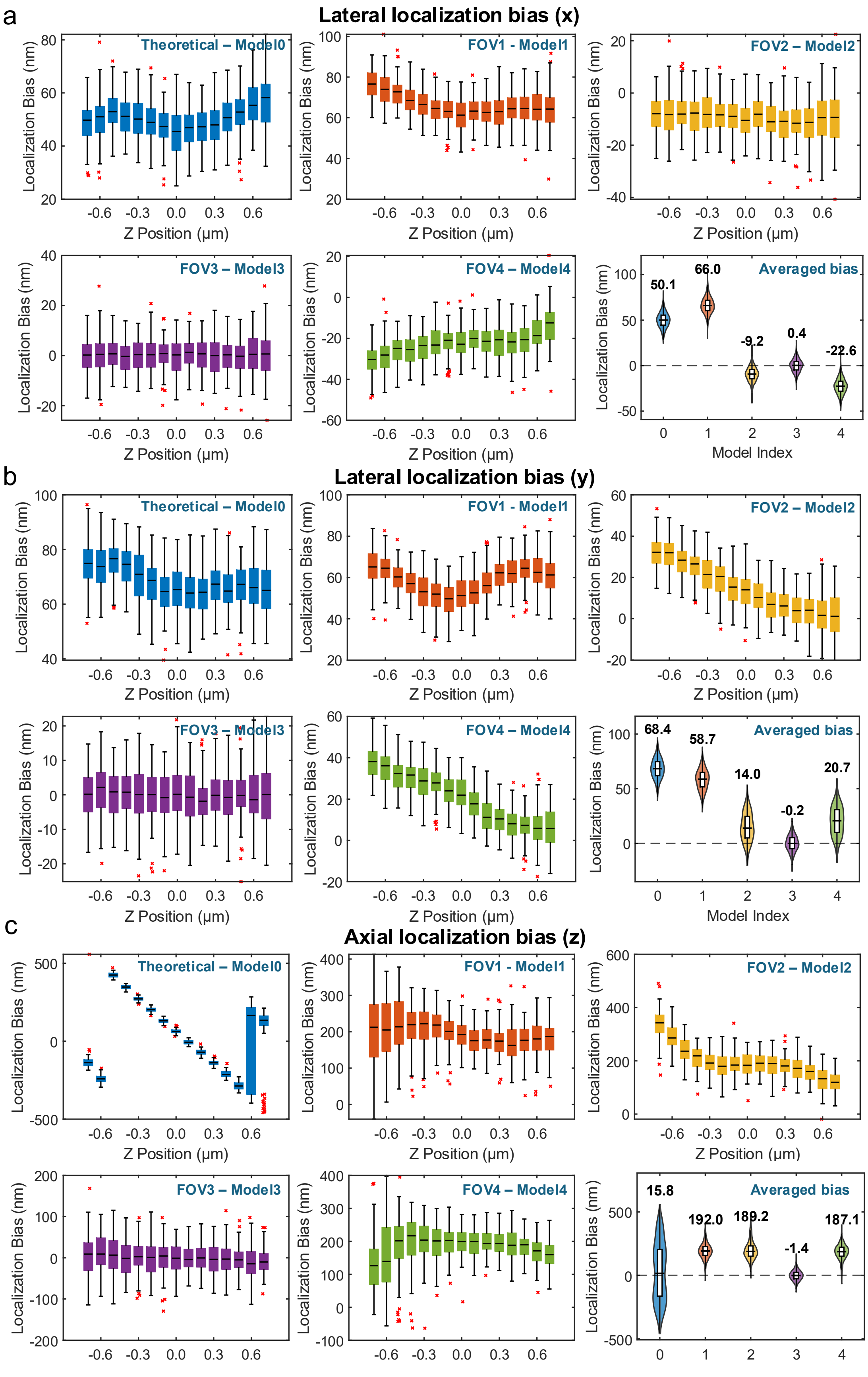
a
Lateral localization bias (x)
Theoretical – Model0
FOV1 - Model1
FOV2 – Model2
FOV3 – Model3
FOV4 – Model4
Averaged bias
50.1
66.0
-9.2
0.4
-22.6
Localization Bias (nm)
Z Position (μm)
Model Index
b
Lateral localization bias (y)
Theoretical – Model0
FOV1 - Model1
FOV2 – Model2
FOV3 – Model3
FOV4 – Model4
Averaged bias
68.4
58.7
14.0
-0.2
20.7
Localization Bias (nm)
Z Position (μm)
Model Index
c
Axial localization bias (z)
Theoretical – Model0
FOV1 - Model1
FOV2 – Model2
FOV3 – Model3
FOV4 – Model4
Averaged bias
15.8
192.0
189.2
-1.4
187.1
Localization Bias (nm)
Z Position (μm)

**Supplementary Fig. 11**. **Localization bias analysis for grid-dependent PSF models.**

Test data were generated using the PSF model retrieved from a bead in FOV3 (**Fig. 5a**), spanning axial positions from –0.7 μm to +0.7 μm in 100 nm steps (100 frames per z-position). Simulations were performed with a signal photon count of $1 \times 10^4$ and a background photon count of 400. Five PSF models were used for 3D localization: Model 0 (theoretical pupil function), Models 1–4 (experimental PSFs from beads in FOV1-4 in **Fig. 5a**). **(a-c)** localization bias as a function of axial position, and corresponding bias distributions aggregated over all positions for x, y, z

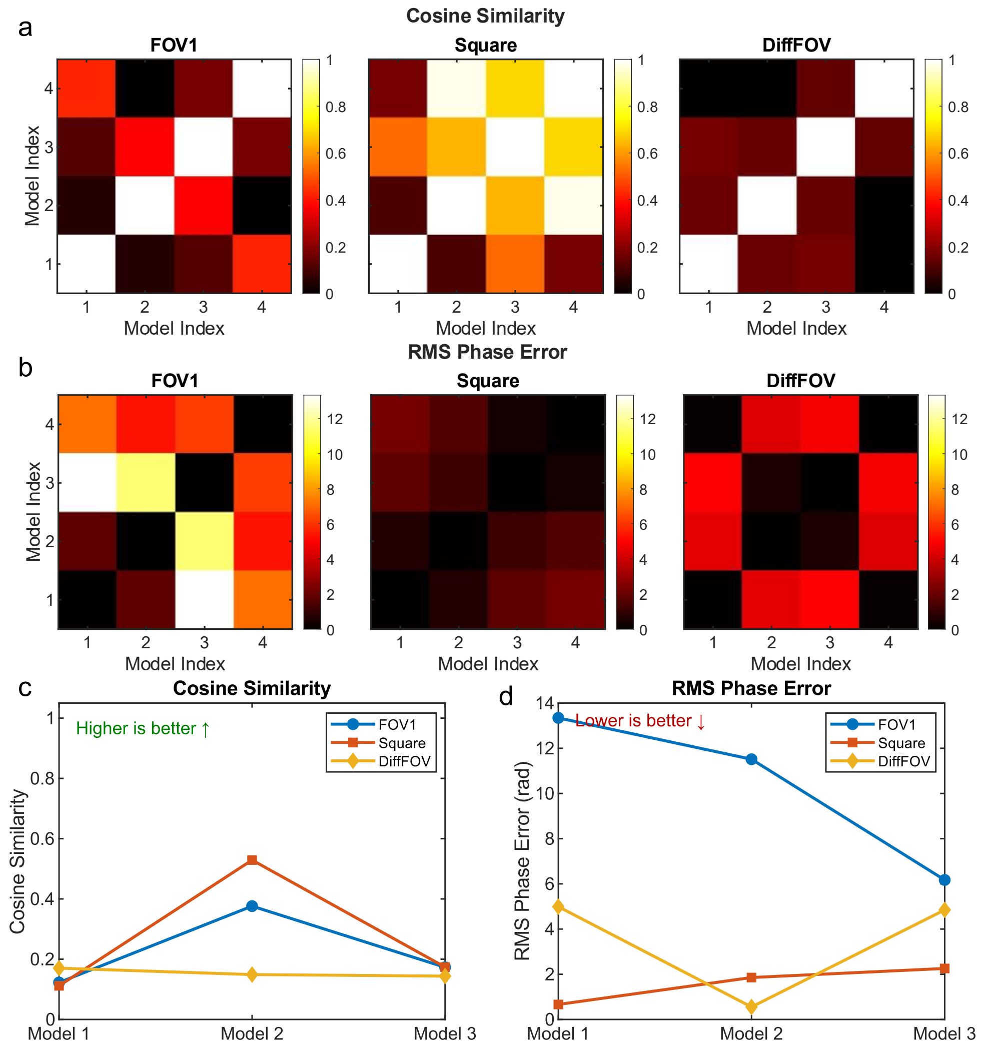

**Supplementary Fig. 12. Comparison of FOV-dependent, square-dependent, and grid-dependent full-set aberrations.**

The full aberration set includes Zernike polynomials of Noll orders 5$^{th}$ to 36$^{th}$. **(a)** Cosine similarity matrix of Zernike coefficients within FOV-dependent (FOV1), square-dependent (Square), and grid-dependent (DiffFOV) aberration groups. **(b)** RMS phase error matrix for the same three aberration groups. **(c)** Direct comparison of cosine similarity and RMS phase error for one representative PSF model in each group. The same PSF model is used across all three comparisons—S3 in FOV1 (FOV-dependent, **Fig. 3a**), Bead 1 in Square1 (square-dependent, **Fig. 3a**), and FOV3 in a different FOV (grid-dependent, **Fig. 5a**). Comparisons are also made with three other models from the same respective groups to evaluate intra-group variability.

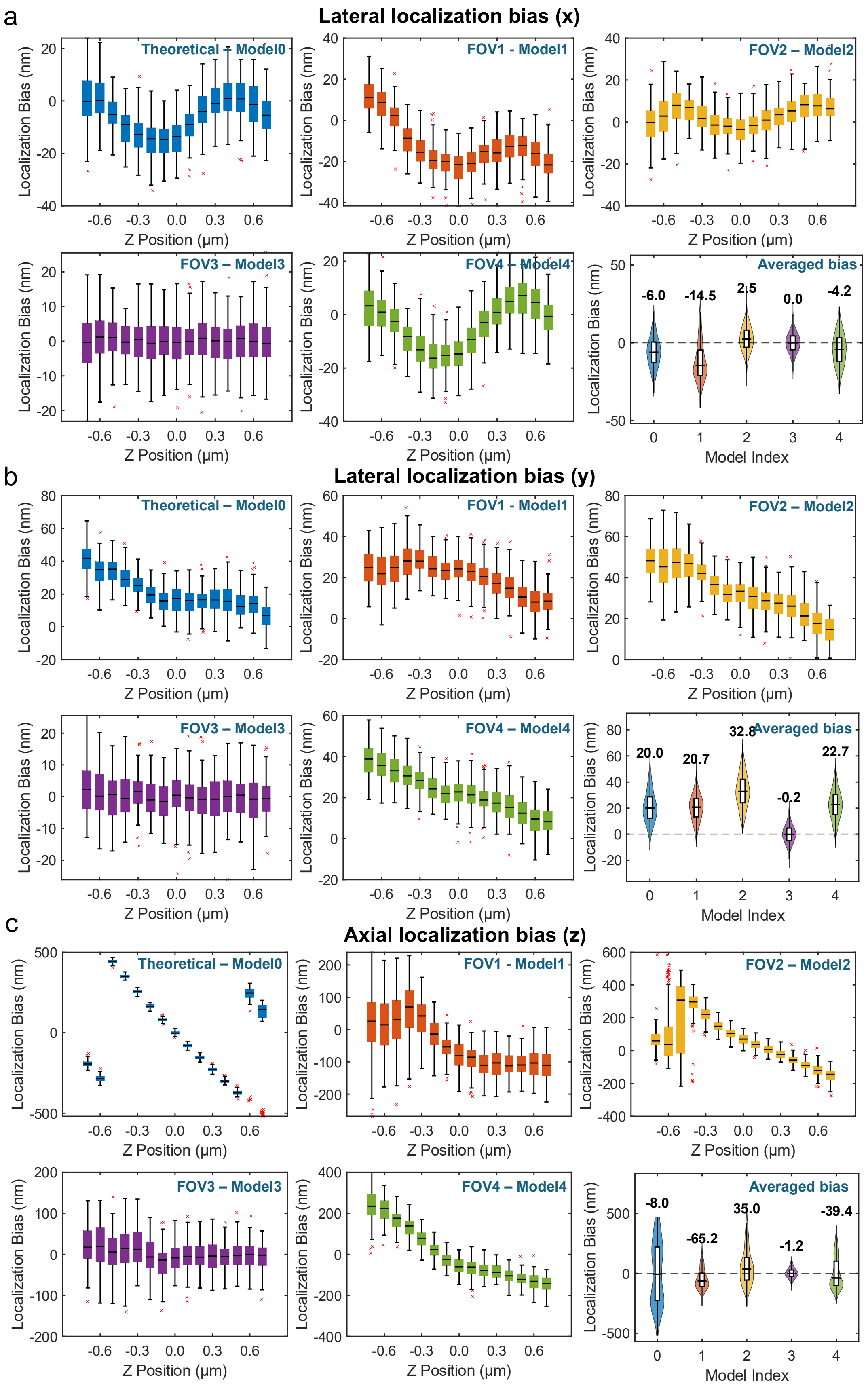

a
Lateral localization bias (x)
Theoretical – Model0
FOV1 - Model1
FOV2 – Model2
FOV3 – Model3
FOV4 – Model4
Averaged bias
-6.0
-14.5
2.5
0.0
-4.2
Localization Bias (nm)
Z Position (μm)
Model Index
b
Lateral localization bias (y)
Theoretical – Model0
FOV1 - Model1
FOV2 – Model2
FOV3 – Model3
FOV4 – Model4
Averaged bias
20.0
20.7
32.8
-0.2
22.7
Localization Bias (nm)
Z Position (μm)
Model Index
c
Axial localization bias (z)
Theoretical – Model0
FOV1 - Model1
FOV2 – Model2
FOV3 – Model3
FOV4 – Model4
Averaged bias
-8.0
-65.2
35.0
-1.2
-39.4
Localization Bias (nm)
Z Position (μm)
Model Index

**Supplementary Fig. 13**. **Localization bias analysis for an additional case of grid-dependent PSF aberrations.**

Test data were generated using the PSF model retrieved from a bead in FOV3 (**Supplementary Fig. SS9**), spanning axial positions from –0.7 µm to +0.7 µm in 100 nm steps (100 frames per z-position). Simulations were performed with a signal photon count of $1 \times 10^4$ and a background photon count of 400. Five PSF models were used for 3D localization: Model 0 (theoretical pupil function), Models 1–4 (experimental PSFs from beads in FOV1-4). **(a-c)** localization bias as a function of axial position, and corresponding bias distributions aggregated over all positions for x, y, z.

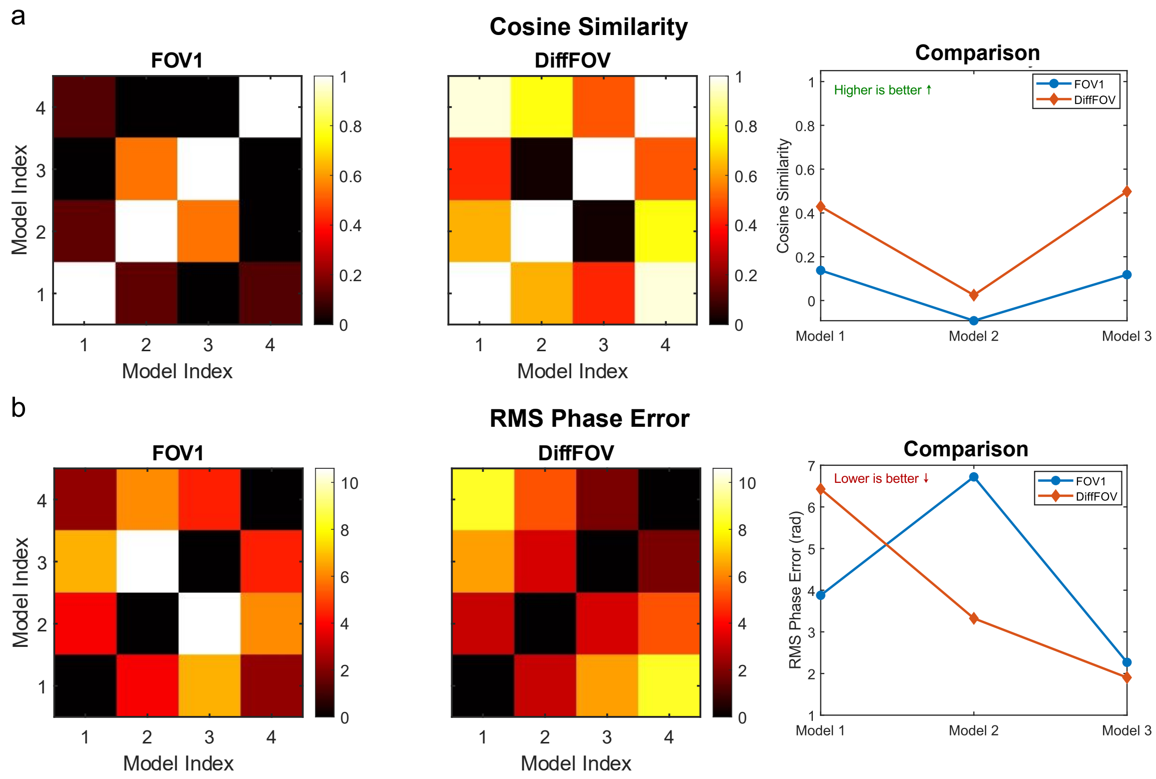

**Supplementary Fig. 14. Another case study comparing FOV-dependent (FOV1-S1) and grid-dependent (DiffFOV) primary aberrations.**

The primary aberration set includes Zernike polynomials of Noll orders 5$^{th}$ to 11$^{th}$. **(a)** Cosine similarity matrix of Zernike coefficients within FOV-dependent (FOV1, FOV1-S1 **in Fig. 3**), and grid-dependent (DiffFOV, FOV1 in **Supplementary Fig. SS9**) aberration groups. **(b)** RMS phase error corresponding to each aberration condition. Direct comparison of cosine similarity and RMS phase error for one representative PSF model in each group. The same PSF model is used across both comparisons—S1 in FOV1 (FOV-dependent), and FOV3 in a different FOV (grid-dependent). Comparisons are also made with three other models from the same respective groups to evaluate intra-group variability.

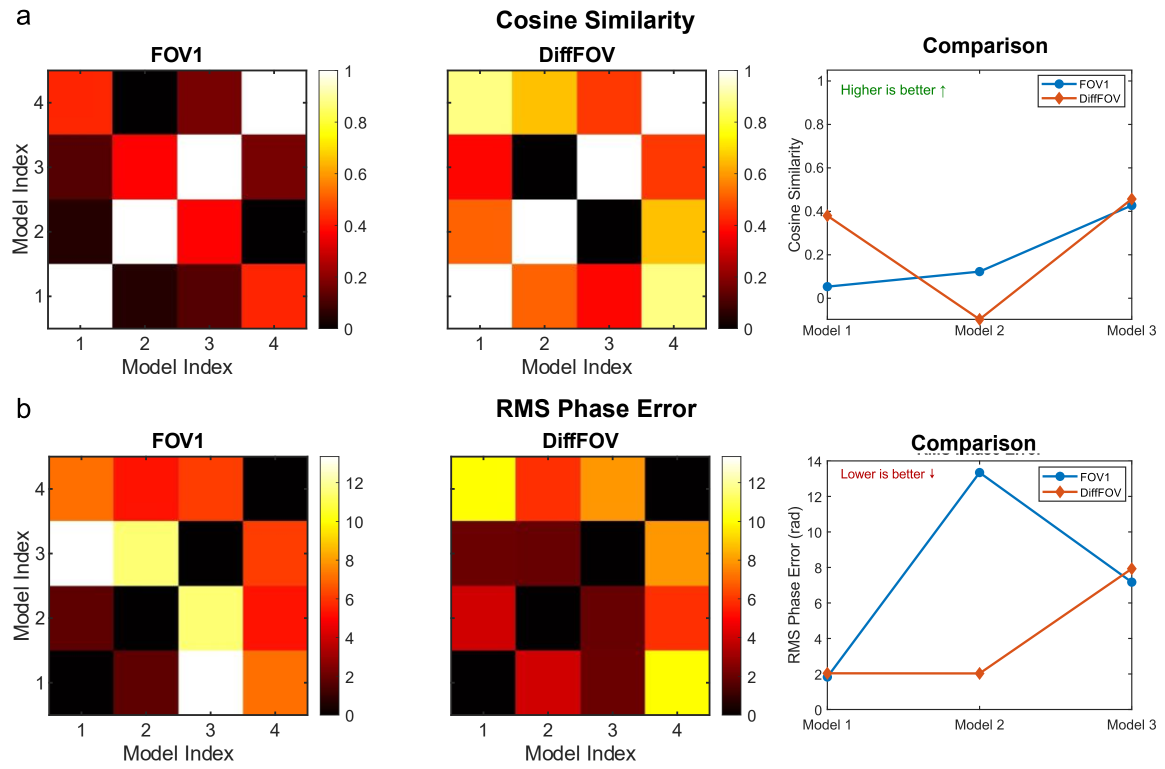

**Supplementary Fig. 15. Another case study comparing FOV-dependent (FOV1-S1) and grid-dependent (DiffFOV) full-set aberrations.**

The full aberration set includes Zernike polynomials of Noll orders 5th to 36th. **(a)** Cosine similarity matrix of Zernike coefficients within FOV-dependent (FOV1, FOV1-S1 **in Fig. 3**), and grid-dependent (DiffFOV, FOV1 in **Supplementary SS9**) aberration groups. **(b)** RMS phase error corresponding to each aberration condition. Direct comparison of cosine similarity and RMS phase error for one representative PSF model in each group. The same PSF model is used across both comparisons—S1 in FOV1 (FOV-dependent), and FOV3 in a different FOV (grid-dependent). Comparisons are also made with three other models from the same respective groups to evaluate intra-group variability.

**Supplementary Table 1. The localization RMSE for theoretical, RT and cryoT PSF models on cryoT dataset.**

| | Localization model | Photon number=$\mathbf{1 \times 10^4}$ | Photon number=$\mathbf{4 \times 10^5}$ |
|---|---|---|---|
| x | Theoretical | 9.80 | 7.40 |
| | RT | 16.75 | 15.11 |
| | CryoT | 6.62 | 0.70 |
| y | Theoretical | 11.45 | 11.63 |
| | RT | 19.70 | 10.45 |
| | CryoT | 6.44 | 0.73 |
| z | Theoretical | 209.21 | 271.48 |
| | RT | 182.13 | 227.72 |
| | CryoT | 33.25 | 3.40 |

**Supplementary Table 2. The localization RMSE for theoretical, RT and cryoT PSF models on RT dataset.**

| | Localization model | Photon number=$\mathbf{1} \times \mathbf{10^4}$ | Photon number=$\mathbf{4} \times \mathbf{10^5}$ |
|---|---|---|---|
| x | Theoretical | 8.41 | 5.66 |
| | RT | 4.59 | 0.45 |
| | CryoT | 11.56 | 10.32 |
| y | Theoretical | 16.44 | 10.60 |
| | RT | 4.71 | 0.50 |
| | CryoT | 12.64 | 7.41 |
| z | Theoretical | 62.86 | 44.46 |
| | RT | 22.59 | 5.75 |
| | CryoT | 111.35 | 124.86 |

### Supplementary Notes

#### 1. Supplementary demonstrations

##### 1.1. Room-temperature fluorescent imaging system

**Room-temperature fluorescence imaging systems exhibit greater tolerance to theoretical PSF models due to weaker aberrations.** Although room-temperature (RT) fluorescence imaging has been extensively studied[1,2], we performed a localization analysis under RT conditions to establish a benchmark for cryogenic fluorescence light microscopy (cryo-FLM), as theoretical models (flat pupils) are often used for RT localization when accurate PSFs are unavailable. The simulated biplane data were generated using the RT PSF model, and three-dimensional (3D) localizations were performed using three PSF models: the theoretical pupil model, the RT PSF model, and the cryo-temperature (CryoT) PSF model (**Fig. 2**). As expected, the matched RT PSF model yielded the highest localization accuracy, with lateral root mean square errors (RMSE) below 1 nm and axial errors under 10 nm at photon counts of $4 \times 10^5$ (**Supplementary Table 2**). Notably, the theoretical pupil model introduced smaller localization errors under RT conditions, approximately 5–10 nm laterally and ~50 nm axially, compared to its performance in cryo-FLM, where mismatches led to significantly higher errors (~10 nm lateral and ~270 nm axial; **Supplementary Table 1, Extended Data Fig. 2**). This difference arises despite using the same sample and imaging system, suggesting that while system-specific aberrations remain largely consistent, additional complex aberrations emerge under cryogenic conditions. These may stem from sample-induced effects or subtle changes in optical alignment or refractive index mismatches at low temperatures. Crucially, we found that localization errors caused by PSF model mismatches persisted even at very high photon counts ($4 \times 10^5$), emphasizing that such systematic errors cannot be compensated for by increasing photon statistics alone.

Taken together, these results suggest that cryogenic conditions introduce substantial and complex aberrations, both sample- and system-related, that degrade localization accuracy far more severely than in RT FLM. These aberrations must be carefully characterized and corrected to achieve optimal performance in cryo-FLM.

#### 1.2. More results related to FOV- and square-dependent aberrations

**The PSF profiles show consistently larger FWHM values under cryogenic conditions.** Cryo-temperature PSFs exhibit substantially larger full width at half maximum (FWHM) values compared to their room-temperature counterparts, indicating a notable loss in spatial resolution under cryogenic imaging conditions (**Fig. 2**). Visualization of PSF profiles from multiple phase-retrieved models based on different beads acquired at cryogenic temperatures (**Supplementary Fig. SS1**) consistently showed similarly enlarged FWHM values. This observation underscores the detrimental effect of cryogenic conditions on both spatial resolution and the achievable localization precision.

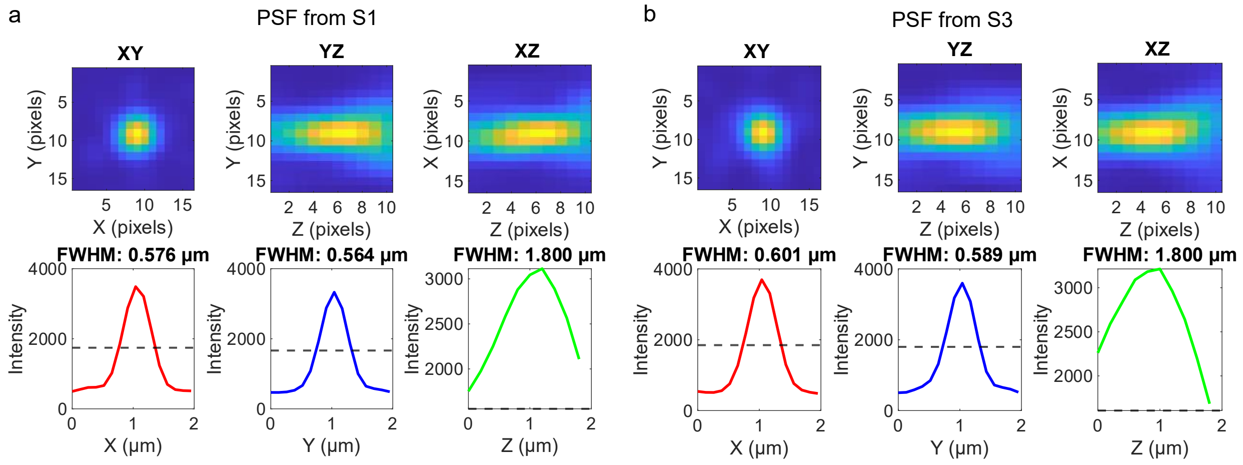

**Supplementary Fig. SS 1. Two PSF profiles under cryo-FLM exhibit consistent larger FWHMs.**
**(a)** Experimental PSF profiles in the XY, XZ, and YZ views for a bead located in FOV1-S1 (see **Fig. 3**).
**(b)** Experimental PSF profiles in the XY, XZ, and YZ views for a bead located in FOV1-S3 (see **Fig. 3**).

**FOV-dependent and system-specific aberrations exhibit strong spatial heterogeneity.** Using our imaging strategy that decouples system-specific effects from sample-induced aberrations (**Fig. 3a**, ***Methods***), we imaged the same fluorescent beads across overlapping FOVs, generating paired datasets such as {S1, S1′}, {S2, S2′}, {S3, S3′}, and {S4, S4′}. In this setup, each bead is captured at different FOV-dependent positions. By comparing images of the same bead across nearby FOVs, we effectively eliminate sample-induced aberrations, allowing any differences in the retrieved PSF models to be attributed solely to system-specific aberrations across the FOV. In contrast, comparisons among beads located in different positions, such as S1–S4, capture all sources of FOV-dependent aberrations, including system-specific distortions, refractive index mismatches, and local sample heterogeneities. Additional bead pairs, such as {S1, S3′}, {S2, S1′}, {S3, S4′}, and {S4, S2′}, are also informative, as they originate from the same square but lie in adjacent FOVs. These pairs are expected to reflect similar system-induced aberrations and focus on the difference brought by the sample-induced aberrations.

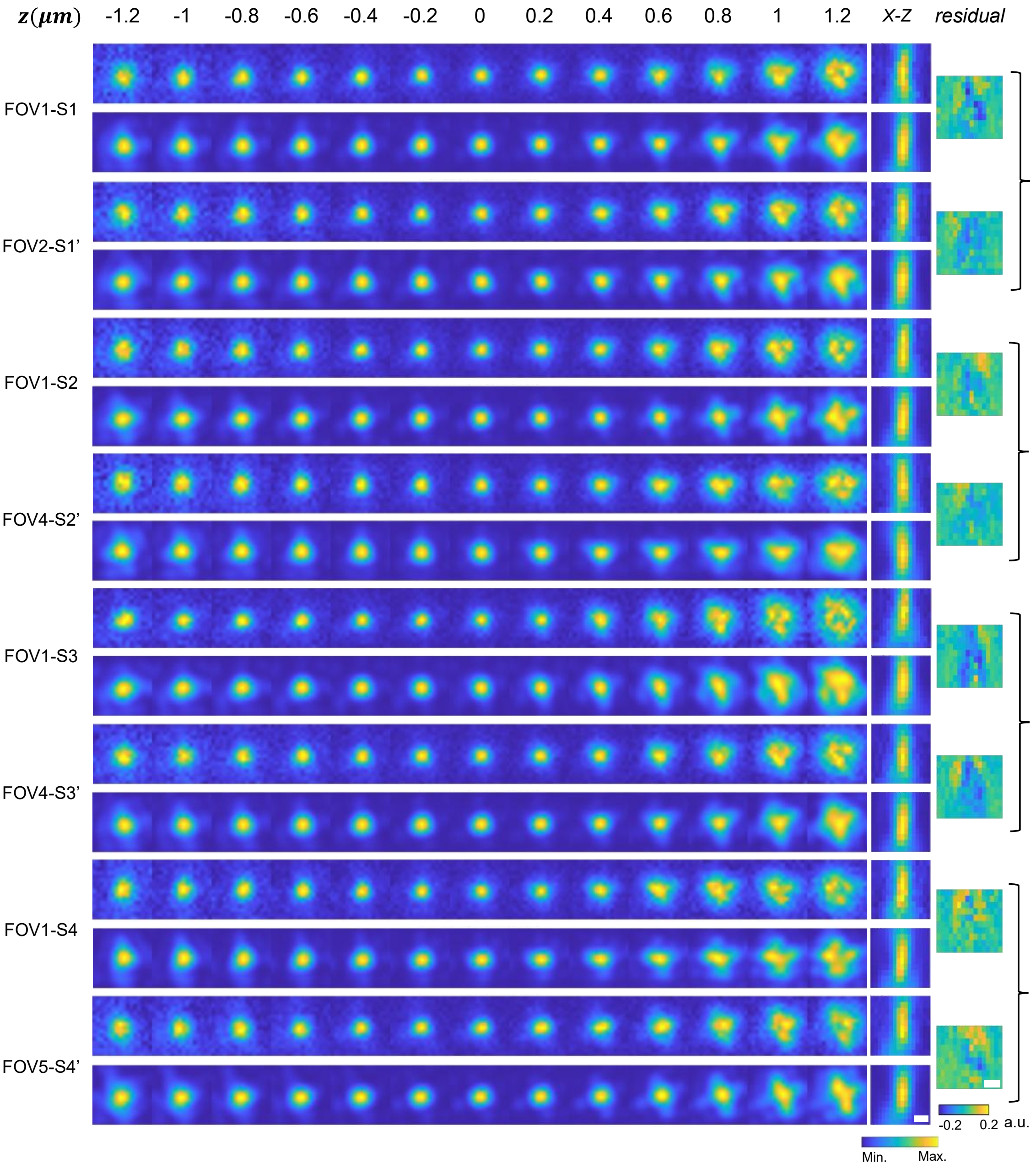

**Supplementary Fig. SS 2. Experimental and phase-retrieved PSFs in FOV1–FOV5.**

xy projections of the experimental and phase-retrieved PSFs across a z-range from –1.2 μm to +1.2 μm for beads in FOV1–FOV5 in **Fig. 3**. Corresponding xz views and residuals between the experimental and reconstructed PSFs are also presented. Brackets indicate the same bead imaged at different positions in nearby FOVs, highlighting the variability in system-induced aberrations. Scale bar: 500 nm.

Despite significant noise, which poses challenges to the phase retrieval process, the retrieved PSFs closely resemble the experimentally measured ones (**Supplementary Fig. SS2**). Differences observed between the same bead imaged in nearby FOVs primarily reflect system-specific aberrations, though minor contributions from additional factors, such as metal-induced scattering from the bars of the EM grid, cannot be entirely ruled out. Notably strong system-specific aberrations were observed in the left region of the FOV, as indicated by elongated XY PSF profiles at axial positions between 1.0 μm and 1.2 μm for bead pairs S3/S3′ and S4/S4′. Moreover, distinct PSF shapes were also observed between different beads within the same FOV, underscoring the complex, position-dependent nature of aberrations arising from a combination of system-specific factors, refractive index mismatches, and heterogeneous sample composition.

To further assess these aberrations, we analyzed the pupil-phase maps corresponding to primary aberrations retrieved from beads positioned at S1–S4 and their paired locations S1′–S4′ (**Supplementary Fig. SS3**). First, the pupil-phase maps for beads S1–S4 reveal the primary aberrations present across the entire FOV. Second, the differences between paired beads (S1/S1′, S2/S2′, S3/S3′, and S4/S4′) highlight the system-specific aberrations, with the polynomial coefficients directly quantifying the magnitude of each primary aberration. Several important observations emerge from this analysis. Aberrations vary significantly across the FOV and are not only position-dependent but also unpredictable in both type and magnitude. This variability reflects contributions from both system-specific and sample-induced factors. Interestingly, spherical aberration was not consistently dominant, despite expectations based on refractive index mismatch. Instead, trefoil and astigmatism frequently emerged as the dominant modes, with coma appearing occasionally. Moreover, the magnitude of these aberrations varied considerably, emphasizing the spatial complexity of the aberration landscape under cryogenic imaging conditions.

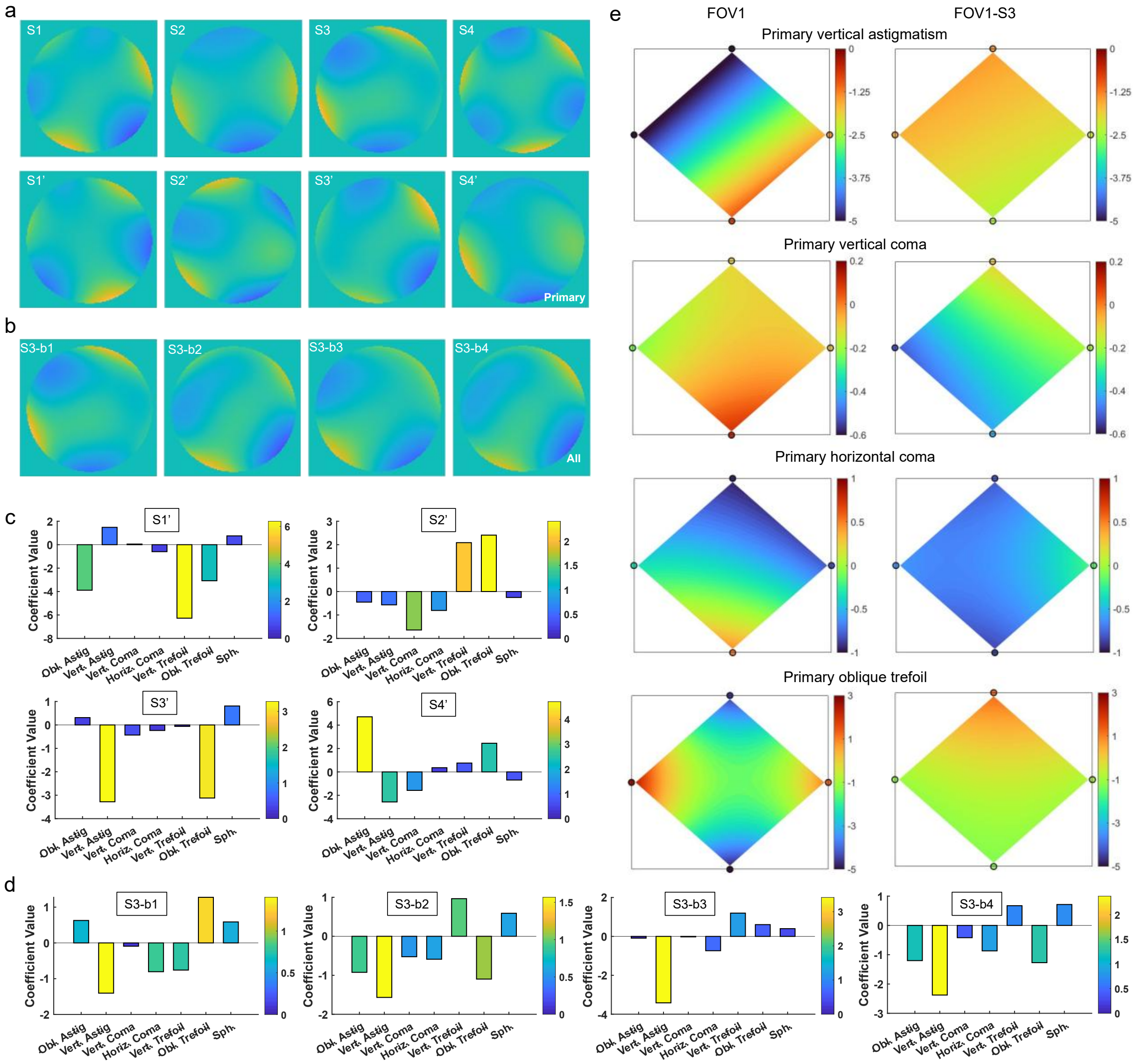

**Supplementary Fig. SS 3. FOV-dependent and square-dependent aberrations exhibit different levels of variability.**

**(a)** Phase maps of pupil functions representing primary FOV-dependent aberrations for PSF models retrieved from beads located in FOV1 (S1–S4) and FOV2–FOV5 (S1′-S4′). Differences between paired images of the same bead (e.g., {S1, S1′}) highlight system-specific aberration variability. **(b)** Phase maps of pupil functions showing all aberrations for PSF models retrieved from four beads located within the same square (S3) in FOV1. **(c)** Zernike coefficients corresponding to primary aberrations for PSF models retrieved from beads in S1′–S4′. **(d)** Zernike coefficients for primary aberrations from four beads in S3, illustrating square-level aberration variation. **(e)** Contour plots showing the spatial variation in the amplitudes of four primary aberration modes, comparing FOV-dependent aberrations (FOV1) and square-dependent aberrations (FOV1-S3).

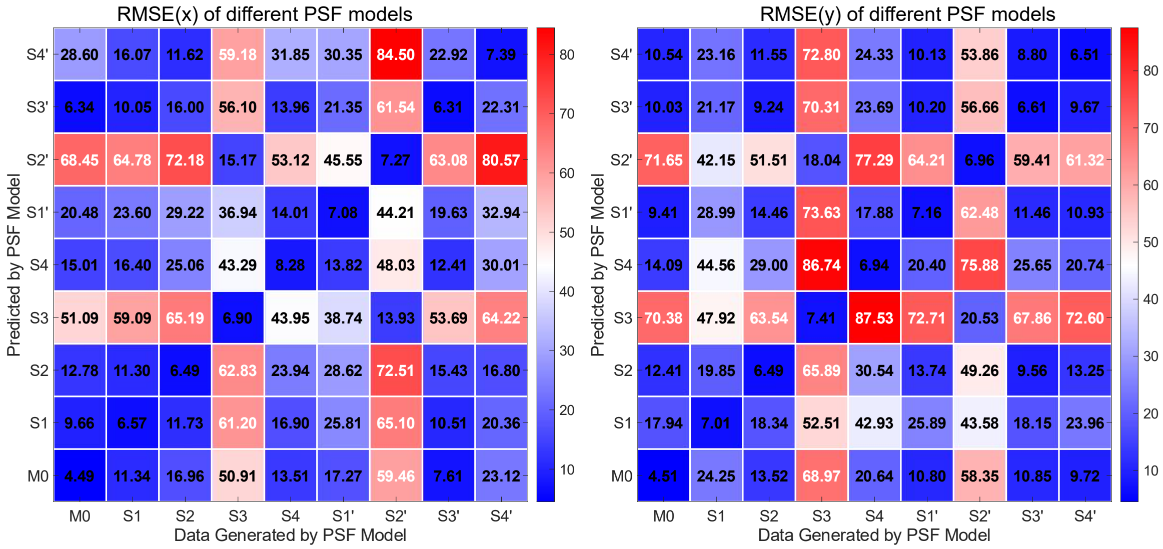

**Supplementary Fig. SS 4. FOV-dependent aberrations result in significant lateral localization RMSEs, reaching up to ~90 nm.**

Simulated datasets generated using the theoretical PSF model and PSF models retrieved from FOV1–FOV5 (**Fig. 3**), spanning axial positions from −0.7 µm to +0.7 µm in 100 nm steps, with 100 frames per z-position. Simulations were performed with a signal photon count of $1 \times 10^4$ and a background photon count of 400. These models were used for 3D localization, and the RMSE in x- and y-localization was calculated.

**FOV-dependent aberrations result in significant lateral localization RMSEs.** Our localization analysis highlights the significant impact of FOV-dependent aberrations, particularly when mismatched PSF models are used (**Supplementary Fig. SS4; Supplementary Fig. 5-6**). The resulting lateral localization RMSE can be substantial, comparable to those induced by using a theoretical pupil function, with values reaching up to ~90 nm. Such errors are considerable and could compromise downstream applications that demand high spatial precision, such as high-magnification cryo-electron microscopy and tomography (cryo-ET) and cryo-focused ion beam milling scanning electron microscopy (cryo-FIB-SEM). Among all tested PSF models, two which are retrieved from beads in FOV1-S3 and FOV3-S2′ stand out for producing notably higher localization errors compared to others. In contrast, most PSF models retrieved from other grid squares, as well as the theoretical pupil model, yield lateral localization RMSEs ranging from ~5 to 30 nm. A closer inspection of **Supplementary Fig. SS3** reveals the origins of these errors. The PSF retrieved from position S3 shows pronounced elongation in the XY-plane, especially at defocused axial positions, while the PSF from S2′ appears compressed in the same plane. These distinct shape distortions likely play a major role in the elevated localization errors observed for these positions. Although mismatched PSF models from other TEM grid squares can also result in large axial localization RMSEs up to ~210 nm, these values are still significantly lower than those obtained using the theoretical PSF model, which produces axial errors in the range of ~250–400 nm.

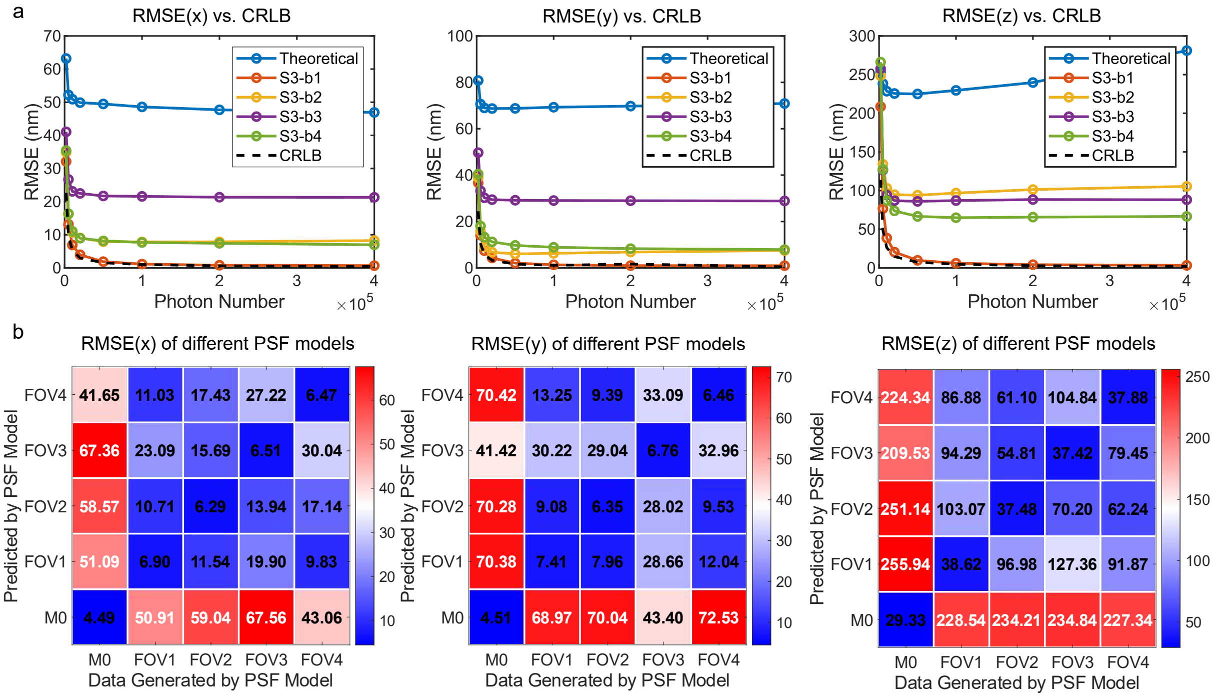

**Supplementary Fig. SS 5. Square-dependent PSF models (FOV1-S3) partially mitigate localization errors compared to FOV-dependent aberrations.**

Test data were generated using the PSF model retrieved from a bead S3-b1 (FOV1-S3-Bead1 in **Fig. 3a**), spanning axial positions from –0.7 μm to +0.7 μm in 100 nm steps (100 frames per z-position). Simulations were performed with a signal photon count of $1 \times 10^4$ and a background photon count of 400. Five PSF models were used for 3D localization: Model 0 (theoretical pupil function), Models 1–4 (experimental PSFs from beads in FOV1-S3). **(a)** RMSE in x-, y-, and z-localization as a function of photon number, compared to the corresponding CRLBs. **(b)** All these five models were used for 3D localization, and the RMSE in x- and y-localization was calculated.

**Square-dependent aberrations exhibit greater spatial consistency and partially mitigate localization errors.** Two key findings emerge from our localization analysis regarding square-dependent aberrations (**Supplementary Figs. SS5**). First, different PSF models exhibit varying performance in both lateral and axial localization accuracy, emphasizing the influence of aberration differences on localization outcomes. Second, PSF models derived from beads located within the same TEM grid square (e.g., FOV1-S3 in **Fig. 3**) display more consistent characteristics and yield significantly lower localization RMSEs, both laterally and axially, compared to the theoretical pupil-based PSF model. These findings highlight the advantage of using region-specific, phase-retrieved PSF models for accurate localization, particularly under aberration-rich conditions, although these models still exhibit some degree of variability.

##### 1.3. More results related to grid-dependent aberrations

**Aberrations in FOV3 across the whole TEM grid are variable.** FOV3 (**Fig. 5**) was selected for simulation because it corresponds to the same bead (FOV1-S3, FOV1-S3-Bead1) analyzed in **Fig. 3**. As we selected beads at similar relative positions within each FOV, this configuration primarily reflects differences arising from sample-induced aberrations. Although PSF models from other regions of the grid exhibit slightly higher NCC with the PSF from FOV3-S3, the resulting localization errors, particularly in the axial (z) dimension, remain high (**Supplementary SS 6**).

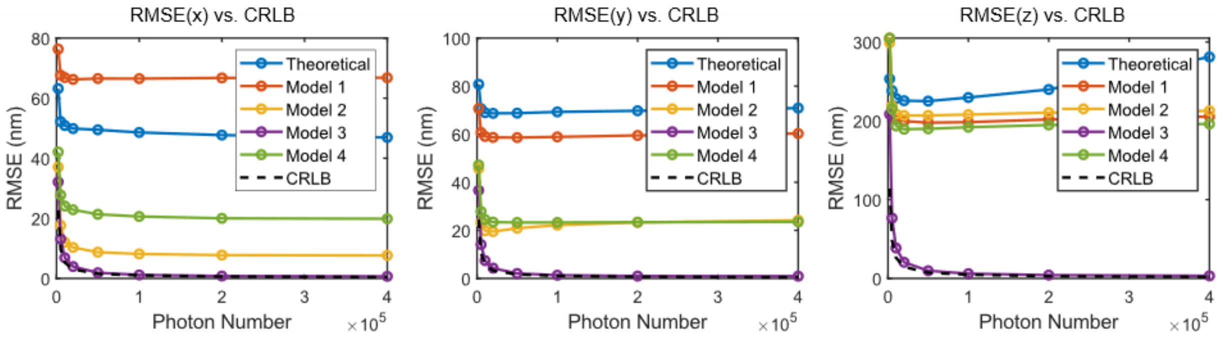

**Supplementary Fig. SS 6. Grid-dependent PSF models introduce high and variable localization errors.**

Test data were generated using the PSF model retrieved from a bead in FOV3 (**Fig. 5a**), spanning axial positions from –0.7 μm to +0.7 μm in 100 nm steps (100 frames per z-position). Simulations were performed with a signal photon count of $1 \times 10^4$ and a background photon count of 400. Five PSF models were used for 3D localization: Model 0 (theoretical pupil function), Models 1–4 (experimental PSFs from beads in FOV1-4 in **Fig. 5a**). RMSE in x-, y-, and z-localization as a function of photon number, compared to the corresponding CRLBs.

**Comparison of FOV-, square-, and grid-dependent aberrations.** Because the same beads were used across our analyses of FOV-dependent, square-dependent, and grid-dependent aberrations, we were able to perform direct comparisons (**Extended Data Figs. 3–4**). While square-dependent aberrations show relatively consistent best behavior, it is difficult to determine whether FOV-dependent or grid-dependent aberrations are better in localization. However, key insights emerge from the localization analysis. Notably, the z-localization RMSE remains high for both FOV-dependent and grid-dependent aberrations. In contrast, the lateral (x and y) localization errors for grid-dependent aberrations are comparatively low—approaching those observed in square-dependent cases. This trend aligns with the high similarity seen in the XY projections of the PSFs (**Supplementary Figs. SS 7**), suggesting that lateral localization is preserved when XY shape similarity is maintained. Since grid-dependent aberrations primarily reflect sample-induced effects, they tend not to alter the XY shape of the PSF drastically, which explains the preserved lateral localization accuracy. However, axial localization proves far more sensitive to subtle aberrations, particularly those introduced by sample-specific effects. Importantly, increasing the photon count alone does not compensate for axial localization errors, reinforcing the necessity of accurate, location-specific PSF models to achieve precise 3D localization in cryogenic imaging systems (**Supplementary Fig. SS 8**).

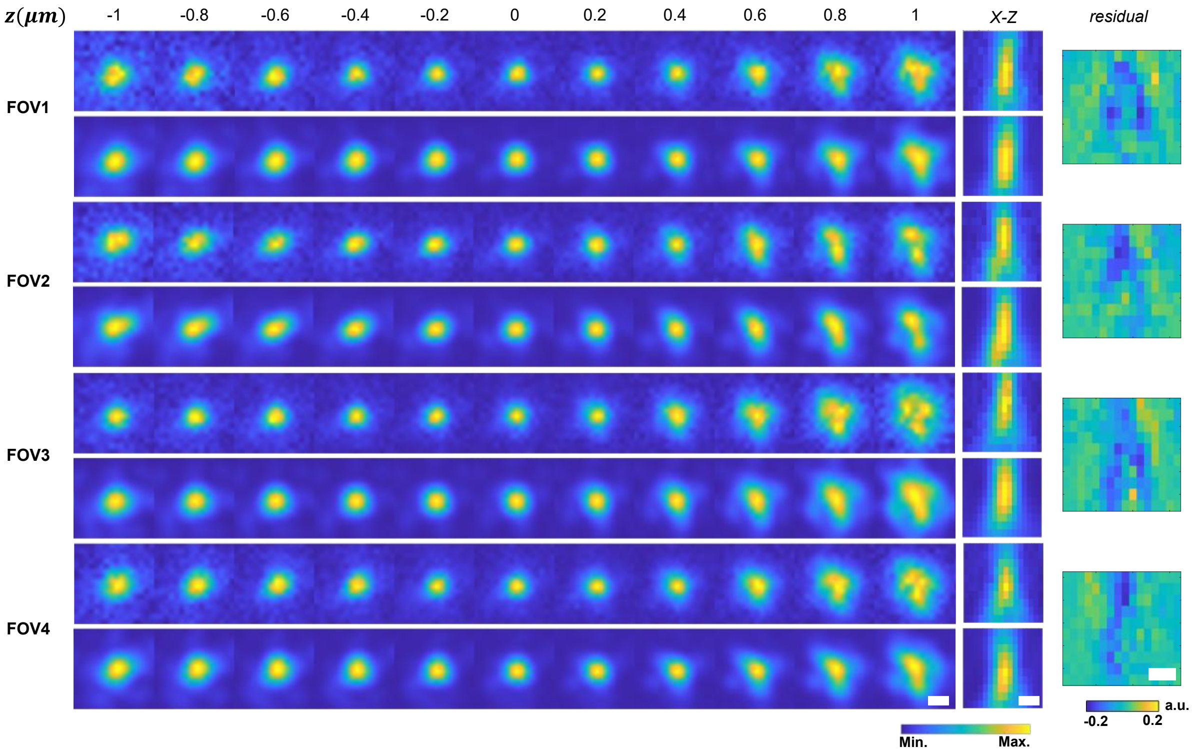

**Supplementary Fig. SS 7. Experimental and phase-retrieved PSFs in FOV1–FOV4 of the TEM grid.**

The xy projections of the experimental and phase-retrieved PSFs across a z-range from –1 μm to +1 μm for beads in FOV1–FOV4 in **Fig. 5**. Corresponding xz views and residuals between the experimental and reconstructed PSFs are also presented. Scale bar: 500 nm.

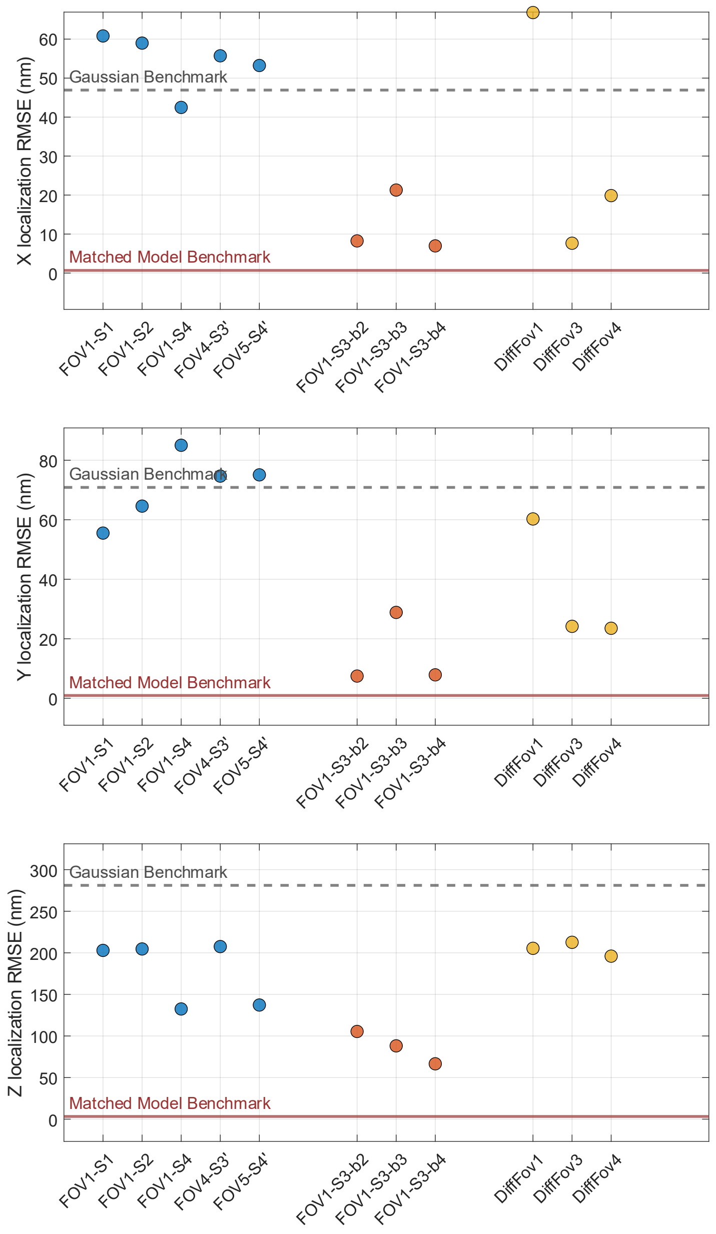

**Supplementary Fig. SS 8. Localization RMSE as a function of FOV-dependent, square-dependent, and grid-dependent aberration mismatches.**

Test datasets were generated using the PSF model from FOV3 in **Fig. 5a** (identical to the PSF models used in FOV1-S3 and FOV-S3-b1 in **Fig. 3a**), spanning axial positions from −0.7 μm to +0.7 μm in 100 nm steps, with 100 frames per z-position. Simulations were conducted with a signal photon count of $4 \times 10^5$ and a background photon count of 400 photons per pixel. Localization was performed using mismatched PSF models corresponding to one FOV, one square, and one grid. RMSE in x-, y-, and z-localization is presented. Scale bar: 500 nm.

**Another case of grid-dependent aberrations further validates their variability.** Due to the presence of strong and complex sample-induced aberrations in the cryo-sample, we selected an alternative square (S1) for further grid-dependent aberrations analysis (**Supplementary Figs. SS9**). FOV3 was chosen for simulation and corresponds to the same bead (FOV1-S1) analyzed in **Fig. 3**. The aberration analysis and localization analysis (**Supplementary Figs. SS10**; **Supplementary Fig. 13-15**) were conducted using the same approach as for square S3.

Two key conclusions emerge from this analysis:

(1) Grid-dependent PSF models exhibit variable performance in both lateral and axial localization. Specifically, the PSF model retrieved from FOV1 yields the highest error in x-localization, while the model from FOV3 performs worst in y. However, all models produce comparable z-localization errors. Notably, this behavior differs from the results observed for square S3, where axial errors were more variable.

(2) The localization errors resulting from aberrations across the entire TEM grid differ in magnitude from those caused by FOV-dependent aberrations, and it is difficult to determine which condition yields more consistent or accurate localization as we demonstrated for the FOV1-S3 in **Fig. 3**. All these findings indicate that even subtle spatial variations in aberrations across the grid can unpredictably impact localization performance, reinforcing the importance of implementing spatially adaptive PSF correction strategies in high-precision cryo-FLM applications.

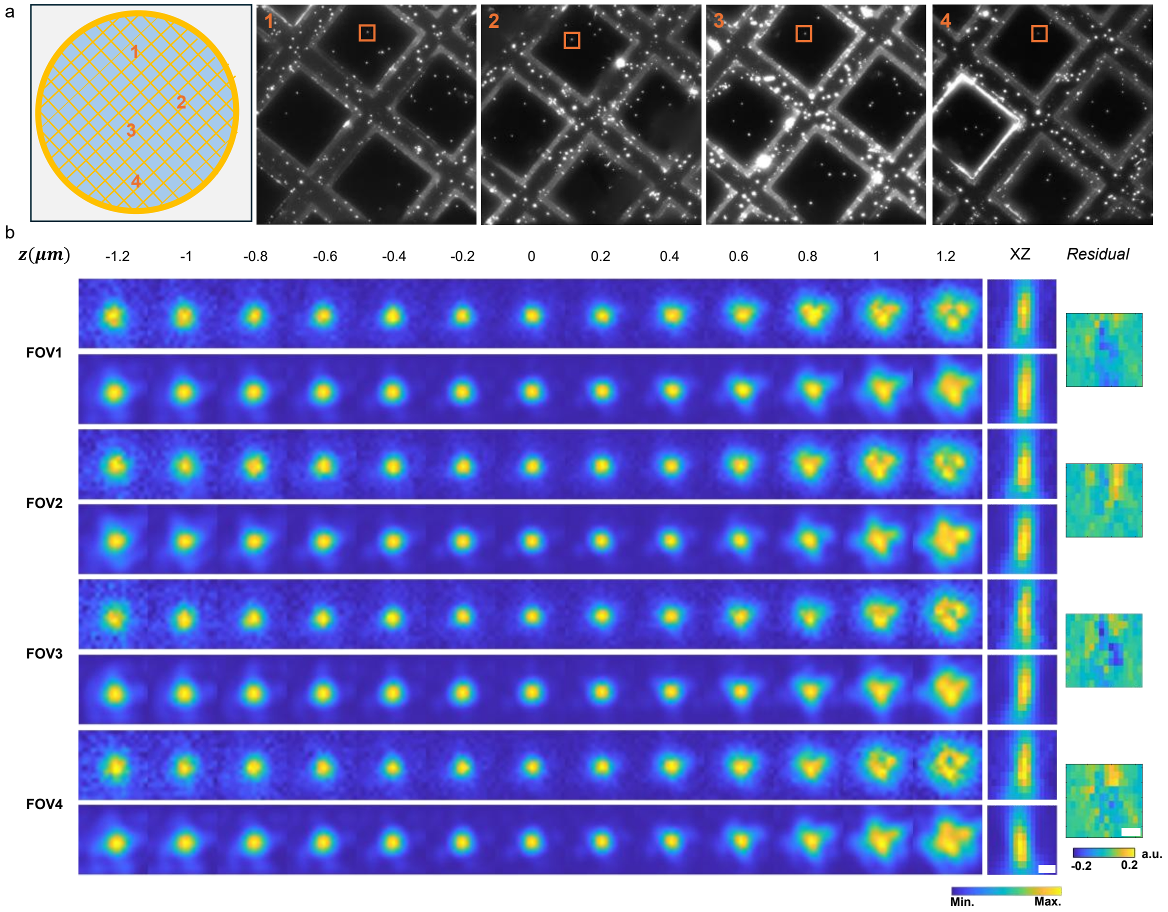

**Supplementary Fig. SS 9. Another case study of grid-dependent aberrations.**

**(a)** Layout of different FOVs across the EM grid, with beads selected from the same relative position (S1) within each FOV for aberration analysis. **(b)** Experimental and phase-retrieved PSFs in FOV1–FOV4. Shown are the XY projections of the experimental and phase-retrieved PSFs over a z-range from −1.2 µm to +1.2 µm for the selected beads. Corresponding XZ views and residuals (experimental minus reconstructed PSFs) are also shown. Scale bar: 500 nm.

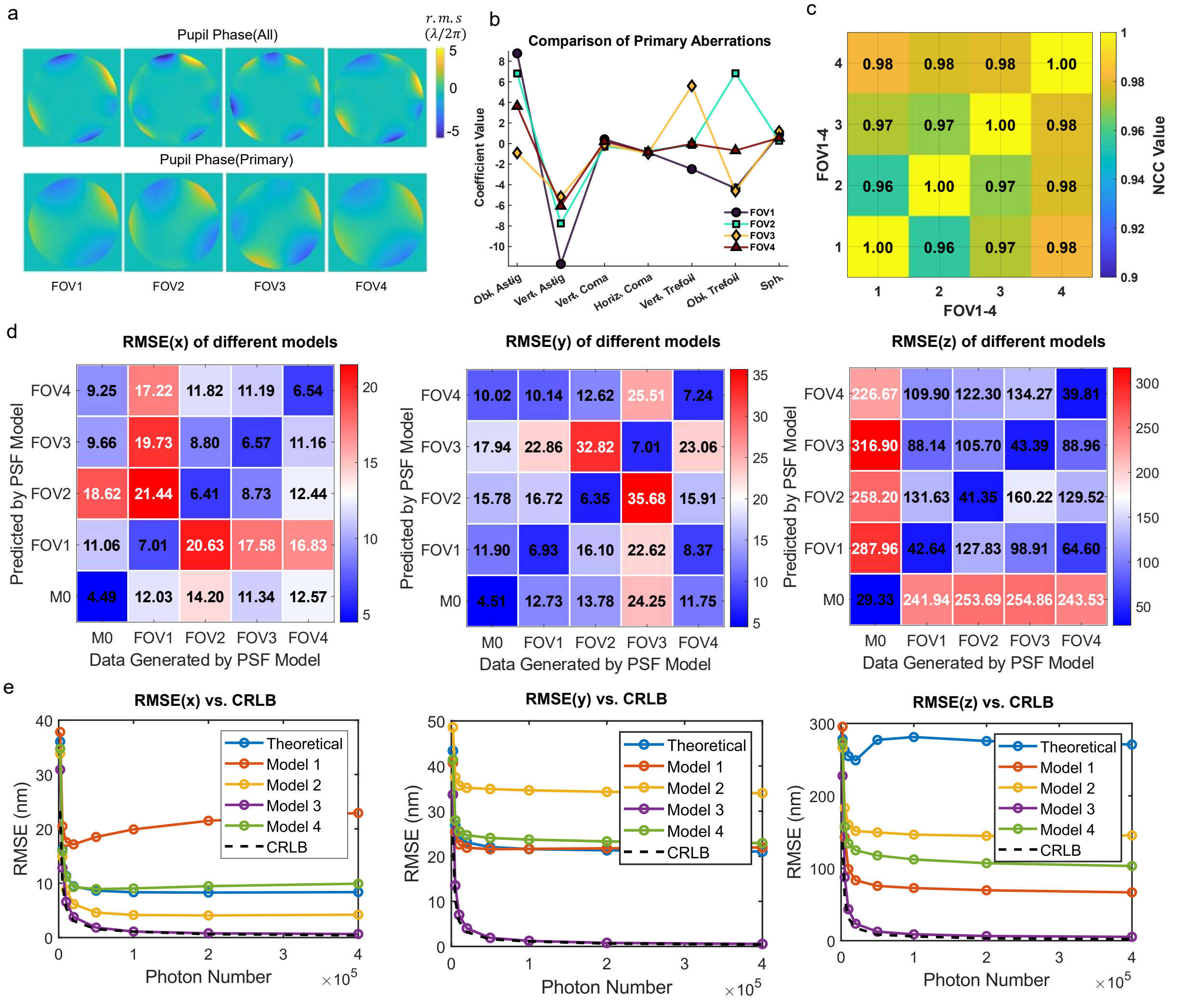

**Supplementary Fig. SS 10. Grid-dependent aberrations and corresponding localization analysis.** **(a)** Phase maps of the pupil functions showing both total aberrations and primary aberrations for PSF models retrieved from beads at position S1 in FOV1–FOV4 (**Supplementary Fig. SS10**). **(b)** Zernike polynomial coefficients corresponding to primary aberrations from the same set of beads. **(c)** NCC values among the four retrieved PSF image stacks, indicating the degree of similarity between PSFs across FOVs. **(d)** Test data were generated using the PSF model retrieved from a bead in FOV3 (**Supplementary Fig. SS10**), spanning axial positions from –0.7 μm to +0.7 μm in 100 nm steps (100 frames per z-position). Simulations were performed with a signal photon count of $1 \times 10^4$ and a background photon count of 400. Five PSF models were used for 3D localization: Model 0 (theoretical pupil function), Models 1–4 (experimental PSFs from beads in FOV1-4). Summary of localization RMSE when using these PSFs for both data simulation and localization, **(e)** RMSE in x-, y-, and z-localization as a function of photon number, compared to the corresponding CRLBs.

##### 1.4. Alignment of the focus and lateral offsets.

**Alignment localization error induced by PSF misalignment.** Although defocus and lateral (xy) offsets are corrected during the preprocess, residual misalignments between two PSF models may still persist. These misalignments can introduce systematic localization errors when simulated datasets are generated using one PSF model and fitted using another. To mitigate this bias, we subtract the average localization error across all axial positions from the localization error at each specific z position, as described by the following equation:

$$Bias_{\mathrm{avg}} = \frac{1}{\mathrm{M}} \sum_{\mathrm{k}=1}^{\mathrm{M}} \sum_{i=1}^{N} \hat{\mathrm{z}}_{\mathrm{i}}(\mathrm{k}),$$

$$\mathrm{B}(\mathrm{k}) = \frac{1}{\mathrm{N}} \sum_{\mathrm{i}=1}^{\mathrm{N}} \left| \hat{\mathrm{z}}_{\mathrm{i}}(\mathrm{k}) - Bias_{\mathrm{avg}} - \mathrm{z}_{\mathrm{i}}^{\mathrm{GT}}(\mathrm{k}) \right|$$

where $\mathrm{k} = 1,2, \dots ,15$ indexes the axial positions and $\mathrm{N} = 100$ is the number of frames per position.

However, this correction may significantly underestimate the true localization errors, as it relies on the assumption that aberration-induced errors are zero-mean around the focal plane. This assumption breaks down in the presence of asymmetric aberrations such as astigmatism and coma, which distort the PSF and break its symmetry. In our experiments, we consistently observed strong astigmatism and coma, along with clearly asymmetric PSF shapes across all datasets. When both the simulated data and localization analysis use asymmetric PSF models, the assumption of zero-mean aberration-induced errors no longer holds, leading to a systematic underestimation of the true localization error. Nevertheless, let us examine what the data reveal.

We computed aligned localization errors for all experiments in this study, including the cryoT datasets (**Supplementary Fig. SS11**), the FOV-dependent aberrations (**Supplementary Fig. SS12-13**), square-dependent aberrations (**Supplementary Figs. SS14-15**), and two case studies of grid-dependent aberrations (**Supplementary Figs. SS16-19**). Across all cases, alignment consistently reduced both lateral and axial localization errors. However, the key findings and conclusions remain unchanged:

1. **Model mismatch introduces substantial localization errors.** Matched PSF models yield the best precision, achieving sub-1 nm lateral and ~5 nm axial RMSE. In contrast, FOV-dependent and grid-dependent aberrations introduce substantial errors. Grid-dependent aberrations show slightly better lateral performance (up to ~15 nm) compared to FOV-dependent ones (up to ~30 nm), while both result in comparable axial errors (up to ~150 nm). Square-dependent aberrations mitigate localization errors more effectively, reducing them to ~10 nm laterally and ~40 nm axially. Although these values are smaller than those reported in our previous results, as expected given the alignment, the core conclusion holds.

2. **The magnitude of improvement varies with PSF mismatch severity.** Even when the simulated dataset was generated using the same PSF model, we observed different levels of error reduction depending on the mismatched model used for localization. Improvements were most pronounced when the mismatch was large, as in the case of PSF models retrieved from S3 and S2'. This finding suggests that a significant portion of the localization error arises from PSF model mismatch, rather than pure misalignment.

These results demonstrate that even under the overly optimistic assumption that all improvements stem from alignment correction, substantial residual errors persist. This underscores the critical importance of accurate, position-specific aberration characterization across the field of view to enable precise localization.

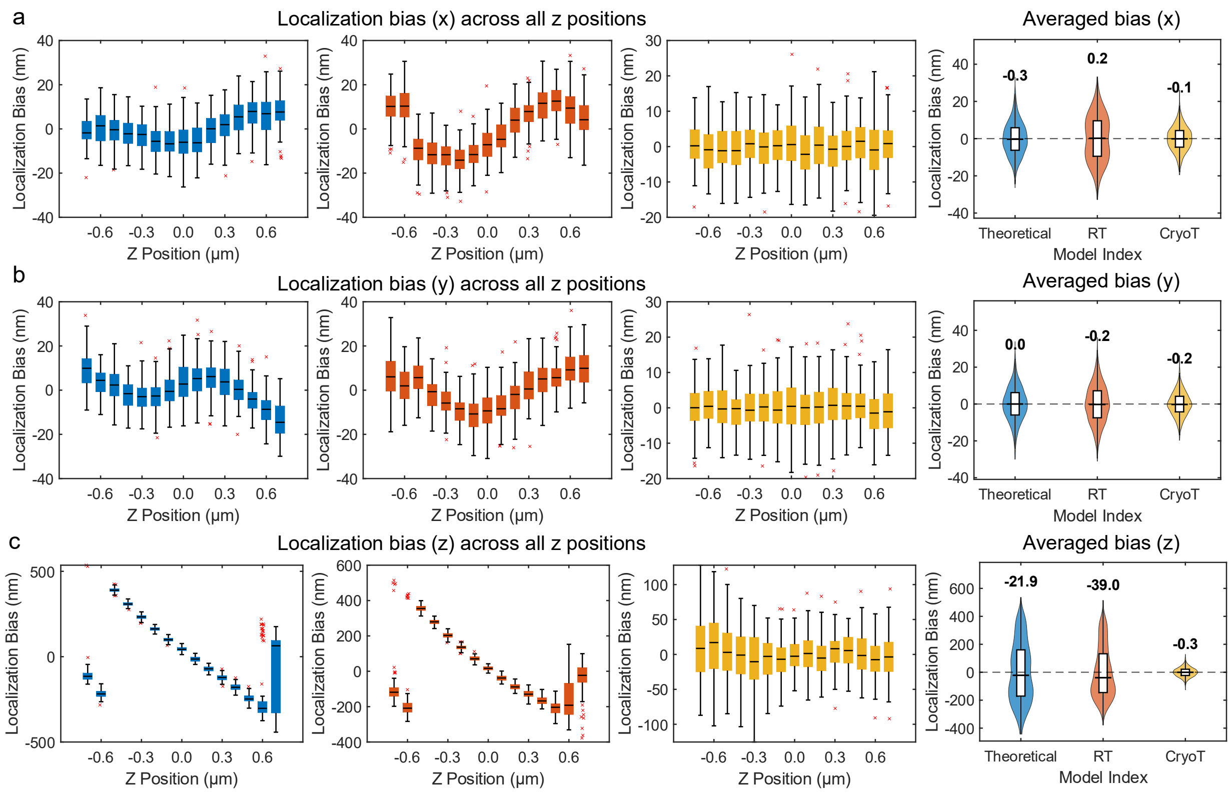

**Supplementary Fig. SS 11. Aligned localization bias analysis of cryo-FLM.**

Three PSF models were used for 3D localization: a theoretical pupil-based model, the room-temperature (RT) PSF model, and the cryogenic-temperature (CryoT) PSF model (**Fig. 2**). Simulated datasets were generated using the CryoT PSF model across axial positions from −0.7 µm to +0.7 µm in 100 nm increments, with 100 frames per z-position. Simulations were performed with a signal photon count of $1 \times 10^4$ and a background photon count of 400. **(a–c)** Localization bias in x, y, and z directions, respectively, across the axial range and corresponding bias distributions aggregated over all positions are also shown. Localization biases were aligned by subtracting the average bias across all localizations, highlighting relative deviations.

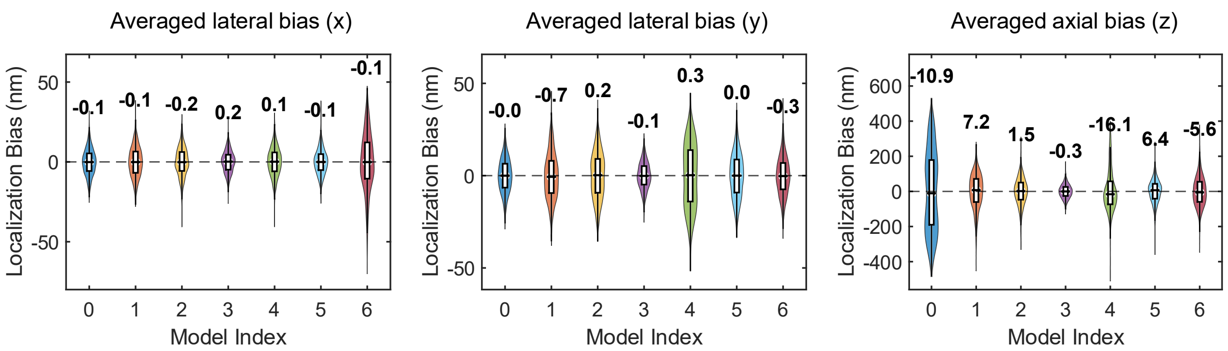

**Supplementary Fig. SS 12. Aligned localization bias analysis induced by FOV-dependent aberrations.**

Test data were generated using the PSF model retrieved from a bead in FOV1-S3 (**Fig. 3a**), spanning axial positions from –0.7 μm to +0.7 μm in 100 nm steps (100 frames per z-position). Simulations were performed with a signal photon count of $1 \times 10^4$ and a background photon count of 400. Seven PSF models were used for 3D localization: Model 0 (theoretical pupil function), Models 1–4 (experimental PSFs from beads in FOV1-S1 to S4), and Models 5–6 (PSFs from S3′ and S4′, representing system-induced and sample-induced aberrations, respectively). Lateral and axial localization bias as a function of axial position, and corresponding bias distributions aggregated over all positions are shown. Localization biases were aligned by subtracting the average bias across all localizations, highlighting relative deviations.

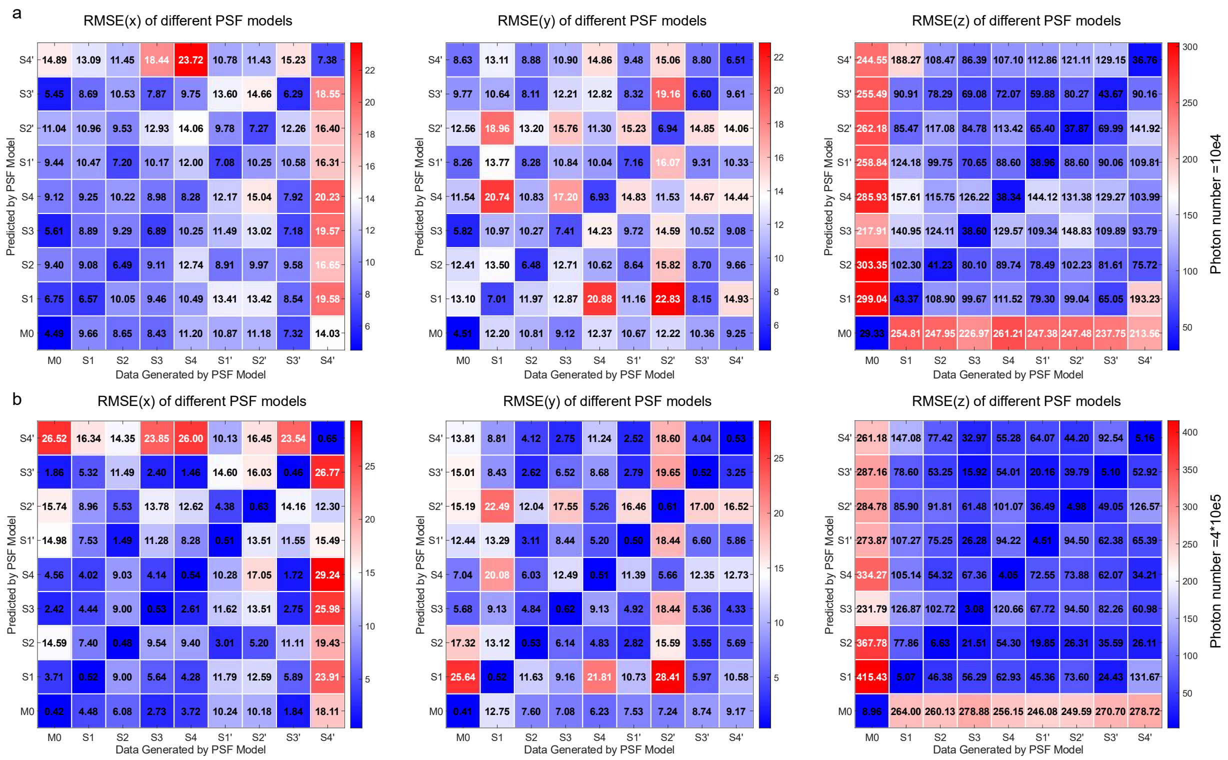

**Supplementary Fig. SS 13. Aligned localization RMSE induced by FOV-dependent aberrations.** Simulated datasets were generated using the theoretical PSF model and PSF models retrieved from FOV1–FOV5 (**Fig. 3**), spanning axial positions from −0.7 µm to +0.7 µm in 100 nm steps, with 100 frames per z-position. Simulations were performed with a signal photon count of 1 × $10^4$ and 4 × $10^5$ for **(a)** and **(b)**, and a background photon count of 400 photons per pixel. These PSF models were used for 3D localization, and RMSE in x-, y-, and z-localization was calculated after alignment to remove average bias.

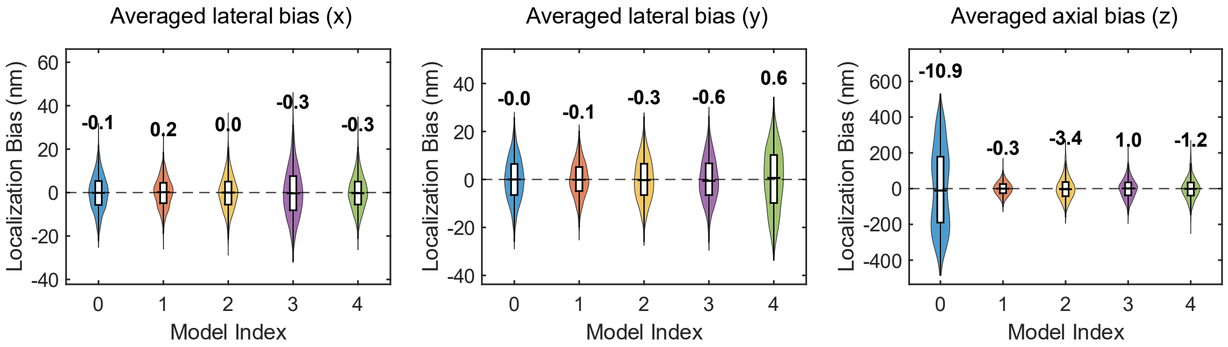

**Supplementary Fig. SS 14. Aligned localization bias induced by square-dependent aberrations (FOV1-S3).**

Test data were generated using the PSF model retrieved from a bead in FOV1-S3-Bead1 (**Fig. 3a**), spanning axial positions from –0.7 µm to +0.7 µm in 100 nm steps (100 frames per z-position). Simulations were performed with a signal photon count of 1 × $10^4$ and a background photon count of 400. Five PSF models were used for 3D localization: Model 0 (theoretical pupil function), Models 1–4 (experimental PSFs from beads in FOV1-S3). Localization bias distributions aggregated over all positions are also shown. Localization biases were aligned by subtracting the average bias across all localizations, highlighting relative deviations.

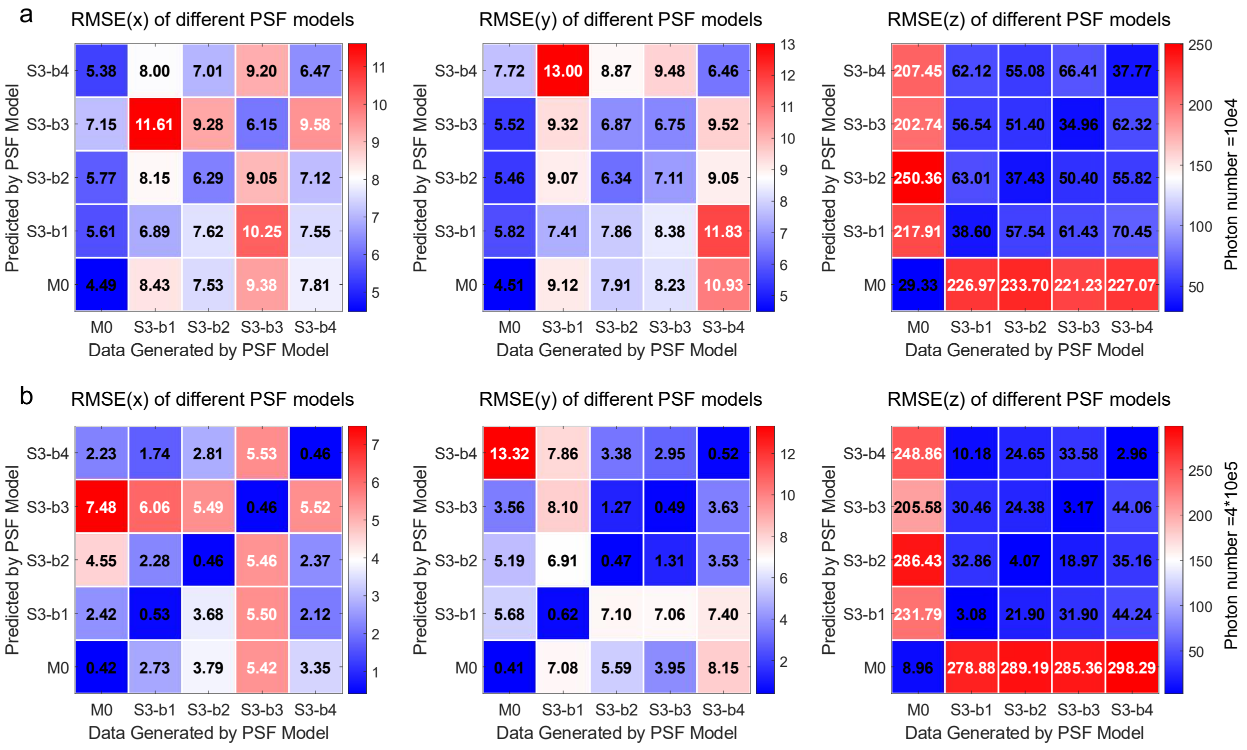

**Supplementary Fig. SS 15. Aligned localization RMSE induced by square-dependent aberrations.** Test data were generated using the PSF model retrieved from a bead in FOV1-S3-Bead1 (**Fig. 3a**), spanning axial positions from –0.7 μm to +0.7 μm in 100 nm steps (100 frames per z-position). Simulations were performed with a signal photon count of $1 \times 10^4$ and $4 \times 10^5$ for **(a)** and **(b)**, and a background photon count of 400 photons per pixel. Five PSF models were used for 3D localization: Model 0 (theoretical pupil function), Models 1–4 (experimental PSFs from beads in FOV1-S3). These PSF models were used for 3D localization, and RMSE in x-, y-, and z-localization was calculated after alignment to remove average bias.

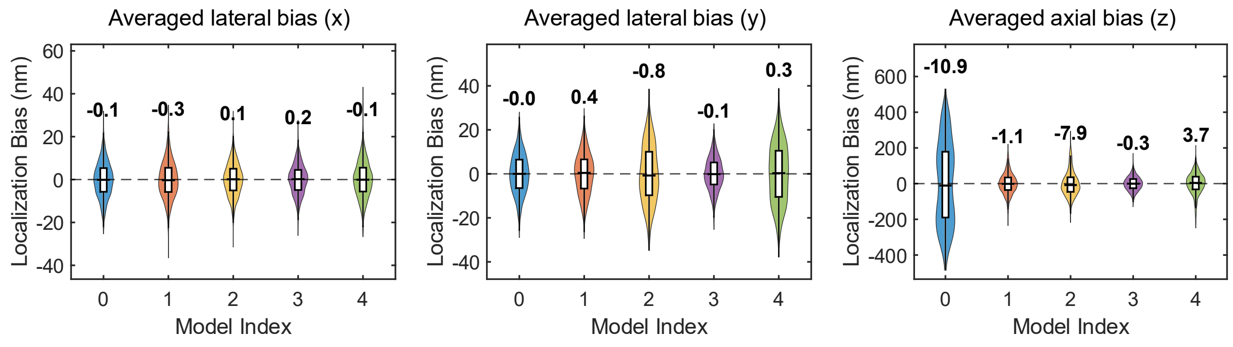

**Supplementary Fig. SS 16. Aligned localization bias induced by grid-dependent aberrations.**

Test data were generated using the PSF model retrieved from a bead in FOV3 (**Fig. 5a**), spanning axial positions from –0.7 μm to +0.7 μm in 100 nm steps (100 frames per z-position). Simulations were performed with a signal photon count of $1 \times 10^4$ and a background photon count of 400. Five PSF models were used for 3D localization: Model 0 (theoretical pupil function), Models 1–4 (experimental PSFs from beads in FOV1-4 in **Fig. 5a**). Localization bias distributions aggregated over all positions are also shown. Localization biases were aligned by subtracting the average bias across all localizations, highlighting relative deviations.

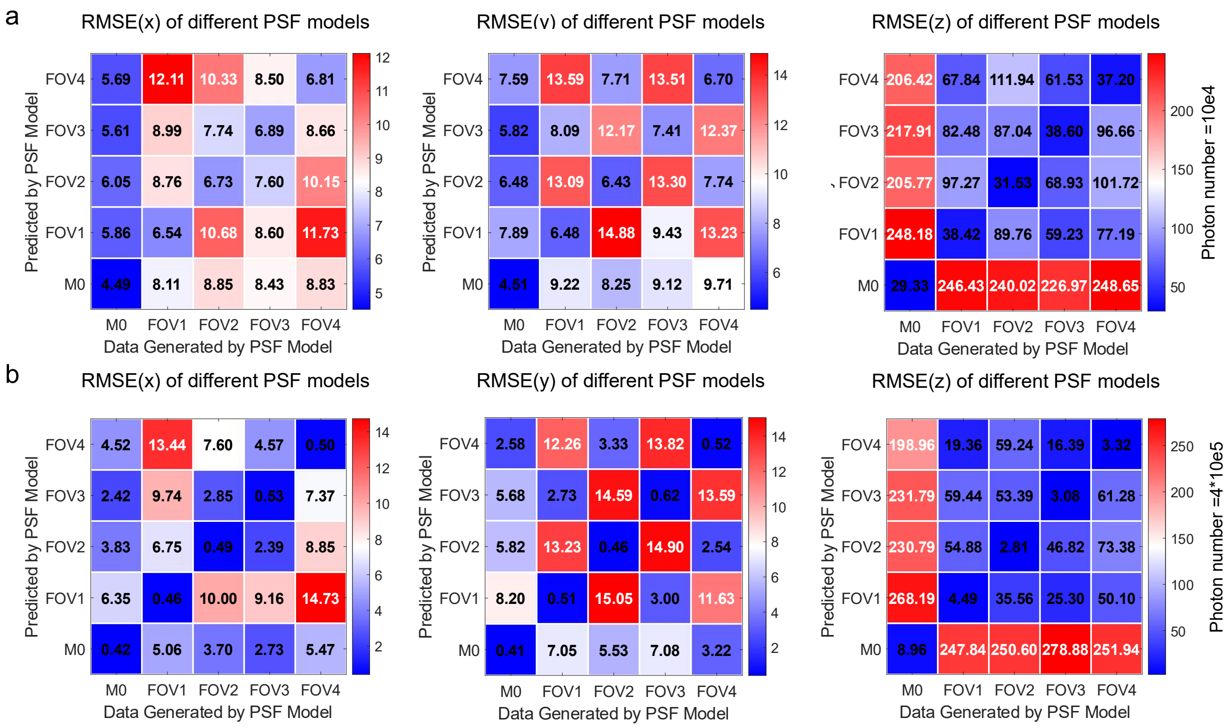

**Supplementary Fig. SS 17. Aligned localization RMSE induced by grid-dependent aberrations.** Test data were generated using the PSF model retrieved from a bead in FOV3 (**Fig. 5a**), spanning axial positions from –0.7 μm to +0.7 μm in 100 nm steps (100 frames per z-position). Simulations were performed with a signal photon count of $1 \times 10^4$ and $4 \times 10^5$ for **(a)** and **(b)**, and a background photon count of 400 photons per pixel. Five PSF models were used for 3D localization: Model 0 (theoretical pupil function), Models 1–4 (experimental PSFs from beads in FOV1-4 in **Fig. 5a**). These PSF models were used for 3D localization, and RMSE in x-, y-, and z-localization was calculated after alignment to remove average bias.

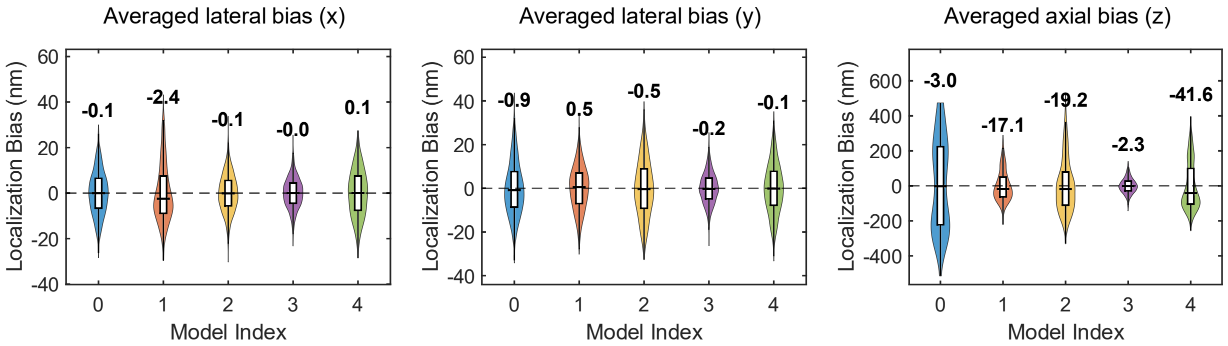

**Supplementary Fig. SS 18. Aligned localization bias induced by another case of grid-dependent aberrations.**

Test data were generated using the PSF model retrieved from a bead in FOV3 (**Supplementary Fig. SS9**), spanning axial positions from –0.7 µm to +0.7 µm in 100 nm steps (100 frames per z-position). Simulations were performed with a signal photon count of $1 \times 10^4$ and a background photon count of 400. Five PSF models were used for 3D localization: Model 0 (theoretical pupil function), Models 1–4 (experimental PSFs from beads in FOV1-4 in **Supplementary Fig. SS9**). Localization bias distributions aggregated over all positions are also shown. Localization biases were aligned by subtracting the average bias across all localizations, highlighting relative deviations.

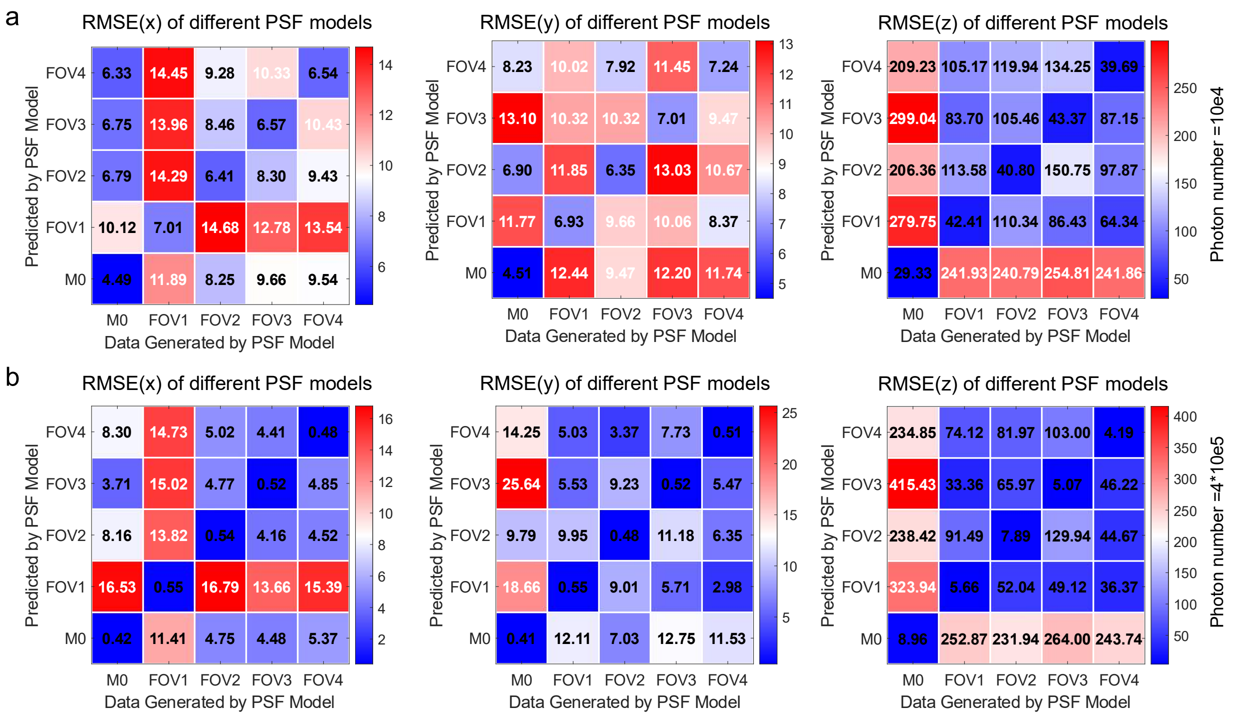

**Supplementary Fig. SS 19. Aligned localization RMSE induced by another case of grid-dependent aberrations.**

Test data were generated using the PSF model retrieved from a bead in FOV3 (**Supplementary Fig. SS9**), spanning axial positions from –0.7 µm to +0.7 µm in 100 nm steps (100 frames per z-position). Simulations were performed with a signal photon count of $1 \times 10^4$ and $4 \times 10^5$ for (a) and (b), and a background photon count of 400 photons per pixel. Five PSF models were used for 3D localization: Model 0 (theoretical pupil function), Models 1–4 (experimental PSFs from beads in FOV1-4 in **Supplementary Fig. SS9**). These PSF models were used for 3D localization, and RMSE in x-, y-, and z-localization was calculated after alignment to remove average bias.

#### 2. Simulation of PSFs Based on Scalar Diffraction Theory

According to scalar diffraction theory[3,4], the PSF can be calculated as the squared magnitude of the inverse Fourier transform of the pupil function:

$$\mu_0(x, y, z) = \left|\mathcal{F}^{-1}\left[h(k_x, k_y)\, e^{i(k_x x + k_y y)}\, e^{i2\pi k_z z}\right]\right|^2,$$

where $\mu_0(x, y, z)$ represents the PSF intensity at position $(x, y, z)$ in the image space, $\mathcal{F}^{-1}$ denotes the inverse Fourier transform, $h(k_x, k_y)$ is the amplitude of the pupil function, $e^{i(k_x x + k_y y)}$ is the lateral phase term, and $e^{i2\pi k_z z}$ accounts for defocus. The axial spatial frequency $k_z$ is defined as:

$$k_z = \left(\left(\frac{NA}{\lambda}\right)^2 - k_x^2 - k_y^2\right)^{1/2},$$

where NA is the numerical aperture of the objective and $\lambda$ is the emission wavelength.

##### 2.1. Modeling Aberrations with Zernike Polynomials

Aberrations were introduced as phase deviations in the pupil plane, modeled as a weighted sum of Zernike polynomials[5]

$$\mu_0(x, y, z; c_1, c_2, \ldots, c_n) = \left|\mathcal{F}^{-1}\left[h(k_x, k_y)\, e^{i(k_x x + k_y y)}\, e^{i2\pi k_z z}\, e^{i\sum_{j=1}^{n} c_j \varphi_{M_j}}\right]\right|^2,$$

where $c_j$ are the Zernike coefficients, $\varphi_{M_j}$ are the corresponding Zernike modes representing specific aberration types, and n is the number of modes considered.

##### 2.2. PSFs with Refractive Index Mismatch

To simulate index mismatch-induced aberrations, an additional phase term $\varphi_\alpha$ was added to the pupil function:

$$\mu_0(x, y, z') = \left|\mathcal{F}^{-1}\left[h(k_x, k_y)\, e^{i(k_x x + k_y y)}\, e^{i\varphi_\alpha}\right]\right|^2,$$
$$\varphi_\alpha = \frac{2\pi}{\lambda}\left[z_0 n_{med}\cos(\theta_{med}) - z_0 \frac{n_{imm}^2}{n_{med}}\cos(\theta_{imm}) + z' n_{med}\cos(\theta_{med})\right],$$

where $z_0 = z_{stage} \cdot n_{med}/n_{imm}$ is the optical path-corrected reference position, with $z_{stage} = 0$ at the coverslip surface. $z'$ is the relative axial position, $n_{med}$ and $n_{imm}$ are the refractive indices of the sample medium and immersion medium, and $\theta_{med}$, $\theta_{imm}$ are the collection angles in each medium.

#### 2.3. Biplane PSF Simulation

For biplane imaging simulation, we introduced an axial separation $Z_d$ between two focal planes and generated a pair of PSFs using a common aberrated pupil function[6]:

$$\mu_{01}(x, y, z; c_1, \dots, c_n) = \left|\mathcal{F}^{-1}\left[h(k_x, k_y)\, e^{i(k_x x + k_y y)}\, e^{i2\pi k_z\left(z - \frac{Z_d}{2}\right)}\, e^{i\sum_{j=1}^{n} c_j \varphi_{M_j}}\right]\right|^2,$$
$$\mu_{02}(x, y, z; c_1, \dots, c_n) = \left|\mathcal{F}^{-1}\left[h(k_x, k_y)\, e^{i(k_x x + k_y y)}\, e^{i2\pi k_z\left(z + \frac{Z_d}{2}\right)}\, e^{i\sum_{j=1}^{n} c_j \varphi_{M_j}}\right]\right|^2,$$

where the defocus terms center the imaging planes symmetrically around the nominal axial position, and $Z_d$ denotes the inter-plane distance.

#### 2.4. Intensity and Background Scaling

After generating normalized PSFs, intensity and background levels were incorporated to simulate experimental imaging conditions:

$$\mu = I \cdot \mu_0 + bg,$$

where $\mu_0$ is the normalized PSF, I represents the total signal intensity (photons), and bg denotes the background photon level. The photon intensity and background levels were estimated from corresponding experimental measurements using 32×32 pixel subregions (**Supplementary Fig. 3**). For all biplane datasets used in the localization bias analysis, the default photon count was set to $1\times10^4$, approximately one-tenth of the measured intensity from 200 nm beads. This downscaling was intentionally applied to better approximate the signal level of single-molecule emitters, which exhibit lower photon budgets than fluorescent beads. Poisson noise was then added to each simulated image to mimic realistic photon shot noise and emulate actual imaging conditions.

##### 2.5. Gaussian Blur for Fluorescent Bead

Large fluorescent beads appear brighter and more stable than single emitters due to their higher fluorophore content. Their images can be modeled as a convolution of the PSF with the fluorophore distribution inside the bead. To simulate this effect, we included a Gaussian blur:

$$\mu_{psf} = \mu \otimes g$$

where $g$ is a Gaussian kernel. The blur parameter (σ) was estimated by comparing simulated PSFs with experimental bead images using NCC. The optimal σ yielding the highest NCC was selected for phase retrieval (see **Supplementary Fig. 2**).

In all experiments, we simulated the following types of PSF models for quantitative evaluation:

1. **Ideal PSF:** without aberrations.
2. **Index mismatch PSF:** incorporating refractive index mismatch.
3. **Aberrated PSF:** with sample- and system-induced aberrations for phase retrieval.
4. **Biplane PSF:** generated using the same phase-retrieved aberrations under a dual-plane configuration for localization analysis.

These simulations enabled systematic assessment of the impact of various aberration sources on localization accuracy under realistic imaging conditions.

#### 3. Localization methods

Due to the ambiguity inherent in 3D PSF-based localization, direct 3D localization of our experimental data is not feasible. To overcome this limitation, we adopted a biplane imaging strategy, which mitigates axial ambiguity by introducing an additional focal plane (as illustrated in **Supplementary Fig.4**), and generated simulated datasets for localization analysis. To closely replicate experimental conditions, optical aberrations were incorporated into the simulated PSFs using phase profiles retrieved from our MLE-based phase retrieval analysis. This approach enables systematic evaluation of localization performance under realistic aberration and noise conditions, while preserving access to the ground truth for quantitative assessment.

##### 3.1. Biplane Distance Determination

In a biplane imaging system, fluorescence emission from single emitters is split into two optical paths, with one plane intentionally defocused relative to the other. This configuration introduces depth-dependent variations in the PSF, enabling precise estimation of the emitter's axial (z) position (**Supplementary Fig. SS20**).

A critical parameter influencing localization precision in biplane systems is the biplane distance which is defined as the axial separation between the two focal planes. To optimize localization accuracy, we determined the optimal biplane distance based on the CRLB. As shown in **Supplementary Fig. SS20**, we computed the CRLB using a fixed PSF model under defined photon and background noise conditions, while systematically varying the biplane distance from 0.1 μm to 2.0 μm in 100 nm increments. Localization precision in x, y, and z was evaluated across emitter positions spanning –1 μm to +1 μm along the axial axis.

Under ideal conditions (i.e., no aberrations) and in room-temperature imaging where aberrations are relatively mild, the optimal biplane distance can be clearly identified as the one that minimizes and balances localization uncertainty across the z-range. However, in cryogenic imaging systems, where optical aberrations are stronger and more complex, the optimal biplane distance becomes highly variable across different axial positions, particularly degrading z-localization precision. To address this challenge, we calculated the average localization precision over the axial range of –0.7 μm to +0.7 μm for each biplane distance. The final biplane distance was then selected to optimize and balance localization precision in x, y, and z dimensions across the entire imaging volume, set to 1 μm in our experimental design.

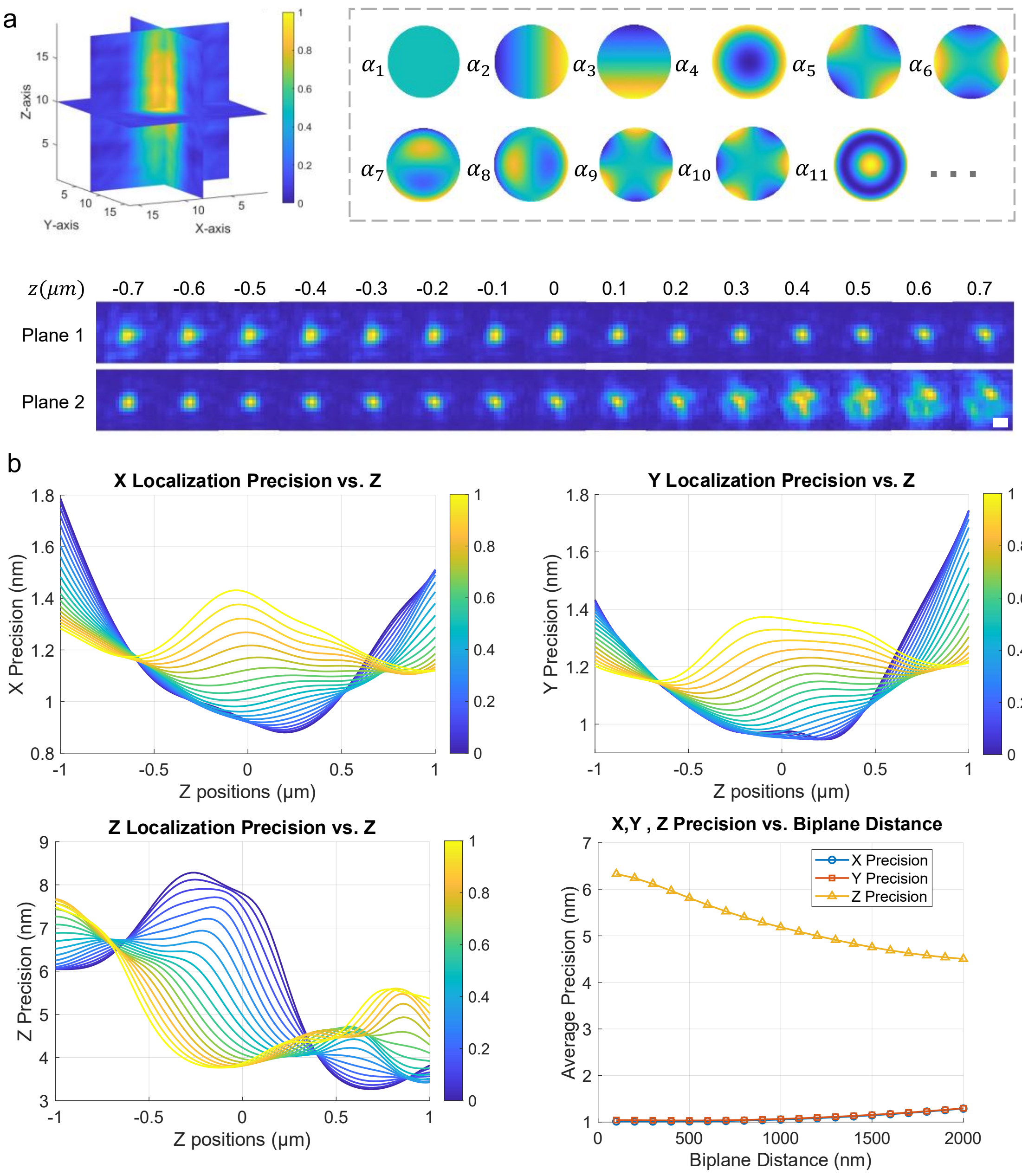

**Supplementary Fig. SS 20. Biplane-based data simulation and optimal biplane distance determination.**

**(a)** Simulation setup using a biplane imaging configuration to resolve axial ambiguity and enable 3D localization. The PSF model is constructed using a Zernike polynomial expansion, where each term corresponds to a specific aberration mode. The simulated dataset consists of two image stacks representing distinct focal planes, mimicking the biplane imaging system. **(b)** Determination of the optimal biplane distance for localization precision. The plots show the Cramér–Rao lower bound (CRLB) for x-, y-, and z-localization across axial positions from −1 μm to +1 μm, under a typical photon count (~1 × $10^5$) with aberrations estimated from representative experimental data. CRLBs are calculated for various biplane distances. The bottom-right panel summarizes the average CRLB within the axial range of −0.7 μm to +0.7 μm as a function of biplane distance for each localization dimension. For a balanced tradeoff between lateral and axial precision, a biplane distance of 1 μm is used in all simulated datasets. Scale bar: 500 nm.

##### 3.2. Initial Guess for 3D Localization

Our 3D localization algorithm, based on cubic spline fitting, estimates five key parameters for each emitter: lateral positions (x, y), axial position (z), photon count, and background photon level. To ensure that the simulated dataset closely mimics the experimental conditions, simulations are generated using the PSF model retrieved from the corresponding experimental data.

For accurate 3D localization, the optimization process is initialized using the ground-truth values of all five parameters: x, y, z, photon number, and background level. This direct initialization significantly enhances localization accuracy, especially under low-photon conditions, when compared to traditional approaches that rely on independent pre-fitting or heuristic estimation. By integrating experimentally derived parameters with an optimized initialization strategy, our simulation framework provides a more realistic and robust benchmark for evaluating 3D localization performance.

##### 3.3. MLE-Based 3D Localization with Channel-Specific PSF and GPU Acceleration

In our 3D localization framework for high-precision single-molecule localization using a biplane imaging strategy, we adopted a channel-specific PSF modeling approach to accurately account for optical distortions between the two detection planes. The implementation was adapted from our previous work[7]. This strategy preserves the statistical integrity of the imaging data—especially under sCMOS detection, where Poisson-distributed photon noise dominates and simplifying assumptions about readout noise are valid.

In this 3D localization algorithm, channel-specific PSF model is directly incorporated into the MLE to jointly estimate seven parameters: emitter position $(x, y, z)$, photon counts in each plane

$(I_1, I_2)$, and background levels $(bg_1, bg_2)$. For each subregion $D = \{D_1, D_2\}$ from the two detection planes, the likelihood function was constructed under a Poisson noise model:

$$L(\theta|D) = \prod_q \frac{(\mu_{1,q})^{D_{1,q}} e^{-\mu_{1,q}}}{D_{1,q}!} \cdot \frac{(\mu'_{2,q})^{D_{2,q}} e^{-\mu'_{2,q}}}{D_{2,q}!},$$

where q indexes the pixels, and $\mu_{1,q}$ and $\mu'_{2,q}$ represent the modeled PSF values in planes 1 and 2, respectively. The parameter vector $\theta = (x, y, z, I_1, I_2, bg_1, bg_2)$ was estimated by minimizing the negative log-likelihood:

$$-\ln L = \sum_q [\mu_{1,q} - D_{1,q}\ln(\mu_{1,q})] + \sum_q [\mu'_{2,q} - D_{2,q}\ln(\mu'_{2,q})].$$

A modified Levenberg–Marquardt algorithm was used to perform the parameter optimization. The first derivative (gradient) of the log-likelihood with respect to θ is:

$$f = \frac{\partial \ln L}{\partial \theta} = \sum_q \left(1 - \frac{D_{1,q}}{\mu_{1,q}}\right)\frac{\partial \mu_{1,q}}{\partial \theta} + \sum_q \left(1 - \frac{D_{2,q}}{\mu'_{2,q}}\right)\frac{\partial \mu'_{2,q}}{\partial \theta},$$

and the second derivative (Hessian approximation) is:

$$f' = \frac{\partial f}{\partial \theta} = \sum_q \frac{D_{1,q}}{\mu_{1,q}^2}\left(\frac{\partial \mu_{1,q}}{\partial \theta}\right)^2 + \sum_q \frac{D_{2,q}}{(\mu'_{2,q})^2}\left(\frac{\partial \mu'_{2,q}}{\partial \theta}\right)^2.$$

Second-order derivatives of the PSF with respect to θ were neglected to reduce computational complexity. Parameter updates followed the standard Levenberg–Marquardt rule:

$$\theta_{n+1} = \theta_n - \frac{f}{f'(1+\beta)},$$

where β is a damping factor that controls convergence speed and was set to zero in our implementation, corresponding to a Gauss–Newton update. The localization speed in this framework is primarily constrained by the computational cost of evaluating 3D PSF models during the fitting process. To alleviate this bottleneck, we precomputed the PSF over a high-resolution 3D grid and employed cubic interpolation to evaluate the PSF at arbitrary positions in real time. This interpolation was implemented on the GPU using CUDA, significantly accelerating the likelihood evaluation step. As a result, our framework achieved a localization throughput of approximately 240 PSFs per second, enabling efficient processing of large-scale biplane imaging datasets.

#### 4. Quantification analysis

##### 4.1. PSF model difference

To quantify differences in aberrations between PSF models, we employed two complementary metrics: (1) the cosine similarity of Zernike polynomial coefficients and (2) the root mean square (RMS) phase error between pupil phase functions. Together, these metrics capture both the nature (type) and severity (magnitude) of aberration differences among FOV-, square-, and grid-dependent PSF models.

Let $c_1$ and $c_2$ denote the vectors of Zernike coefficients (typically the primary terms, e.g., $5^{th}$ to $11^{th}$ in Noll order) for two PSF models. The cosine similarity is defined as:

$$\text{Cosine Similarity} = \frac{\mathbf{c_1} \cdot \mathbf{c_2}}{\|\mathbf{c_1}\|\|\mathbf{c_2}\|} = \frac{\sum_{i=5}^{n} c_{1,i}c_{2,i}}{\sum_{i=5}^{n} c_{1,i}^2 \ \sum_{i=5}^{n} c_{1,i}^2}$$

This metric ranges from –1 (completely dissimilar) to 1 (identical direction), providing a normalized measure of how closely two sets of Zernike coefficients align in direction, independent of magnitude. A higher cosine similarity indicates similar types of aberrations, while lower values suggest more distinct aberration patterns.

To assess the magnitude of aberration differences between two PSF models, we calculated the RMS phase error between their respective pupil phase functions. Let $c_1$ and $c_2$ represent the Zernike coefficient vectors (e.g., $5^{th}$ to $11^{th}$ terms in Noll order) corresponding to two models. The RMS phase error is computed as:

$$\text{RMS Phase Error} = \sqrt{\frac{1}{\mathrm{A}} \int_{\mathrm{P}} [\Phi_1(\rho,\theta) - \Phi_2(\rho,\theta)]^2 \, \mathrm{dA}}$$

where A is the area of the unit pupil. This metric captures the total wavefront deviation between two models and reflects both the spatial distribution and magnitude of aberrations. Unlike coefficient-based comparisons, this phase-space metric provides a more direct and physically interpretable measure of optical error across the aperture.

##### 4.2. Localization Error

To evaluate localization accuracy, we quantified axial localization error across a range of z-positions using a simulated dataset. The ground-truth axial positions ranged from $-0.7\ \mu m$ to $+0.7\ \mu m$ in 100 nm increments, resulting in 15 discrete axial planes. For each position, 100 simulated image frames were generated, yielding a total of 1500 frames for analysis.

The localization bias at each axial position $z_k$ was defined as the median deviation between the predicted axial positions $\hat{z}_i(k)$ and the corresponding ground-truth values $z_i^{GT}(k)$:

$$B(k) = \text{median}(\hat{z}_i(k) - z_i^{GT}(k)), i = 1, \ldots, N$$

where $k = 1,2, \ldots ,15$ indexes the axial positions and N is the number of frames per position.

To summarize overall performance across the full axial range, the average localization bias was computed as:

$$B_{avg} = \frac{1}{K}\sum_{k=1}^{K} B\,(k),$$

where $K = 15$ is the total number of axial planes.

In addition to bias, we computed the RMSE across all frames as a more comprehensive metric that incorporates both bias and variance:

$$B_{RMSE} = \sqrt{\frac{1}{MN}\sum_{k=1}^{M}\sum_{i=1}^{N}\left(\hat{z}_i(k) - z_i^{GT}(k)\right)^2}.$$

This aggregated RMSE measure captures the overall localization accuracy across the 3D volume.

To assess the effectiveness of the localization strategy, we compared the experimental RMSE values to the theoretical CRLB, which represents the fundamental limit of localization precision under specified imaging conditions. This comparison provides insight into how closely the achieved localization performance approaches the theoretical optimum.

##### 4.3. Partial normalized cross correlation

To quantitatively assess the similarity between PSFs while emphasizing aberration-induced distortions, we introduce the partial Normalized Cross-Correlation (Partial NCC, PNCC) metric. Traditional NCC is computed over the entire PSF image stack, including both in-focus and out-of-focus frames. However, the in-focus frames of different PSFs tend to be highly similar, making it difficult to distinguish aberration-induced differences. This often results in artificially high NCC values, masking subtle variations caused by optical distortions.

To overcome this limitation, PNCC is computed by selectively correlating only a subset of out-of-focus frames, where aberrations have a more pronounced impact on PSF shape. Given two PSF image stacks, $I_1(z)$ and $I_2(z)$, defined over axial positions $z \in [-z_{max}, z_{max}]$, the PNCC is expressed as:

$$\text{PNCC}=\max_{\Delta z} \frac{\sum_{z \in Z_{partial}} \sum_{x,y} (I_1(x,y,z) - \bar{I}_1(z))\,(I_2(x,y,z+\Delta z) - \bar{I}_2(z+\Delta z))}{\sqrt{\sum_{z \in Z_{partial}} \sum_{x,y} (I_1(x,y,z) - \bar{I}_1(z))^2}\sqrt{\sum_{z \in Z_{partial}} \sum_{x,y} (I_2(x,y,z+\Delta z) - \bar{I}_2(z+\Delta z))^2}}$$

where $Z_{partial}$ represents the set of selected out-of-focus axial positions, and $\bar{I}_1(z)$ and $\bar{I}_2(z)$ are the mean intensities of the respective image stacks at each axial position. The parameter $\Delta z$ accounts for potential axial misalignment between the two PSFs.

By excluding in-focus frames and focusing on out-of-focus regions, PNCC enhances sensitivity to aberration-induced distortions that might otherwise be overlooked. This approach provides a more accurate assessment of PSF variations and aberrations in cryo-FLM imaging, enabling a more precise evaluation of optical distortions across different imaging conditions.

##### 4.4. CRLB calculation and analytical gradients

To evaluate the optimal localization precision, we compute the Fisher information matrix which quantifies the information content of the single-molecule emission pattern. For single-molecule imaging, assuming the number of photons detected in each pixel are independent random variables following a Poisson distribution and using Stirling's approximation, the Fisher information matrix can be defined as

$$I_{ij}(\theta) = \sum_{k=1}^{K} \frac{1}{\mu_k(\theta)} \frac{\Delta\mu_k(\theta)}{\Delta\theta_i} \frac{\Delta\mu_k(\theta)}{\Delta\theta_j}$$

where $\mu_k(\theta)$ is the value of the PSF model at pixel k, N is the total number of the total number of the pixels in one fitting sub-region and $\theta = [\theta_1, \theta_2, \cdots, \theta_n]^T$ is the parameter vector. According to the Cramér–Rao inequality, the estimation variance of any unbiased estimator $\theta_i$, i.e. CRLB was calculated as

$$\text{var}(\theta_i) \geq [I(\theta)^{-1}]_{ii}$$

where $\text{var}(\theta_i)$ is the estimation variance of an estimator. For biplane setup, by incorporating the noise characteristic (Poisson noise and pixel-independent readout noise) of the sCMOS camera and the channel-specific PSF model, the relevant Fisher information in each element can be calculated as

$$I_{ij}(\theta) = \sum_{M}^{m=1} \sum_{N_m}^{k_m=1} \frac{1}{\mu_{m,k_m}(\theta)} \frac{\Delta\mu_{m,k_m}(\theta)}{\Delta\theta_i} \frac{\Delta\mu_{m,k_m}(\theta)}{\Delta\theta_j}$$

where M is the number of planes, 2 for the biplane system; $N_m$ is the pixel numbers in the m-th plane, in our simulation we used $32 \times 32$ pixels subregions, so $N_m = 1024$; $\mu_{m,k_m}(\theta)$ is the value of the PSF model at pixel k in m-th plane; $\theta$ is the parameter vector to be estimated.

In the context of assessing the impact of aberrations on localization precision, the parameters of interest include $\lfloor\{x, y, z, I, bg\}\rfloor$, where $\lfloor\{x, y, z\}\rfloor$ is the 3D position (unit: nm) of the single molecule, I is the detected photon count, bg is the background photon count per pixel. When calculating the derivative numerically, we set the increments $\Delta x = 0.1$ pixel, $\Delta y = 0.1$ pixel and $\Delta z = 0.01\text{um}$. Consequently, the Fisher information matrix was a $5 \times 5$ matrix. The diagonal terms of the inverse of Fisher information matrix correspond to the CRLB for the respective parameter. The square root of the diagonal terms in the CRLB matrix provided the theoretical achievable estimation precision for each parameter. To remove the influences of photon numbers emitted by a single molecule which improves the estimation variance with a reciprocal dependence, we quantified the precision per photon count without background noise.

#### 5. Protocol for Analyzing Aberrations in Cryo-FLM Imaging Systems

This protocol is designed to highlight the impact of aberration-induced localization errors in cryo-FLM and to provide a practical framework for analyzing such aberrations in both system-specific and sample-dependent contexts. Our goal is to enable users to perform aberration analysis tailored to their own imaging setups and sample types.

The protocol consists of three major components: (1) **Sample Preparation** – for generating well-controlled fluorescent bead samples suitable for aberration characterization; (2) **Imaging** – including room-temperature and cryo-FLM acquisition using standardized settings and procedures; (3) **Analysis** – covering phase retrieval, aberration quantification, and optional simulation-based evaluation of localization errors.

By following this workflow, users can systematically evaluate the sources and effects of optical aberrations in cryo-FLM and implement corrections to improve localization accuracy.

We suggest the use of 200-nm TetraSpeck fluorescent beads to maximize the signal per PSF and allow potential future evaluation of chromatic aberrations; however, any fluorescent bead smaller than the PSF radial diameter will work.

##### 5.1. Protocol: Preparation of Fluorescent Bead Samples for Cryo-FLM

**Objective:**

Prepare vitrified fluorescent bead samples for cryo-FLM, ensuring optimal dispersion and vitrification.

**Materials:**

200 nm TetraSpeck fluorescent beads (Thermo Fisher Scientific)
1× Phosphate-Buffered Saline
Holey carbon-coated TEM grids (e.g., C-Flat 2/2-300)
Glow discharge unit
Vitrobot or other plunge-freezer
Liquid ethane and nitrogen
Cryo-grid box and forceps
Filter paper, aluminum foil

**Procedure:**

Bead Dilution:

- Dilute bead stock 1:50 in PBS.
- Sonicate in a waterbath sonicator for 5 min and briefly vortex to break up clumps
- Protect from light using foil.

Grid Preparation:

- Glow-discharge TEM grids to improve hydrophilicity according to instrument protocols
- Apply 3 µL of bead suspension onto grid.
- Blot using optimized conditions (e.g., 4 s, force –3).
- Plunge-freeze in liquid ethane.

Storage:

- Transfer grids to cryo-grid box under liquid nitrogen.
- Store in liquid nitrogen-cooled Dewar at –196 °C until imaging.

##### 5.2. Protocol: Preparation of Fluorescent Bead Samples for Room-Temperature and Cryo-FLM Imaging

**Objective:**

To prepare a single fluorescent bead sample suitable for both room-temperature and cryo-FLM, while preserving spatial bead distribution across conditions.

**Materials**

200 nm TetraSpeck fluorescent beads (Thermo Fisher Scientific)

1× PBS

Holey carbon-coated London Finder TEM grids

Glow discharge unit

Filter paper

Aluminum foil

Coverslip

Deionized (DI) water

Vitrobot

Liquid ethane and nitrogen

Cryo-grid box and forceps

Cryo storage dewar

**Procedure:**

Bead Dilution and Dispersion:

- Dilute TetraSpeck stock 1:50 in PBS.
- Sonicate for 5 min and vortex briefly.
- Wrap in aluminum foil to protect from light.

Grid Preparation:

- (Optional) Coat EM grids with ~12 nm amorphous carbon for enhanced mechanical stability.
- Glow-discharge the grids to increase surface hydrophilicity.

Room-Temperature Sample Preparation:

- Apply 3 μL of diluted bead suspension to a Finder EM grid.
- Allow to settle for ~5 seconds.
- Gently blot with filter paper for even distribution.
- Dry overnight in the dark on a coverslip.

Conversion to Cryo-FLM Sample:

- Rehydrate dried grid with 3 μL DI water.
- Use Vitrobot to vitrify the sample (e.g., 4 s blot time, –3 blot force, 100% humidity).
- Store in cryo-grid box submerged in liquid nitrogen until imaging.

##### 5.3. Protocol: Sample Mounting for Cryo-FLM Imaging

Follow instrument protocols to load grid onto microscope. Perform this procedure after the cryo-FLM system has reached operating temperature.

##### 5.4. Protocol: Cryo-FLM Imaging Using commercial Leica LAS X

**Objective:**

Perform cryogenic fluorescence imaging of vitrified samples using the Leica LAS X system.

**A. Software Initialization & Imaging Setup**

Launch LAS X in widefield mode and open the Assay Editor.

Configure imaging in Acquire → Acquisition:

- Select Brightfield and GFP channels.

- Adjust filters, illumination, exposure, and acquisition mode (Z-stack or single frame).
- Save these settings for reuse.

**B. Mosaic Acquisition (Optional for Grid Mapping)**

Create a Carrier Model in the Assay Editor to define grid geometry (3 mm × 3 mm chambers, 2×1 layout).

Use Spiral mode to scan the grid in the Brightfield channel.

Align software tiles to the physical EM grid using STP controls.

Finalize coverage and merge Brightfield and GFP mosaics as needed.

**C. Multi-frame Z-Stack Imaging**

From the GFP mosaic, select ROIs with isolated fluorophores.

Set Z-stack parameters (adjust as needed based on your imaging system and experimental requirements):

- Z-range: 4 µm
- Step size: 200 nm

Enable Time acquisition to capture multiple frames at each Z-slice.

Toggle between Z and T modes to configure acquisition:

- Pre-calculate Z positions in the Z-stack tab.
- Define frame number in the Time tab.

Repeat for all ROIs.

Save the project and export data (e.g., TIFF format).

**D. Sample Unloading & System Shutdown**

Attach cooled transfer shuttle and retrieve the cartridge.

Return the grid to storage in liquid nitrogen.

To shut down:

- Raise objective and start stage bake-out.
- After 30 min, remove and dry the pump.
- Power down all system components.
- Empty and dry the LN dewar.

#### 5.5. Protocol: Room-temperature FLM Imaging Using Leica LAS X

The imaging procedure at room temperature closely follows that used under cryogenic conditions, with two key differences: the LN pump is not activated during imaging, and liquid nitrogen cooling is not required during sample transfer. The entire workflow is conducted under ambient temperature conditions.

#### 5.6. Protocol: Phase Retrieval Using MATLAB Toolbox

We provide a MATLAB-based phase retrieval toolbox to extract aberrations from bead image stacks. This process enables recovery of the pupil function and Zernike polynomial coefficients that describe system-specific and sample-induced aberrations.

**Input Requirements:**

- A 3D z-stack of a single bead, e.g. acquired from –1.4 μm to +1.4 μm in axial position, using a 200 nm step size.
- You may adjust the z-range and step size based on your imaging setup and dataset.

**System Parameters to Specify:**

- Pixel size (in microns)
- Numerical aperture (NA) of the objective
- Emission wavelength (in nanometers)
- Refractive index of the sample medium (typically 1.33 for vitrified PBS)
- Refractive index of the objective immersion medium (typically 1.00 for cryogenic systems using liquid nitrogen)
- Camera offset and gain

**Noise Considerations and Imaging Recommendations:**

- Due to the strong noise inherent in cryo-FLM imaging, we recommend:
  - Using high laser intensities to boost signal
  - Enabling low-noise acquisition mode on the camera
- Noise may impact the robustness of the phase retrieval algorithm. Therefore, we recommend repeating the retrieval multiple times and selecting the best result based on evaluation metrics.

**Output Files:**

- A .mat file containing:
  - Predefined system parameters
  - Phase-retrieved Zernike coefficients
  - Reconstructed PSF and pupil function
  - Evaluation metrics including:
    - Normalized cross-correlation (NCC)
    - Maximum likelihood estimation (MLE) loss

**Note:** Please ensure that the bead used for phase retrieval is well-isolated and centrally located in the field of view to minimize edge effects and improve reconstruction fidelity.

##### 5.7. Optional: Simulation-Based Evaluation of Aberration-Induced Localization Error

This step is optional. Based on the phase retrieval results, one can qualitatively and quantitatively assess the presence and magnitude of aberrations across different regions of the dataset. However, if the impact of these aberrations on localization accuracy is unclear, we provide a MATLAB-based simulation framework to evaluate the expected localization error introduced by specific aberration profiles.

We offer two dedicated MATLAB toolboxes:

1. **Biplane Data Simulation Toolbox:** This toolbox generates synthetic biplane datasets using user-defined imaging parameters, including:
   - Photon counts
   - Background levels
   - 3D PSF models with or without aberrations
2. **Biplane 3D Localization Toolbox:** This toolbox performs 3D localization using cubic spline fitting of the PSF model (including aberrations). It supports:
   - Accurate emitter localization in x, y, and z
   - Support for different noise levels and PSF configurations

#### Reference

1. Liu, S. *et al.* Universal inverse modeling of point spread functions for SMLM localization and microscope characterization. *Nat. Methods 2024 21:6* **21**, 1082–1093 (2024).
2. Jouchet, P., Roy, A. R. & Moerner, W. E. Combining deep learning approaches and point spread function engineering for simultaneous 3D position and 3D orientation measurements of fluorescent single molecules. *Opt. Commun.* **542**, 129589 (2023).
3. Born, M. *et al.* Principles of Optics: Electromagnetic Theory of Propagation, Interference and Diffraction of Light. *Principles of Optics* (1999) doi:10.1017/CBO9781139644181.
4. Goodman, J. Introduction to Fourier optics. (2005).
5. Noll, R. J. Zernike polynomials and atmospheric turbulence*. *JOSA, Vol. 66, Issue 3, pp. 207-211* **66**, 207–211 (1976).
6. Juette, M. F. *et al.* Three-dimensional sub–100 nm resolution fluorescence microscopy of thick samples. *Nat. Methods 2008 5:6* **5**, 527–529 (2008).
7. Xu, F. *et al.* Three-dimensional nanoscopy of whole cells and tissues with in situ point spread function retrieval. *Nat. Methods 2020 17:5* **17**, 531–540 (2020).
